# Molecular profiling of sponge deflation reveals an ancient relaxant-inflammatory response

**DOI:** 10.1101/2023.08.02.551666

**Authors:** Fabian Ruperti, Isabelle Becher, Anniek Stokkermans, Ling Wang, Nick Marschlich, Clement Potel, Emanuel Maus, Frank Stein, Bernhard Drotleff, Klaske Schippers, Michael Nickel, Robert Prevedel, Jacob M Musser, Mikhail M Savitski, Detlev Arendt

## Abstract

A hallmark of animals is the coordination of whole-body movement. Neurons and muscles are central to this, yet coordinated movements also exist in sponges that lack these cell types. Sponges are sessile animals with a complex canal system for filter-feeding. They undergo whole-body movements resembling “contractions” that lead to canal closure and water expulsion. Here, we combine 3D optical coherence microscopy, pharmacology, and functional proteomics to elucidate anatomy, molecular physiology, and control of these movements. We find them driven by the relaxation of actomyosin stress fibers in epithelial canal cells, which leads to whole-body deflation via collapse of the incurrent and expansion of the excurrent system, controlled by an Akt/NO/PKG/A pathway. A concomitant increase in reactive oxygen species and secretion of proteinases and cytokines indicate an inflammation-like state reminiscent of vascular endothelial cells experiencing oscillatory shear stress. This suggests an ancient relaxant-inflammatory response of perturbed fluid-carrying systems in animals.

**Highlights:**

- Sponge deflation is driven by tension release in actomyosin stress fibers of epithelial pinacocytes
- Akt kinase/Nitric oxide/Protein kinase G/A regulate actomyosin relaxation
- Agitation-induced deflation coincides with an inflammatory state
- The sponge relaxant-inflammatory response is evolutionary related to similar responses in the vertebrate vascular system

## Introduction

The evolution of multicellularity in animals triggered the emergence of spatially and temporally coordinated responses across different cells and tissues driving shape changes and body movements (1). In most animals, whole-body reactions are orchestrated by the nervous system and implemented by musculature and glands. Sponges (Porifera) are an early-branching lineage of animals that lack neurons and muscle cells (**Figure 1A**), yet display simple forms of coordinated physiology and movement, offering a unique window into how simple animals regulate multicellular responses.

**Figure 1.**
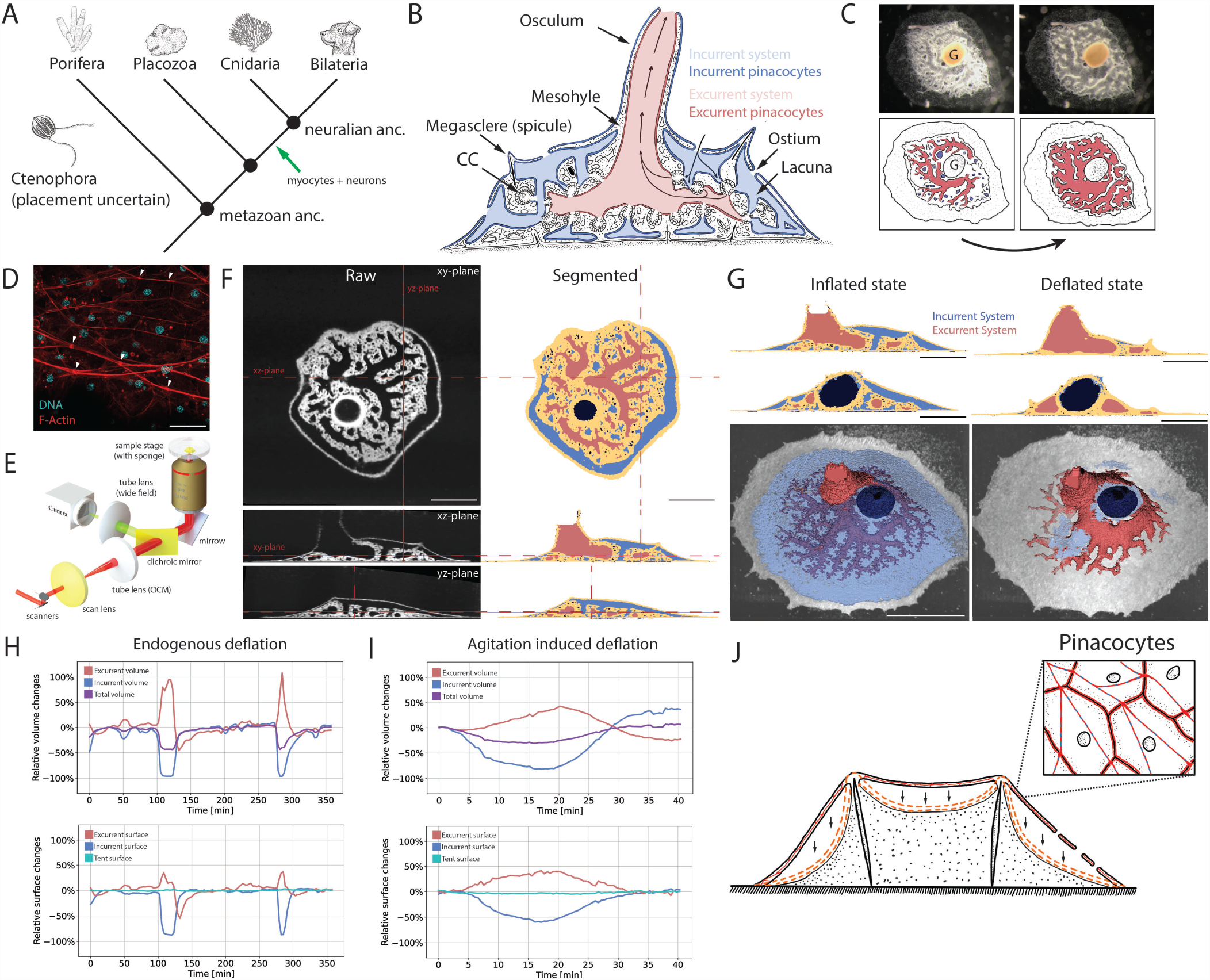
3D in vivo optical coherence microscopy (OCM) of juvenile *Spongilla*. **(A)** Animal phylogeny marking proposed origin of neurons and myocytes. **(B)** Illustration of juvenile *Spongilla lacustris* specimen showing incurrent (blue) and excurrent (red) canal systems. Epithelial pinacocytes are highlighted by darker color tones. CC: Choanocyte chamber. **(C)** Photograph and illustration of endogenous deflation of *S. lacustris* highlighting incurrent (blue) and excurrent (red) systems. G: Gemmule. **(D)** Confocal max intensity projection of actomyosin stress fibers of the tent pinacocytes stained for F-actin (phalloidin, red) and DNA (DAPI, cyan). White arrowheads depict the cell-cell junctions connecting neighboring stress fibers. Scale bar, 30 um. **(E)** Schematic of the microscope body part of the optical coherence microscope (OCM). **(F)** OCM volumes showing orthogonal views of *S. lacustris* in inflated state. Segmented incurrent (blue), excurrent (red) systems, mesohyl/tent (yellow) and gemmule (dark blue). Scale bar, 500 μm. See also Table S1. **(G)** 3D reconstruction of inflated and deflated states. 2D Scale bar, 500 μm. 3D Scale bar, 1000 μm. See also Figure S1. **(H)** Volume and surface area changes measured from segmented OCM volumes during two cycles of endogenous sponge whole-body deflations over 6 hours (f = 1/3 min-1). Excurrent system (red), incurrent system (blue), total sponge (purple), tent (turquoise). **(I)** Relative volume and surface area changes measured from segmented OCM volumes during one cycle of whole-body deflation induced by mechanical agitation (f = 1/30 s-1). **(J)** Illustration of epithelial tent collapse and tent pinacocytes highlighting cell traversing actomyosin stress fibers.

Sponges are composed of an aquiferous canal system lined by epithelial cells called pinacocytes and a collagenous mesenchyme termed mesohyl (**Figure 1B**) (2). In the freshwater sponge *Spongilla lacustris* the canal system comprises incurrent and excurrent canals, interconnected via spherical pumping, filtering and digestive chambers (3). These are lined by the choanocytes with a motile cilium propelling the water flow and a microvillar collar for food capture and ingestion. Water initially enters through openings (ostia) in an outer tent-like epithelial cover into incurrent lacunae and canals. It then passes through the choanocyte chambers and into the excurrent canals before exiting via a chimney-like osculum (**Figure 1B**).

The sessile sponges close and flush their canal system via a spatially and temporally coordinated whole-body movement, previously termed “contraction” or “sneezing” (**Figure 1C**) (4–6), which is thought to remove obstructions in their canal system or regulate gas and nutrient exchange (7). In freshwater sponges this behavior occurs endogenously or upon physical agitation, often as a peristaltic-like wave that initiates at the sponge periphery and spreads centrally in two stages (8– 13). First, closure of ostia, collapse of the tent and incurrent canals, as well as expansion of the excurrent canals can be observed. From a lateral view, the collapse of incurrent lacunae gives the impression of a *deflation* of the sponge body (8, 13), prompting us to relate to the movement as “sponge deflation”, independent of the underlying cellular behavior (see also **Supplemental Note**). In the second phase, the sponge inflates by reopening the ostia and returning incurrent canals and excurrent canals to their original state.

In recent years, various studies have investigated the cellular and molecular underpinnings of sponge deflation (4, 7– 9, 12, 14–17). Current models propose cellular contraction of incurrent pinacocytes (8, 9, 12). Supporting this, pinacocytes have been shown to express genes involved in smooth and non-muscle actomyosin contractility (13, 18) and contain actin stress fibers that traverse the cytoplasm (**Figure 1D, J**), connecting neighboring cells via mixed focal adhesion/adherens-like junctions (8, 19). Treatment with selected pharmacological compounds has indicated involvement of myosin light chain kinase (MYLK), cGMP, and nitric oxide (NO) signaling (13, 18); and a variety of other paracrine signals and second messengers have been shown to initiate, inhibit, or alter these deflations in other sponges (8–11, 13, 20). Yet, the exact cellular processes and physiological changes driving and accompanying these processes have largely remained obscure.

Here, we report a detailed characterization of coordinated whole-body deflations in the freshwater sponge *Spongilla lacustris* - taking advantage of live 3D imaging with a custom optical coherence microscope, pharmacological assays, and functional proteomics applied for the first time in sponges. We show that contrary to current models, the deflation of the sponge body does not involve cellular-level actomyosin contraction. Instead, it is driven by the simultaneous isometric relaxation of both incurrent and excurrent pinacocytes from a default tense state. Functional profiling with quantitative phosphoproteomics (21), thermal proteome profiling (TPP) (22, 23), and secretomics identify a pathway involving Akt kinase, nitric oxide and protein kinase G and A that regulate actomyosin stress fiber tension. Concomitant with deflation, sponges activate an inflammatory state highly reminiscent of the response of the vertebrate vascular system to oscillatory shear stress. Our study indicates evolutionary conservation of an ancient relaxant-inflammatory module in perturbed fluid-carrying epithelial systems that was active in the last common ancestor of sponges and other animals.

## Results

### Optical coherence microscopy (OCM) reveals unchanged or expanding epithelial surfaces during sponge deflation

Wholemount in vivo 3D-imaging of mesoscale animals at cellular resolution poses numerous challenges due to their size, the lack of specific (fluorescence) labeling or radiation damage (e.g. in X-ray based approaches). We reasoned that near-infrared Optical Coherence Microscopy (OCM) would provide strong, label-free contrast between the tissue and water-filled cavities, while ensuring overall low phototoxicity, a large field of view and fast imaging speed. We therefore customized an OCM system for three-dimensional morphological imaging of whole juvenile *S. lacustris* during deflation (**Figure 1E**) (24). To facilitate quantitative 3D image analysis we developed a self-supervised denoising approach that suppressed dominant OCM noise sources (**Figure S1A, B**) for subsequent high-quality segmentation of the *S. lacustris’* aquiferous system (**Figure S1C**). Together, this enabled non-invasive volume imaging of whole juvenile *Spongilla* specimens up to 2-3 mm size at an isotropic optical resolution of ∼2.5 μm and at a time resolution of up to 1 Volume / s, which is unprecedented for sponges and thus allow a unique insight into their internal anatomy before and during deflation.

OCM imaging and segmentation of 8-day old juvenile, inflated sponges (n=5) revealed an intricate connection between vertical incurrent canals and the horizontal excurrent system, which together represent approximately 49 ± 9% of the total sponge volume (**Figure 1F, Table S1, Figure S1D, Video S1**). The epithelial tent is pitched across the whole sponge surface using megascleres as “tent poles” and covers the incurrent lacunae (**Figure 1F, G and J, Video S1**). 3D reconstruction of the excurrent canal system revealed a self-similar, fractal organization maximizing surface area and providing equal volumes for the outflow from individual choanocyte chambers (**Figure 1G**).

We next performed overnight time-lapse OCM imaging to capture endogenous deflations. During these events we observed an overall reduction in sponge volume of approximately 40%, fully consistent with a deflation. Closure of ostia and collapse of the tent and incurrent canals resulted inear total loss (∼96%) of incurrent volume, with simultaneous 108% volume and 36% surface area increase in the excurrent system (Video S2 and S3; **Figure 1G, H; Figure S1E**). Subsequently, sponges returned to their inflated state by restricting excurrent canals and reopening the incurrent system, including a short “overshoot” period before canals returned to their original volume. Strikingly, the tent surface area remained more or less constant, and we did not observe a reduction in area as would be expected if the tent was contracting (**Figure 1H**). In contrast, mesenchymal tissue became condensed, contributing to the reduction in overall sponge volume, supporting previous observations that water completely evacuates intercellular spaces in the mesenchyme during deflation (25). Similar volume and surface area changes were measured when we induced deflation through oscillatory circular agitation on a plate shaker (5 min, 500 rpm, 1.25 cm/s) (**Figure 1I**) or through treatment with the NO donor NOC-12 (**Figure S1F**) (13).

### An Akt kinase/NO/PKG/PKA pathway triggers tension release in epithelial stress fibers

To explore the molecular nature of cellular shape changes driving sponge deflation we next investigated pathways regulating actomyosin contractility in the sponge pinacocytes. We identified a near-complete set of genes regulating smooth and non-muscle actomyosin contractility in *Spongilla*, congruent with surveys in other sponge species (26, 27). This included orthologs of smooth muscle myosin heavy chain (MYH9/10/11/14) and light chain (essential and regulatory MYL), and their direct regulators myosin light chain kinase (MYLK) and phosphatase (MLCP), which alter contractile state via phosphorylation and dephosphorylation of MYL (**Figure 2A, B; Figure S2A**). Confocal imaging of phalloidin-stained sponges consistently identified prominent epithelial stress fibers as major phenotypic manifestations of actomyosin bundles (**Figure 1D, J**) (12, 19). Phylogenetic analysis of the MYLK kinase domain revealed the existence of a MYLK/SPEG/TTN/Twitchin kinase ortholog in *Spongilla*, as well as an ortholog of the related STK17A/B kinase (**Figure S2B**). *Spongilla* also possesses an ortholog of striated muscle myosin heavy chain (MYH1-8/13/15), although other genes specific to vertebrate striated muscles such as the troponin complex and z-disk proteins are absent. Taking advantage of our recent *Spongilla* cell expression atlas (13), expression of striated *Myh, Mylk*, and the *Pp1a* subunit of *Mylp* was shown to be differentially upregulated in pinacocytes, whereas other actomyosin components are more broadly expressed (**Figure S2A**).

**Figure 2.**
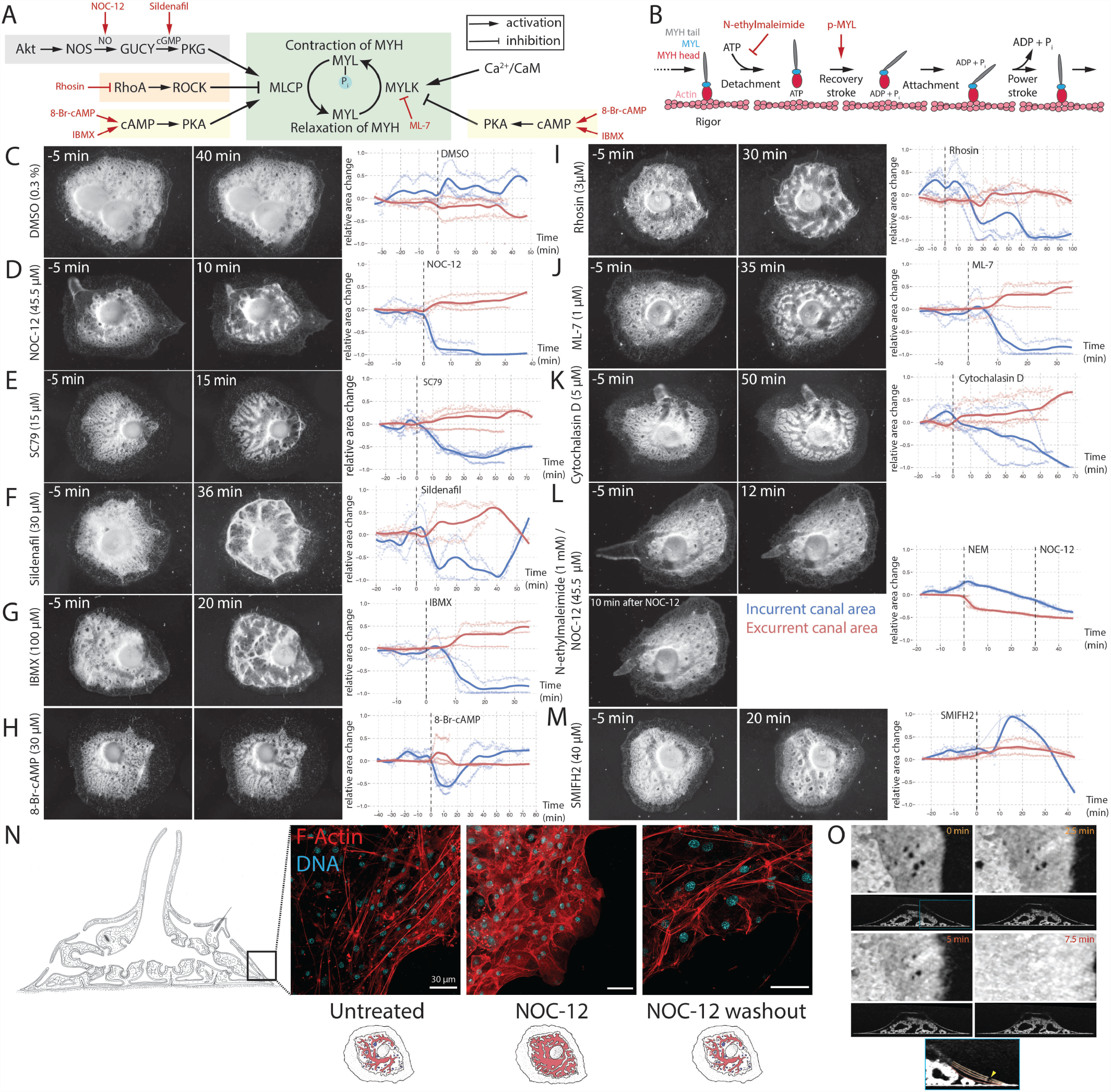
Pharmacological perturbation of actomyosin activity. **(A)** Smooth/non-muscle actomyosin regulatory pathways with pharmacological compounds highlighted in red. MYL: myosin light chain, MYH: myosin heavy chain, MYLK: myosin light chain kinase, MYLP: myosin light chain phosphatase, NOS: nitric oxide synthase, NO: nitric oxide, GUCY: guanylate cyclase, PKG: protein kinase G, ROCK: RhoA/Rho-associated protein kinase, PKA: protein kinase A, CaM: calmodulin. Background colors relate to Figure S2A. See also Figure S2B. **(B)** Actomyosin cross-bridge cycle showing states of myosin conformation and actin binding. N-ethylmaleimide (NEM) treatment blocks ATP binding and myosin release, stabilizing a rigor state. **(C-M)** Time-lapse imaging (f = 1/30 s-1) of *S. lacustris* during the treatment with pharmacological compounds. Overhead images show sponges 5 min pre-treatment (left) and during respective behavior (right). Plots show average change in relative areas across replicates of segmented incurrent (dark blue) and excurrent (dark red) canal areas. See also Figure S2C-N. Dashed line marks application of compound. Individual replicates shown in light tones. **(N)** Confocal max intensity projections of actomyosin stress fibers of the tent pinacocytes stained for F-actin (red, phalloidin) and DNA (cyan, DAPI) in an untreated sponge, NOC-12 treated sponge (deflated) and sponge after NOC-12 washout and return to inflated state. Scale bars, 30 um. See also Figure S2O. **(O)** Ostia closure during agitation induced deflation. Simultaneous tent collapse and ostia closure is visible in the overlay section (blue square). Annotated tent colors correspond to time points in upper panels. Ostium: yellow arrow head. See also Figure S2.

We also identified orthologs for members of key signaling pathways that regulate MYLK and MLCP (**Figure 2A**). Calmodulin, a calcium-dependent activator of MYLK, and Protein kinase A (PKA), which inhibits MYLK and activates MLCP, are broadly expressed across *Spongilla* cell types (13). Other pathways exhibited also specific upregulation of expression in pinacocytes, including key members of the Akt/NO/Protein kinase G (PKG) pathway, nitric oxide synthase (NOS1/2/3) and the nitric oxide receptor guanylate cyclase (GUCY1B3), known to regulate relaxation of vascular endothelial cells in vertebrate blood vessels (**Figure 2A**) (28). Lastly, we found orthologs of RhoA and ROCK1/2 enriched in pinacocytes, whose activation in other systems leads to inhibition of MLCP causing relaxation of smooth/non-muscle actomyosin fibers (29).

To gain insight into the function of these molecules and pathways, we perturbed the activity of actomyosin machinery and of the upstream regulatory pathways using a diverse set of pharmacological compounds (red in **Figure 2A, B**) and assessed sponge responses using 2D time-lapse imaging (**Figure 2C-M; Figures S2C and S2D-N**). Treatment with NOC-12 induced deflation within 5 min after administration (**Figure 2D,** and see above). Treating *Spongilla* with the Akt kinase activator SC79 (30) likewise caused deflation, confirming the involvement of Akt activity (**Figure 2E**). Akt kinase is one of the main activators of NOS in endothelial cells (31, 32). Conversely, NO-induced Akt kinase activation has been reported (33) and offers a straightforward explanation for the self-propagation through positive feedback of a deflation wave spreading across the sponge body despite the short diffusion potential of NO. Consistent with this, the average NO diffusion rate (∼44 μm/s) (34) is similar to the measured wave propagation in *Ephydatia muelleri* (∼0.5 - 70 μm/s (8)). Sildenafil as well as IBMX, both phosphodiesterase inhibitors that increase the concentration of cGMP and cAMP, leading to activation of PKG and PKA (35, 36), induced deflation within 5-10 min (**Figure 2F, G**). Direct usage of cell-permeable 8-Br-cAMP showed the same effect (**Figure 2H**). These results indicated that PKG, as well as PKA, both belonging to the AGC family of protein kinases (37), trigger deflation. Inhibiting the RhoA/ROCK pathway with the selective RhoA inhibitor Rhosin induced deflation (**Figure 2I**), suggesting an additional layer of regulation (38). Direct inhibition of MYLK with ML-7 (39) consistently induced deflation within 20 min after application (**Figure 2J**). Likewise, treatment of the freshwater sponge *Ephydatia muelleri* with the MYH inhibitor para-Aminoblebbistatin had been shown to induce deflation within two hours of application (12).

These observations strongly suggested that *Spongilla* deflation is initiated by a reduction of actomyosin crossbridge cycling through MLCP activation which equals actomyosin relaxation - either via activation of Akt kinase/NO/PKG or PKA or via inhibition of RhoA/ROCK, and subsequent dephosphorylation of the MYL. Corroborating this further, cutting locally into the epithelial tent of the sponge periphery via laser-microdissection led to immediate recoil of the tissue towards the center of the sponge, affirming that the sponge tent is under constant tension in the inflated state (Video S4). Strengthening this new paradigm, we treated sponges with Cytochalasin D, an actin depolymerizing agent. In these cases, sponges consistently but more gradually went into the deflated state, providing further evidence for a tension reduction/relaxation of actin fibers as the driver for the deflation (**Figure 2K**). Finally, confocal imaging of the tent pinacocyte actin bundles revealed that NOC-12 (NO donor) treatment leads to a degradation of the actin fibers (**Figure 2N**), making a clear case that they are dispensable for the deflation and strengthening their role as primarily supportive structures in the inflated state. Removal of NO via the washout of NOC-12 leads to a return of the sponge to the inflated state, coinciding with the repolymerization of the pinacocyte stress fibers within about 15 min (**Figure 2N, Figure S2D**). NO treatment is known to reverse stress fiber formation, e.g. in human umbilical vein endothelial cells (40) or platelets mediated by RhoA inhibition (41).

Our pharmacological data implies isometric tension release in tent pinacocytes (42) as shown in non-muscle cells such as fibroblasts (43), impacting mechanical stability without considerable length changes (44), consistent with our OCM observations of a constant tent surface area. For the epithelial lining of the excurrent system (showing expanded surface, see above) our data is consistent with pinacocytes able to stretch due to stress fiber tension release. Furthermore, the tense state of the pinacocyte actin fibers through continuous crossbridge cycling is consistent with the detection of phosphorylated MYL in the actin bundles of an inflated sponge via a specific antibody (12) (**Figure S2O**).

Additional evidence corroborated the importance of stress fibers for keeping the pinacocyte tent pitched in the inflated state. N-ethylmaleimide (NEM) is an inhibitor of actomyosin tension release, as it irreversibly inhibits ATP binding to the myosin II ATPase domain and thus prohibits the detachment of myosin heads from the actin filaments, which freezes crossbridge cycling (“rigor mortis”) (**Figure 2B**) (45). Treatment of the sponge with NEM accordingly “froze” the tent and entire canal system in its tense state and completely abolished deflation even after NO treatment (**Figure 2L**). Additionally, NEM treatment led to an immediate cessation of cell crawling throughout the sponge. Finally, we treated the sponge with the pan-formin inhibitor SMIFH2 (46). Formins control actin fiber assembly, degradation and nucleation (47) and have been implicated in stress fiber formation (44). After treatment with SMIFH2, the sponges showed an immediate widening of the incurrent canals (**Figure 2M**), in striking contrast to normal deflation, indicating likely increased tension in the actin stress fibers of the epithelial tent, similar to inhibition of the formin DIAPH1 in other studies (48). Taken together, our pharmacological agonist and antagonist treatments make a strong case that actin stress fibers are key to withholding a tensional state in the pinacocyte tent and canals in the inflated state, and that this state is relaxed during deflation.

### Active pumping continues during actomyosin relaxation

To investigate the role of water flow during *Spongilla* inflation, we applied non-digestible ink particles followed by time-lapse imaging. Previous studies had reported that during deflation the sponge continues to pump water through the canal system (49), although reduces exhalant jet speed from the osculum (20). We reproduced these results, observing a pronounced and continuous expansion of the osculum during deflation (**Figure S3A**). These observations can only be explained by unceased pumping of the choanocyte chambers, even in the fully deflated state. The resulting hydrodynamic forces and gauge pressure differences (50) offer an elegant explanation of all observed phenotypes under full relaxation, i.e. total tent and incurrent canal collapse, excurrent canal expansion and osculum widening. The collapse of the tent also involves simultaneous (not preceding) ostia closure (**Figure 2O**), comparable to the actin depolymerization-induced closure of plant stomata (51–53) or relaxation induced chromatophore closure in cuttlefish skin (54). Similarly, pre-tensile forces in the tent may be responsible for an open ostia state. The closure of ostia combined with continuous choanocyte pumping creates a low pressure situation in the incurrent system leading to its further collapse while water is actively pumped into the relaxed excurrent canal system (**Figure S3B**). Due to the tight spatial interconnectedness of both systems and relatively high pressure in the excurrent system (55), this allows for the concurrent expansion of the excurrent canals and eventual water release once full expansion is reached (**Figure S3A**).

### Quantitative Phosphoproteomics and Thermal Proteome Profiling of sponge deflation identify Akt kinase/NO/PKG/PKA, MAPK/ERK, and cell-cell junctions as important players

Since our pharmacological assays indicated tension that release in pinacocytes was broadly regulated by diverse pathways we set out to further investigate the molecular mechanisms and physiological reactions underlying sponge deflation with functional proteomics. We used quantitative phosphoproteomics (56–58) and Thermal Proteome Profiling (TPP) (22, 23, 59–62) to detect post-translational, functional changes between sponges in inflated and deflated state (**Figure 3A**). Quantitative phosphoproteomics identified significantly altered, site-specific phosphorylation states of proteins. TPP measured proteome-wide changes in protein abundance and thermal stability, which has been shown to reflect protein state changes such as binding or dissociation of small molecules, the formation or loss of protein-protein interactions, a change in post-translational modifications, or differences in subcellular localization (22, 23, 59, 60, 62).

**Figure 3.**
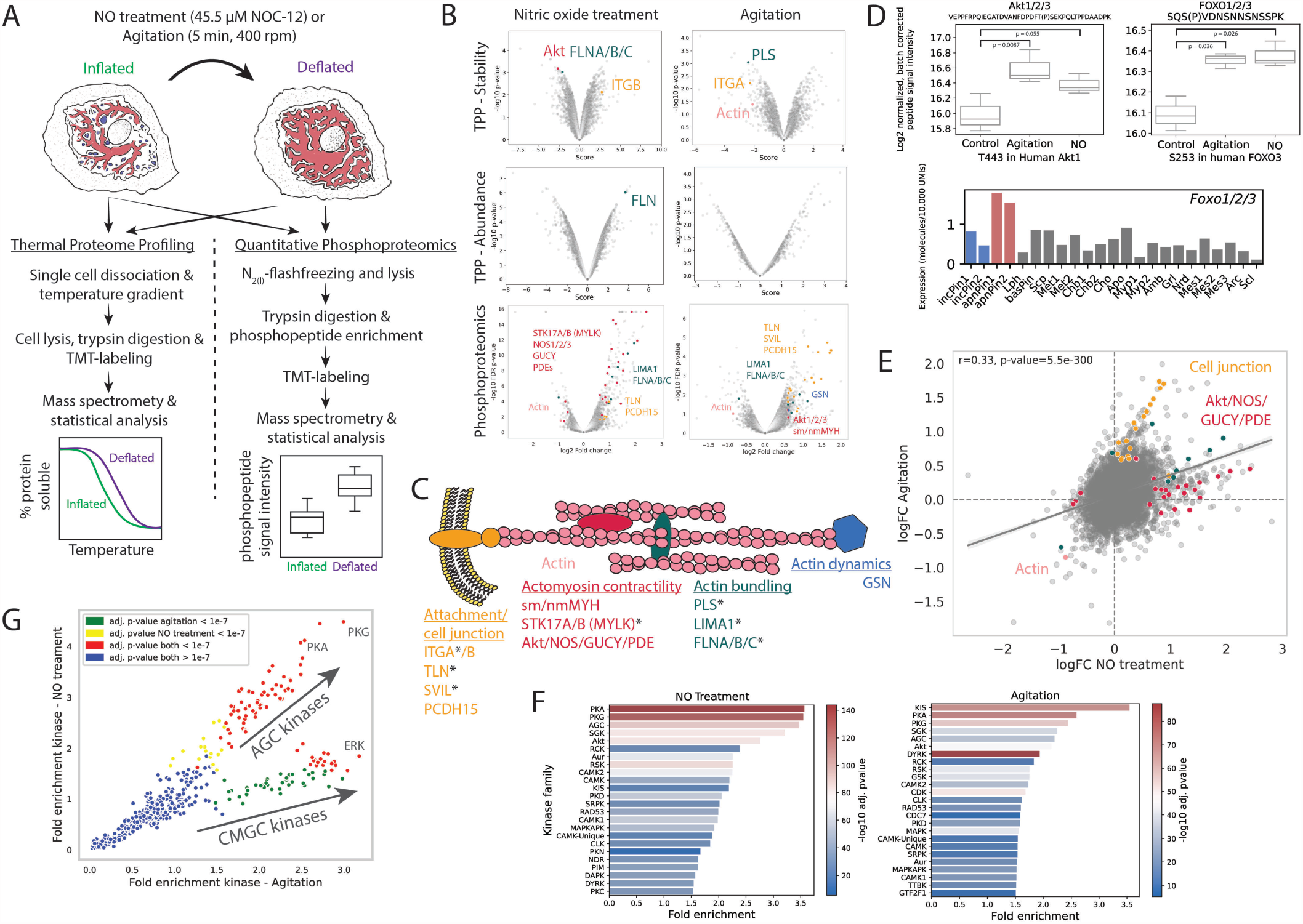
Phosphoproteomics and thermal proteome profiling of sponge deflation. **(A)** Experimental setup of sponge functional proteomics. *S. lacustris* deflation was induced by circular agitation or NO treatment. **(B)** Volcano plots of TPP and quantitative phosphoproteomic results of NO treated and agitated sponges. TPP results show individual protein changes in stability and abundance. Phosphorylation differences are quantified by unique phosphopeptides. Proteins related to stress fibers with significant changes are individually colored: actomyosin contractility (red), actin (rose), actin bundling (green), actin dynamics (blue), membrane attachment or cell junction (yellow). FLNA/B/C: filamin. GSN: gelsolin. ITGA/B: integrin alpha/beta. LIMA1: LIM-domain and actin-binding protein 1. PDEs: phosphodiesterases. PLS: plastin. PCDH15: protocadherin 15. SVIL: supervillin. TLN: talin. **(C)** Illustration of actomyosin proteins significantly changing stability, abundance or phosphorylation state. Colors relate to (B). Proteins with asterisk relate to Figure S4A. **(D)** Quantification of *S. lacustris* Akt kinase 1/2/3 phosphopeptide corresponding to *H. sapiens* Akt1 T443. Quantification of *S. lacustris* FOXO 1/2/3 phosphopeptide with phosphorylated serine corresponding to *H. sapiens* FOXO3 S253. The barplot shows normalized single-cell RNAseq expression of *Spongilla Foxo1/2/3*. Incurrent pinacocytes 1 (incPin1) and 2 (incPin2) are highlighted in blue. Excurrent/Apendopinacocytes 1 (apnPin1) and 2 (apnPin2) are highlighted in red. Lph: Lophocytes, basPin: Basopinacocytes, Scp: Sclerophorocytes, Met1: Metabolocytes 1, Met2: Metabolocytes 2, Chb1: Choanoblasts 1, Chb2: Choanoblasts 2, Cho: Choanocytes, Apo: Apopylar cells, Myp1: Myopeptidocytes 1, Myp2: Myopeptidocytes 2, Amb: Amoebocytes, Grl: Granulocytes, Nrd: Neuroid cells, Mes1: Mesocytes 1, Mes2: Mesocytes 2, Mes3: Mesocytes 3, Arc: Archaeocytes, Scl: Sclerocytes. **(E)** Correlation of phosphopeptide fold changes of NO treated and agitated sponges. Peptides of proteins highlighted in (B) and (C) are highlighted in the same colors. **(F)** Enrichment of kinase family activity after NO treatment and agitation (prediction by GPS 5.0 (63)). **(G)** Correlation of kinase activity enrichment between agitated and NO-treated sponges.

We profiled proteomic changes following NO- and agitation-induced deflation (**Figure 3A**). In total, we measured relative thermal stability and abundance changes of 5595 and 3331 proteins for NO-treated and agitated sponges, respectively. Quantitative phosphoproteomics identified and quantified 12165 unique phosphopeptides in the sponge. NO treatment resulted in quantitative changes of phosphorylation levels on 390 unique phosphopeptides mapping to 270 unique proteins and stability and abundance changes of 106 and 23 proteins, respectively (**Figure 3B**) (Full lists as Files S1, 2). In turn, agitation led to quantitative changes of phosphorylation levels on 303 unique phosphopeptides (229 proteins) and stability and abundance changes of 40 and 22 proteins, respectively (**Figure 3B**).

Corroborating the role of stress fibers, we found significant changes in phosphorylation state, stability or abundance for proteins involved in the regulation of actomyosin contractility (MYH, STK17A/B, Akt kinase, NOS1/2/3, GUCYs, phosphodiesterases (PDEs)), actin dynamics (gelsolin (GSN)), bundling (plastin (PLS1), filamin, LIMA1) and cell to cell or cytoskeleton to membrane adhesion (integrin alpha (ITGA), talin (TLN1/2), supervillin (SVIL), protocadherin 12 (PCDH15)) (**Figure 3B, C**). Some of these proteins are highly and specifically expressed in the pinacocyte family, supporting their role during relaxation (**Figure S4A**). Actin crosslinkers such as filamins, LIMA1 and PLS1 are known to play a central role in tension creation and are necessary for stress fiber bundling and stability (64–67), and phosphorylation of MYH and LIMA1 has been associated with the disassembly of contractile actin fibers (68, 69) (see above).

We also detected significant phosphorylation changes in members of the Akt/NO/PKG pathway proteins during deflation. Both agitation and NO treated sponges led to an increase in phosphorylation of a conserved Akt kinase phosphosite (T443 in human Akt1) necessary for Akt kinase activity (70) and an Akt kinase specific phosphorylation site in the transcription factor FOXO1/2/3 (S253 in human FOXO3) (71, 72), a marker of apendopinacocytes (**Figure 3D**). Phosphorylation of this site in *H. sapiens* has been shown to inhibit its transcriptional activity, promoting cell survival (73). In NO-treated sponges we observed significant changes in phosphorylation state of NOS, GUCY and pinacocyte specific PDEs, and measured a stability change of Akt kinase, corroborating their involvement (**Figure 3B**). In general, phosphorylation events of NO-treated and agitated sponges were positively correlated (**Figure 3E**), particularly in proteins related to actomyosin contractility. Whereas NO treatment led to a stronger upregulation of the phosphorylation signal in the Akt/NO/PKG signaling pathway, agitation and the apparent mechanical stress favored phosphorylation of cell adhesion proteins, suggesting a mechanosensitive role of the stress fiber adjacent cell-cell junctions (74).

To further elucidate kinase activity in NO and agitation treatments, we predicted all potential kinase-specific phosphorylation sites of proteins detected in the phosphoproteomic experiments using GPS 5.0 (63). Comparing predicted phosphorylation sites with experimentally detected upregulated phosphorylation sites of NO treated sponges (with logFC > 0.25) showed enrichment of sites that are predicted to be phosphorylated by the AGC family kinases PKG, PKA, Akt kinase and the closely related serum/glucocorticoid kinase SGK (**Figure 3F**), consistent with the activation of these pathways in NO-induced deflation of *Spongilla*. The cell type-specific expression of PKG and the expression of experimentally detected PKG targets revealed a positive correlation with cell types of the pinacocyte family exhibiting highest expression of PKG and PKG targets (**Figure S4B**). Predicted kinase activity of agitated sponges strengthened the close connection of sponge deflation and the Akt/NO/PKG pathway (**Figure 3F**). Additionally, agitated sponges showed higher predicted phosphorylation by kinases of the GMGC family, especially MAPK/ERK (**Figure 3G**). Although ERK has been shown to activate NOS (75), their primary role is the uptake of a broad spectrum of external stimuli, such as mechanical stress and stretch of actin stress fibers (76), and the translation into intracellular biochemical and physiological responses.

### Oscillatory shear stress induces an inflammatory-like response through proteinase and cytokine secretion and upregulation of reactive oxygen species (ROS)

Circular agitation of *Spongilla* induces deflation and exerts oscillatory shear stress and mechanical strain, disturbing laminar water flow. Phosphoproteomics and TPP experiments performed after circular agitation revealed a distinct signature for secretory activity. Protein hits were enriched for endo/exocytosis gene ontology terms (**Figure 4A**), and included changes in stability and abundance of secretory vesicle proteins VAMP4, SCAMP2/3/4/5 and DNAJC5/B (**Figure 4A**). To explore this further, we performed TMT-based quantitative proteomics on the surrounding medium of agitated, deflated sponges compared against an untreated control (**Figure 4B**). In total, 146 proteins were detected, 47 of which were significantly upregulated in the deflated state (**Figure 4C**) (full list as **File S3**). These proteins are enriched in bearing an N-terminal signal sequence (p=4E-09, 13.7-fold enrichment, prediction by SignalP 6.0 (77, 78)), consistent with secretion rather than cell necrosis and spilling.

**Figure 4.**
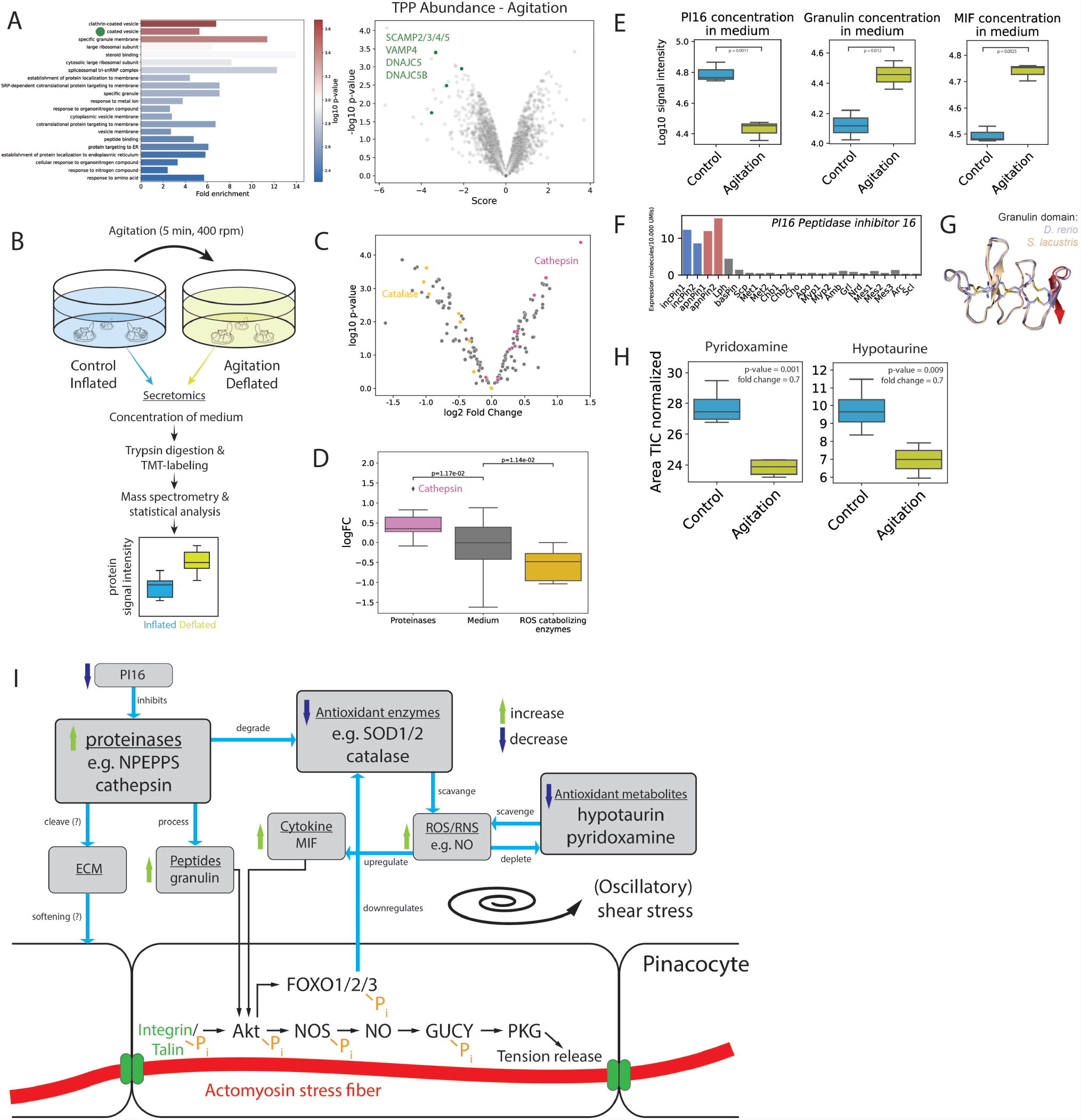
Secretomic and metabolomic profiling of agitation-induced deflated sponges. **(A)** GO term enrichment of significant protein stability changes (relates to Figure 3A). Green dot highlights GO term for proteins in volcano plot (relates to Figure 3B). SCAMP2/3/4/5: secretory carrier-associated membrane protein 2/3/4/5, VAMP4: vesicle associated membrane protein 4, DNAJC5/DNAJC5B: DnaJ homolog subfamily C member 5/B **(B)** Secretomics experimental setup. **(C)** Secreted protein fold changes, highlighting proteinases (pink) and ROS catabolizing enzymes (yellow). See also Figure S5A, C. **(D)** Quantification of secreted proteinases (pink) and ROS catabolizing enzymes (yellow). P-values were calculated using the Wilcox rank sum test. Colors relate to (C). **(E)** Relative protein abundance of PI16, Granulin, and MIF in sponge medium after agitation-induced deflation. See also Figure S5B. **(F)** *PI16* normalized expression from *S. lacustris* single-cell RNAseq atlas. **(G)** ColabFold structural prediction (79, 80) of *Spongilla* granulin domain (beige) aligned with its best morpholog (*Danio rerio* granulin domain, UniprotID: Q7T3M4, aa 95 - 147). The peptide detected in the secretomics experiment highlighted in red. Conserved disulfide bonds are shown in yellow. **(H)** Quantification of pyridoxamine (power = 0.99, precision = 15.5%) and hypotaurine (power = 0.87, precision = 9.2%) in the sponge body after agitation-induced deflation by manual integration of TIC normalized areas. **(I)** Schematic of proposed relaxant-inflammatory reaction of *Spongilla* in response to (oscillatory) shear stress.

Following agitation, we detect a significant increase of secreted proteases in the medium (Cathepsin, TPP1, CPA1/2/3/4, NPEPPS, CNDP1, PREP, PEPD, NPEPL1, LAP3) (**Figure 4C, D, I**), coinciding with a significant decrease of the proteinase inhibitor PI16 (**Figure 4E, I**), indicative of the establishment of a proteolytic extracellular environment. PI16 is highly and specifically expressed in the pinacocyte family (**Figure 4F**), whereas the proteases are more broadly expressed (**Figure S5A**). Some of these proteases have been shown to be secreted in other organisms, exhibiting diverse extracellular roles (81–83) including ECM remodeling, regulation of inflammation (84), digestion of specific protein targets or aggregates and the processing of proteins to produce biologically active peptides (85). TPP of agitated sponges showed significant changes in stability of two neuropeptide processing proteinases, endothelin converting enzyme (ECE) and an ortholog of thyrotropin-releasing hormone degrading enzyme (TRHDE) (**Figure S5F**). To detect secreted, potentially biologically active peptides, we conducted a peptide-centric, semi-tryptic search of the secretome data. We only considered enriched peptides stemming from full-length precursors that are predicted to bear an N-terminal signal sequence. SMART domain search (86) suggested the secretion of a granulin, whose identity we confirmed via structural similarity (80). The additional validation was necessary given that the sequences of biologically active peptides such as neuropeptides are often poorly conserved in evolution (87) (**Figure 4G**). Granulin is highly expressed in the peptidocyte cell type family, including choanocyte chambers and myopeptidocytes and in the archaeocyte family (**Figure S5B**). Granulins are secreted glycoproteins that play an important role in inflammation responses and are processed from pro-granulin by lysosomal cathepsins (88) which are highly secreted by the sponge (**Figure 4C, D, I**). An extracellular ER multiprotein chaperone complex has been shown to be necessary for granulin secretion as well (89). We detected six proteins (HSP90B1, P4HB, HYOU1, PDIA3, PPIB/C, HSPA5) that are part of this complex in the medium (90) (**Figure S5D, E**) as well as six subunits of the chaperonin containing CCT/TRiC complex (T-complex protein Ring Complex) (CCT2, 3, 5, 6, 7, 8) (**Figure S5D, E**). Granulins and their precursor are able to activate Akt/NOS (91) as well as MAPK kinases in endothelial cells (92), which makes them prime candidates to be upstream of Akt/NO signaling in sponge relaxation (**Figure 4I**). As a further indication of an inflammatory-like response, we detected the secretion of the pro-inflammatory cytokine macrophage migration inhibitory factor (MIF) (MIF is a short protein (120 aa) and was identified with one semi-tryptic peptide) (**Figure 4E**). In *Spongilla* we found MIF to be highly expressed across almost all sponge cell types (**Figure S5B**). Intriguingly, MIF is also known to activate Akt kinase (93) as well as ERK/MAPK (94) (**Figure 4I**). Additionally, it has been shown to be upregulated by increased reactive oxygen species (ROS) (95) (**Figure 4I**).

The production of reactive oxygen (ROS) or nitrogen (RNS) species such as NO are protective responses of the innate immune system (96, 97). We observed a decrease of ROS catabolizing, antioxidant enzymes in the medium after agitation (**Figure 4C, D**) including catalase, superoxide dismutase 1 and 2, peroxiredoxin, peroxidasin, two members of aldehyde dehydrogenases as well as an unspecified peroxidase (all showing broad expression, **Figure S5C**). Extracellular ROS and their catabolizing enzymes are well documented in model species (98). A reduction of these proteins in the supernatant could be a direct consequence of the increased proteinase activity (**Figure 4I**). In *M. musculus*, SOD has been shown to be a direct target of NPEPPS (99), which is also secreted by *Spongilla*. A decrease in ROS catabolizing enzymes could indicate an increase in ROS (**Figure 4I**). To test this, we applied an untargeted metabolomics approach which is able to detect metabolites that are differentially regulated in the sponge body due to agitation induced relaxation. ROS themselves are short lived, highly reactive molecules that are difficult to detect and quantify. However, we found two antioxidants, hypotaurine (100) as well as pyridoxamine (101) decreased after agitation, which likely reflects their depletion by the increase of ROS (**Figure 4H, I**). Additionally, catalase is a prominent target gene of FOXO transcription factors (102, 103). Inactivation of FOXO by phosphorylation of S253 by Akt kinase (see above) is shown to increase ROS concentration by the downregulation of ROS degrading enzymes such as catalase (104) (**Figure 3D, 4I**).

We thus show that oscillatory shear stress activates an inflammatory state characterized by the upregulation of ROS and the secretion of proteinases, MIF and granulin. These processes have been shown to be closely interconnected in various model systems (**Figure 4I**). Additionally, the Akt/NO pathway offers a potential intersection between inflammation and tension release of stress fibers (**Figure 4I**).

## Discussion

### Sponge deflation vs “contraction”

Our results suggest a new model for coordinated whole-body sponge movement. It posits that sponge pinacocytes are, by default, in a tensional state, manifested in the pinacocyte stress fibers which respond to oscillatory shear stress by relaxation. It follows that what was traditionally described as sponge “contraction” is instead initiated by a release of pinacocyte tension. To introduce a consistent naming convention, we propose the term “sponge contraction” to be omitted in the future and replaced by “sponge deflation” for the entire process of bodily volume reduction (see also **Supplemental Note**). The continued pumping of the choanocyte chambers concomitant with tent and ostia collapse creates low pressure and thus leads to a collapse of the incurrent canals and a sneeze-like expansion of the excurrent system. Supporting our new model, rheological measurement on marine sponges revealed initial tissue softening upon application of oscillatory shear stress (105). Actomyosin-mediated tension is energetically expensive to build up (requiring ATP hydrolysis) but can be efficient to uphold (106, 107), resulting in an energetically favorable inflated state. Many metazoans make use of similar mechanisms, most impressively the mollusc “catch” muscle, which passively sustains a tense state but can be relaxed rapidly by cAMP (108). The evolution of directed motion through isotonically contracting myocytes in Cnidaria and Bilateria from originally formgiving and stabilizing isometric stress fibers is a tantalizing scenario for the evolution of myocytes in Metazoa.

Our observation of morphological changes during the deflation of *S. lacustris* are in accordance with other related fresh-water sponges such as *Ephydatia muelleri* (8, 9, 12, 14) and marine sponges such as *Halichondria panicea* (4, 7). The collapse of the incurrent and expansion of the excurrent system is a shared feature of these demosponges. As an exception, *Tethya wilhelma* shows similarities in that the pinacoderm is responsible for the deflation (17) but also differences in that both incurrent and excurrent systems show reduced volumes and only mesohyl tissue is visible during the deflation. Also, the apparent narrowing of the osculum tip reported in *H. panicea* and *E. muelleri* (8) is not conclusively resolved. Although the upright, inflated state of the osculum seems to be accomplished by the pressure of passing water (112), circumferential actin fibers might induce an active closure of the osculum tip leading to pressure buildup and expansion of the excurrent system (4).

Signal transduction in cellular sponges - in contrast to syncytial glass sponges - has been attributed to paracrine, diffusible small molecules (49, 113) or mechanotransduction through cell tugging (114). Several agents such as cAMP, caffeine, glycine, serotonin, glutamine or GABA are able to induce or modulate the deflation in *E. muelleri, T. wilhelma* or *H. panicea* (4, 8–10). The importance of nitric oxide in the regulation of deflation has been previously highlighted in *E. muelleri* (9, 12) as well as *T. wilhelma* (10). Our work suggests that NO and cAMP both converge in PKG/PKA pathways that control tension release of pinacocyte stress fibers. Significantly, we found no evidence for an involvement of glutamate or GABA in our unbiased functional proteomics, nor were we able to reproduce glutamate-induced deflation in *Spongilla*. Nevertheless, we can not rule out that multiple pathways feed into deflation control. These may also include adrenergic or serotonergic pathways, glutamate/GABA (10) or lipid signaling such as eicosanoid leukotrienes which regulate both contractility as well as inflammation in innate immune responses, with their synthesizing enzymes highly expressed in pinacocytes (13).

A recent study in *E. muelleri* suggested that NO leads to intracellular Ca^2+^ increase and subsequent cellular “contraction” through activation of MYLK (12). We show however that p-MYL staining is already present in the inflated, pre-tensional state, consistent with our model. Their reported collapse of the incurrent system after treatment with MYH inhibitor para-Aminoblebbistatin is also congruent with our relaxation model. It appears difficult to disentangle direct cellular effects of Ca^2+^ as it has both been shown to induce a contraction through MYLK activation in smooth muscles but also to activate NOS which consequently induces relaxation (115). NO production in endothelial cells and actomyosin contractile modules in smooth muscle cells allow spatial separation of these processes in the vertebrate vasculature whereas sponge pinacocytes express both systems (**Figure 5**). Ca^2+^-imaging of the sponge will clarify the role of Ca^2+^. Finally, in contrast to our findings, treatment with MYLK inhibitor ML-7 did not induce a deflation in *E. muelleri*. Despite our effort to replicate this result using identical concentrations and suppliers, *S. lacustris* consistently deflated after treatment. Further detailed studies on the molecular physiology of incurrent and excurrent pinacocytes in both species will be needed to disentangle these results.

**Figure 5.**
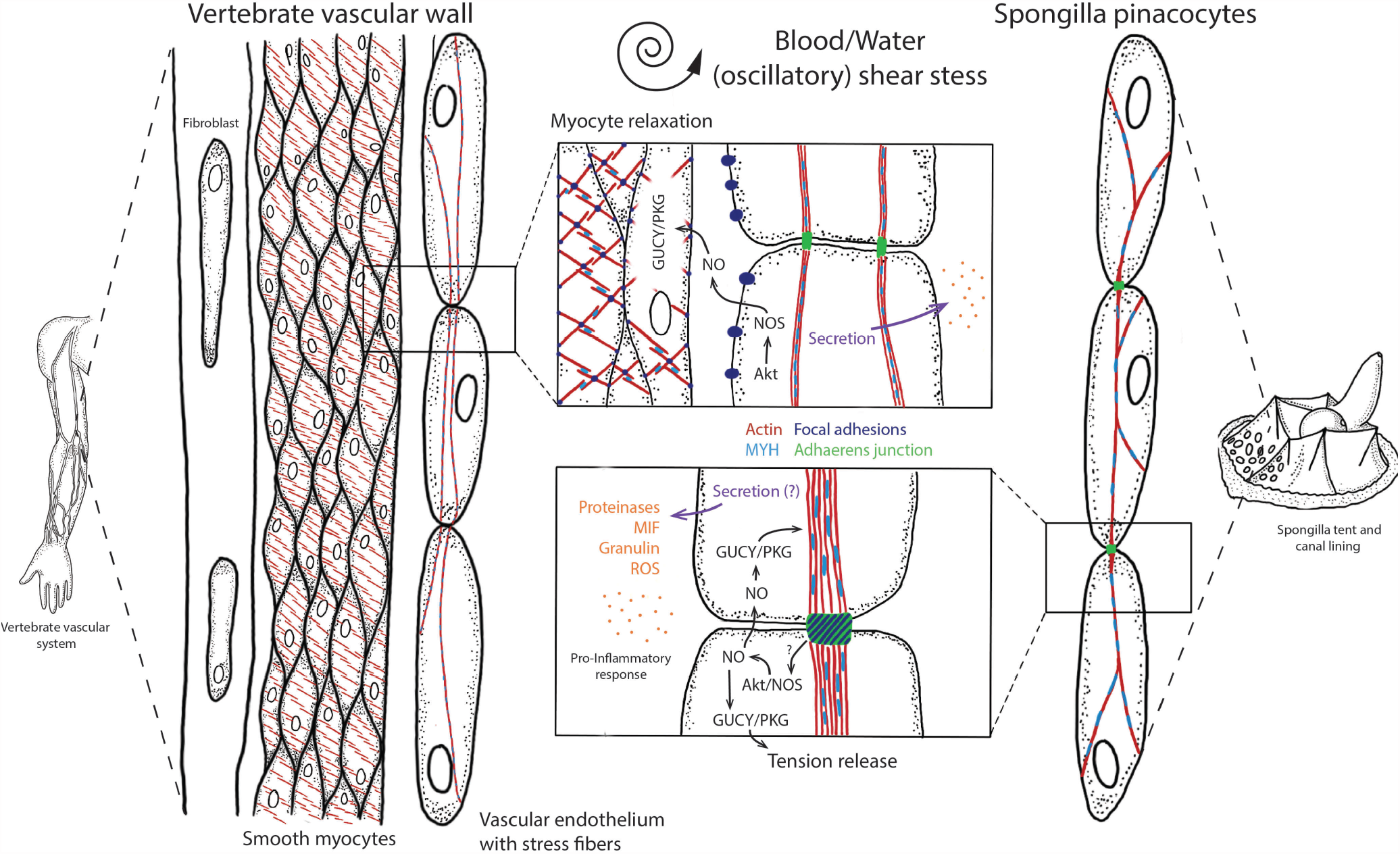
Comparison of oscillatory shear stress reactions in vertebrate vasculature and *S. lacustris* pinacoderm. Cellular components of the vertebrate vascular wall include vascular endothelial cells with actomyosin stress fibers (actin: red, myosin: light blue) oriented parallel to blood flow and connected by adherens junctions (green) (109, 110), smooth myocytes, and fibroblasts. Endothelial cells are connected to the ECM via focal adhesions (111) In the vascular system, endothelial cells adopt a sensory and signaling role whereas the smooth muscle cells are primarily responsible for the maintenance and regulation of vascular tone. *Spongilla* pinacoderm apparently assume both roles. Pinacocytes have actomyosin stress fibers connected by cell-cell junctions composed of focal adhesion and adherens junction proteins (19) Alteration of flow by oscillatory shear stress leads to Akt kinase activation and NO production, resulting in tension release or relaxation of actin stress fibers and smooth myocytes. In both systems, pro-inflammatory reactions are mediated by ROS increase and secretion of granulin, proteinases, and MIF.

### Sponge deflation involves an inflammatory state

Our most intriguing result is the resemblance of the physiological response during *Spongilla* deflation - with secretion of proteinases, granulin and MIF and the upregulation of ROS - to the inflammatory response of the innate immune system (116, 117). We show that these responses are regulated biochemically (via an Akt/NO signaling pathway) and likely transcriptionally (via FOXO inhibition). Interestingly, nitric oxide has been shown to play a central role in both the innate defense against microbial pathogens and the regulation of actomyosin contractility (118). Consistent with our data, innate immune responses to injury or microbial infection have been recorded in other sponge species (119–121). Regeneration of *H. panacea* explants suggested ECM remodeling and differential expression of genes such as cathepsins, filamins, protocadherins and integrins that are important for angiogenesis in vertebrates (119). Our work suggests that pinacocyte stress fibers take over a stabilizing and mechanosensing role (122). In this “tensegrity” (tensional integrity) model of mechanosensation (123), the state of pinacoderm tensile equilibrium would serve as both a sensor and actor (14). Tensional isometric prestress in the actin fibers allows the system to be stable in equilibrium but responsive to internal and external forces (123). These forces create shear stresses that can be monitored and - in case of deviations from the equilibrium state - translated into defense strategies, both of morphological as well as physiological nature.

### The ocean in me: Evolutionary origin of the vertebrate vascular system response from an ancient relaxant-inflammatory module

Strikingly, we found the combined sponge reactions to be highly reminiscent of those of vascular endothelial cells in vertebrates experiencing oscillatory shear stress (OSS) (124), in which atherosclerotic plaques lead to a turbulent, disturbed flow, different from “healthy” laminar flow (125) (**Figure 5**). Stress fibers formed in endothelial cells show a remarkably similar appearance to the sponge pinacocyte stress fibers (126). Mechanosensation of OSS in vascular endothelial cells is carried out by integrins and talin (74, 127, 128) and hallmark reactions of endothelial cells towards OSS such as the upregulation of MIF (129), attenuation of PI16 (124), reduced antioxidant production and increase in ROS (130) are mirrored in the reaction of the sponge following oscillatory agitation (**Figure 4I and 5**). Interestingly, OSS was shown to upregulate the expression of NOS in endothelial cells in combination with elevated ROS in the form of hydrogen peroxide (131), connecting OSS to NO-induced relaxation. Restoring laminar flow in both the vertebrate vascular and sponge canal system through a relaxant-inflammatory response appears indispensable for survival (3, 132). Disturbed flows in the canal system (e.g. due to mechanical obstructions, predation or strong currents and waves (119, 121, 133– 136)) otherwise can lead to the sponge’s starvation or infection with pathogens (120, 137). Similar responses are also found elsewhere, for example in lymphatic endothelial cells (138–140), the urothelium (141–143), cardiac myocytes (144), osteoblasts (145) or even respiratory epithelial cells (146–149). Stress response regulation via Akt/FOXO are also observed in Cnidaria such as *Hydra vulgaris* (150). The detection and evolutionary traceability of such an “relaxant-inflammatory” module suggests the conservation of an ancient response against disturbances in normal, laminar flow as part of metazoan epithelia.

## Limiations of the study

In our study the morphological as well as physiological reactions during the deflation of *S. lacustris* have been profiled using a variety of advanced tools such as 3D optical coherence microscopy, functional proteomic profiling or pharmacological compound testing. These strategies have advantages but also certain limitations. Inhibitors or activators of proteins have been developed and tested largely in model species and their specific protein orthologs. We can not say with certainty that each compound functions in an equivalent way in *Spongilla*, however the specific reaction of the sponge towards many contractility modulating compounds used speaks for a more universal applicability. We cannot exclude the possibility that NEM treatment leads to cell death to some extent.

Functional proteomics such as TPP and quantitative phosphoproteomics open a so far unprecedented and unbiased window in studying sponge behavior and molecular mechanisms. In this study, we compared the inflated state with a fully deflated state but did not consider e.g. dynamic phosphorylation changes during the deflation. Additionally, interpretation of the results relies on previous knowledge collected from well-studied model organisms. This impedes the venture into the more “hidden” biology of sponges (151). The results of the unbiased profiling presented in this study should be viewed as a starting point for further investigation once genetic manipulations are available and easily deployable in sponges.

A proximal goal would be the endogenous tagging of proteins for subsequent localization and live imaging. Such tools would allow, for instance, live imaging of actin fibers in pinacocytes, something we were not able to achieve despite multiple attempts. Likewise, live imaging of NO production or Ca^2+^-signaling would benefit from genetically encoded sensors (152, 153). Gene knockdown would additionally help to differentiate separate pinacocyte cell type populations and their role in the deflation.

## ACKNOWLEDGEMENTS

Nikolaos Papadopoulos and Martin Larrarde for their support in python coding. Scott Nichols, Alba Diz-Muños, Alison M. Sweeney, Jörg Hammel, and Leanne Strauss for the fruitful discussions and input for the manuscript. Manuel Gunkel for the help with the laser microdissection. The Arendt lab, especially Leslie Pan and Emily Savage for their constant support. F.R. has received funding from the European Union’s Horizon 2020 research and innovation program under the Marie Sklosowska-Curie grant agreement No. 764840/IGNITE as well as EMBL International PhD Program. The work was supported by the European Research Council Advanced grant 788921/NeuralCellTypeEvo (D.A.). C.M.P. was supported by a fellowship from the EMBL Interdisciplinary Postdoc (EI3POD) programme under Marie Skłodowska-Curie Actions COFUND (grant number 664726). M.M.S. is supported by the Allen Distinguished Investigator award through the Paul G. Allen Frontiers Group. R.P. and L.W. acknowledge support by the European Research Council Consolidator grant 864027/Brillouin4Life (RP) as well as from a HORIZON-INFRA-2022-TECH-01grant 101094250/IMAGINE.

## AUTHOR CONTRIBUTIONS

F.R., J.M.M., and D.A. conceptualized the study. F.R. J.M.M., M.M.S., I.B. and D.A. designed the project. F.R. and M.N. collected sponge gemmules. I.B., C.P. and F.R. performed the proteomics experiments. L.W. and F.R. performed the OCM experiments. F.R., N.M. and K.S. performed the pharmacological treatments, confocal imaging and image analysis. B.D. and F.R. performed the untargeted metabolomic experiment and analysis. A.S., L.W., E.M. and F.R. performed the OCM data processing, segmentation and analysis. L.W. designed and realized OCM system together with R.P. F.S., I.B., C.P. and M.M.S. designed and implemented the proteomics data analysis strategies; F.R. analyzed the proteomics data; M.N., M.M.S,

K.S. and R.P. contributed to the manuscript and gave advice; and F.R., J.M.M. and D.A. wrote the manuscript.

## STAR Methods

### Resource availability

#### Lead contact

Further information and requests for resources and reagents should be directed to and will be fulfilled by the lead contact, Detlev Arendt.

#### Materials availability

No new unique reagents were generated in this study.

#### Data and code availability

The code that produced the analysis as well as figures is available online. In particular:

- SpongeProt
- Project home page: https://git.embl.de/grp-arendt/spongeprot/
- Archived version: v1.0, available on Zenodo (https://doi.org/10.5281/zenodo.8116913)
- Programming language: bash, Python, R, matlab
- License: GPL 3.0

Supplemental videos, files and data are deposited in a Zenodo repository (https://doi.org/10.5281/zenodo.8116913). The mass spectrometry proteomics data have been deposited to the ProteomeXchange Consortium via the PRIDE (154) partner repository with the dataset identifiers PXD044169, PXD044171 and PXD044175. Raw metabolomics data will be deposited before publication in the MetaboLights repository (155) under the identifier MTBLS8137. Any additional information required to reanalyze the data reported in this paper is available from the lead contact upon request.

## Key resources table

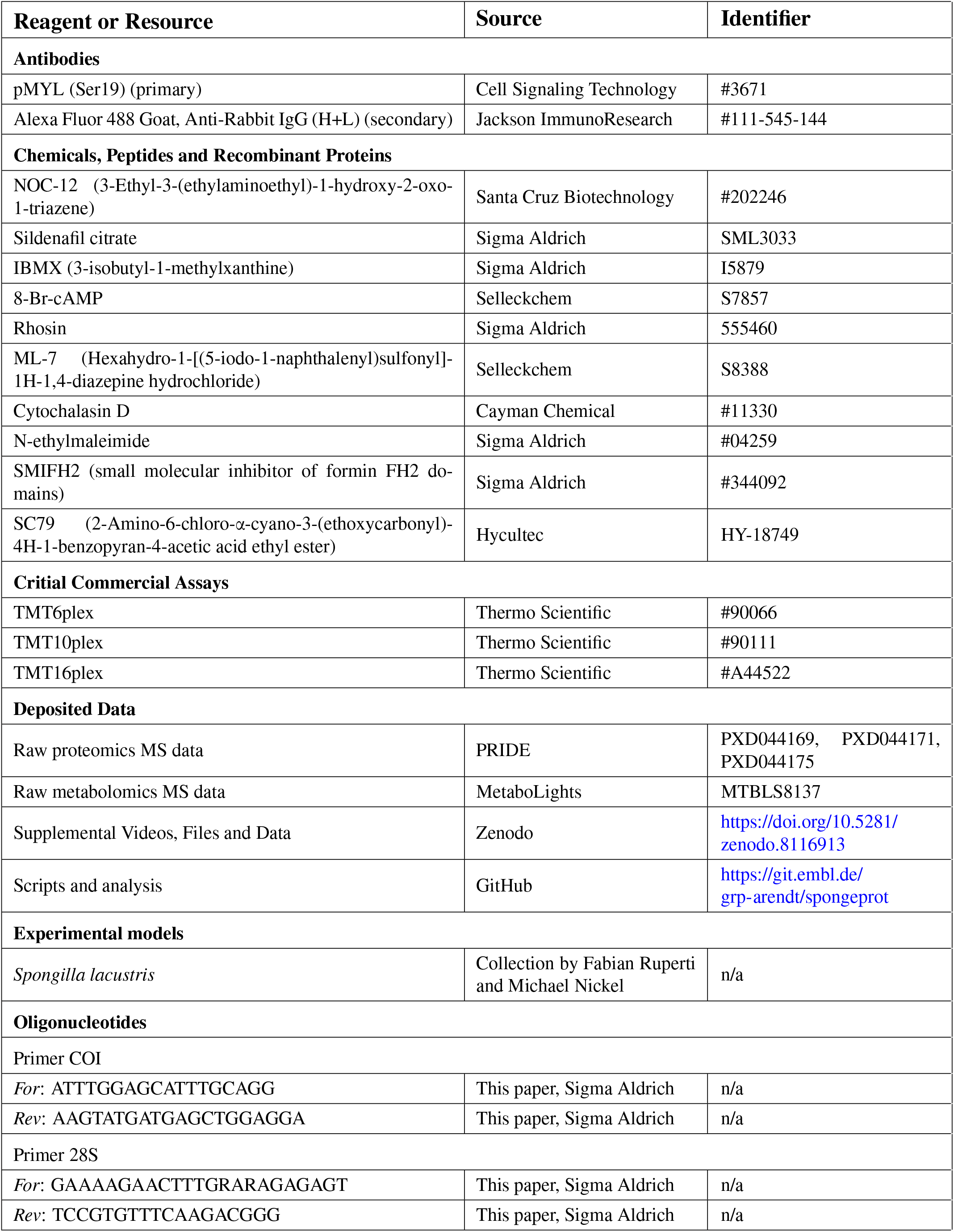
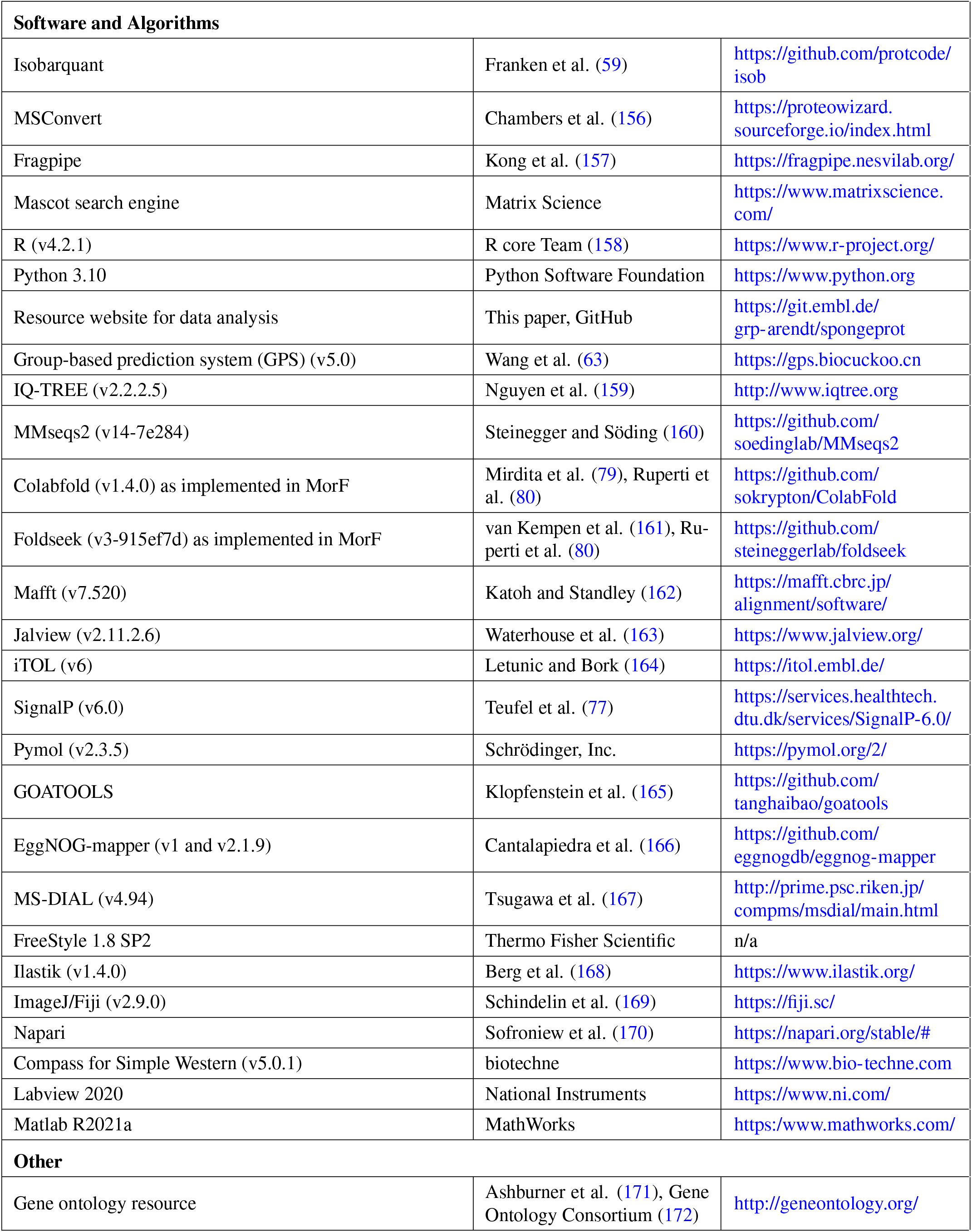

## Experimental models and subject details

### Collection and cultivation of sponges

During winter *Spongilla lacustris* specimens undergo gemmulation, a process that produces gemmules, which are protected packets of stem cells that can be collected and stored in the lab at 4°C. These gemmules can then be induced to hatch and form juvenile sponges. Gemmules of *S. lacustris* embedded in previous years sponge remains were collected on 8 March 2018, 24 February 2020, 1 December 2021, 10 January 2022 and 22 February 2023 from Lake Constance, near Kressbronn, Germany (47°35’09.0”N 9°35’56.4”E). Individual sponge patches, composed of the sponge skeleton and gemmules, were kept in lake water at 4°C until gemmule isolation. The gemmules were extracted from the adult tissue by gently rubbing the sponge patches over sandpaper (grit 500) using one finger covered in nitrile gloves. Gemmules from different sponge patches were washed and stored separately at 4°C in Vittel mineral water. Gemmules from individually cleaned patches were barcoded using 18S and COI sequences to confirm the species.

To culture the sponges, gemmules were placed in culture dishes with sterile filtered water collected from Lake Constance and kept at 18°C in the dark. Different numbers of gemmules were used depending on the experiment. To culture sponges for OCM, pharmacological testing, or confocal microscopy, 2-4 gemmules were placed in a glass-bottom culture dish (Greiner Bio-One cat# 627860) containing 5 mL of filtered lake water. To culture sponges for phosphoproteomics, thermal proteome profiling, secretomics and untargeted metabolomics, 45 - 50 gemmules were placed in a 55 mm culture dish containing 20 mL of filtered lake water or M-medium (1 mM CaCl2 x 6 H_2_O, 0.5 mM MgSO_4_ x 7 H_2_O, 0.5 mM NaHCO_3_, 0.05 mM KCl, 0.25 mM Na_2_SiO_3_) (details in the respective Methods sections).

The medium was replaced 7 days after plating when juvenile sponges adhered strongly to the glass/dish surface. After 8-9 days of growth, the juvenile sponges were sampled or fixed for experiments. At this stage, they had acquired all major features of adults, including an osculum, well-developed canal system, and numerous choanocyte chambers. The exact time at which this stage occurs can vary greatly, even for neighboring sponges grown from clonal gemmules that are present in the same culture dish. In general, only dishes were sampled or fixed in which all sponges exhibited these features.

## Method details

### Protein functional annotation

Annotation of the *Spongilla lacustris* proteome was adopted from Musser et al. (13) which followed a multi-step procedure. In short, first priority was given to orthology assignments to human genes inferred from a *Spongilla* phylome (accessible at http://spongilla.compgenomics.org/ at which all phylome trees can be searched and viewed). In cases where no human ortholog was detected, names were assigned by mapping the longest protein for each gene model to the eggNOG database (166) (version 4.5.1) using eggNOGmapper v1. In the remaining cases, the best blastp hit against human RefSeq using default parameters was used. All gene names assigned via this automated pipeline use a similar nomenclature, beginning with a gene ID (c###_g###), followed by either the orthology relationship to human genes inferred from the *Spongilla* phylome, or the name designed via emapper or blastp. Gene trees from the *Spongilla* phylome have been consulted for correct annotation and homology relationships.

As an additional method to transfer functional annotations, we implemented the structure-based homology search pipeline MorF as described in Ruperti et al. (80). In short, MorF predicts the structures of all predicted *Spongilla* proteins using ColabFold (79), an implementation of AlphaFold2 (173), and aligns the structures to available and annotated protein structures of model species. MorF-based annotations do not supersede sequence-based annotations but rather supplement annotations with structural/functional information. All MorF-based annotations in this manuscript are signalized by square brackets.

### Optical coherence microscopy and data processing

#### Optical coherence microscopy

We engineered a tailored spectral-domain Optical Coherence Microscopy (OCM) system optimized for in vivo imaging and quantitative 3D mapping of *S. lacustris* at a high and near-isotropic spatial optical resolution of ∼2.5 μm with a 4x objective lens (NA=0.13). The optical system schematic diagram of the OCM system resembles that in (24). Briefly, the light source is a superluminescent diode module (SLD, M-T-850-HP-I, Superlum Diodes Ltd, Cork, Ireland) emitting light centered at 850 nm. Its bandwidth is approximately 155 nm corresponding to a theoretical axial resolution of 2.06 μm in air. During our experiment, only the inverted part of the microscope was utilized. The laser illuminated samples through the coverslip with a power of about 1.8 mW. To minimize the impact of specular reflectance at the air-glass and glass-tissue interfaces, the normal direction of the coverslip surface and the optical axis were slightly tilted at approximately 8 degrees. This tilt angle, combined with the scale difference between the axial and lateral directions in images led to a shear transformation in the raw OCM images which was corrected in post-processing with a customized program (see below). The signal light from the sample interfered with the reference arm light at the fiber coupler and thereafter was directed to a homemade spectrometer by the fiber on the detection arm. The maximum read-out speed of the camera (EV71YO1CCL2010-BA3, Octoplus, Teledyne e2V, Saint-Egrève, France) used in the spectrometer is 250 kHz. However, in our experiment we set the exposure time of each A-line measurement to 20 μs to achieve higher image contrast, resulting in a reduced A-line rate of 50 kHz. The magnification factor and NA of the objective lens (CFI Plan Fluor, Nikon, Japan) in the microscope was 4x and 0.13, respectively. Lateral scanning of the laser beam was achieved by a galvo mirror pair which was synchronized to the camera read-out clock. Each volume scan contained 512x512 steps along x and y direction with a step size of 5.05 μm, resulting in a field of view of 2.6 mm. We note that the spatial laser scanning step resolution was approximately twice as large as the optical resolution, which we found sufficient for high-quality segmentation, but kept overall data size of the longitudinal recordings manageable. Furthermore, the actual measured axial resolution of the system was around 2.5 μm in water, slightly worse than the theoretical achievable value. This was attributed to the reduction in effective bandwidth after apodization of the interference spectra in post-processing. A wide field microscope was integrated through a dichroic mirror with cutoff wavelength 662 nm (F38-662_T3, AHF analysentechnik AG, Tübingen, Germany), assisting the sample alignment. The OCM setup has a maximum imaging speed of 250 k-Alines/s when the exposure time is set to 4 μs, resulting in a volumetric imaging speed of ∼1 Hz under 512x512 lateral scanning resolution. For the longitudinal imaging, the speed was set to ∼5 s/volume (exposure time 20 μs for each A-line) resulting in higher signal-to-noise ratio and image contrast.

#### Sample preparation and OCM imaging protocol

For live imaging, individual 8-day old *Spongilla* specimens grown in 35 mm glass-bottom culture dishes (Greiner Bio-One cat# 627860) containing 5 mL of filtered lake water and positioned on an X, Y, Z manual translation sample stage. Natural attachment of the *Spongilla* specimen to the glass bottom enabled consistent imaging using an inverted set-up. For the purpose of recording endogenous deflations, one specimen was imaged for ∼6 hours (119 volumes with ∼1 volume/3 min). Within that time, the sponge underwent 2 deflation cycles, each lasting around 30 min.

For the agitation induced deflation, the sponge specimen was agitated on a plate shaker (5 min, 500 rpm, 1.25 cm/s) with subsequent OCM image acquisition (∼1 volume / 30 sec). For nitric oxide induced deflation, 1 mL of NOC-12 (nitric oxide donor) solution was carefully added to the dish to a final concentration of 45.5 μM. OCM image acquisition (∼1 volume / 15 sec) started before the addition of NOC-12 and was stopped ∼30 min after a deflation was observed.

#### OCM raw data processing

An intensity profile of depth scan (A-line) was reconstructed from the interference spectral signal following a regular OCT postprocessing procedure including dispersion compensation, background subtraction, spectrum reshaping and inverse fast Fourier transformation (174). The fast scan along X direction generated a cross-sectional image, i.e. B-scan, containing 512 A-lines. Then, the slow scan along Y direction generated a volumetric image containing 512 B-scans. Finally, the 3D intensity dataset was scaled using a logarithmic gray scale. The raw volumetric images exhibited artifacts of distortion, tilt, and anisotropic resolution, caused by optical distortions in the scanning microscope and the tilt between the cover-slip and optical axis. These were corrected by custom post-processing scripts. Finally, the anisotropic resolution was corrected by a simple rescaling algorithm.

#### OCM Denoising

To facilitate the segmentation of the canal systems of *Spongilla lacustris* in the OCM recordings, we developed a custom 3D variant of the Noise2Noise approach based on a 3D U-net architecture for denoising the data (175). We chose a self-supervised Noise2Noise approach as it does not require any clean labels for training (**Figure S1A**). Our self-supervised deep learning method markedly suppressed the OCM image noise by an average 12 dB, which significantly facilitated down-stream post-processing and segmentation (**Figure S1B**). The 3D U-net architecture consists of an encoder-decoder structure interconnected by skip connections (176, 177). The encoder network consists of 5 hierarchically organized encoder blocks, each containing two successive units: a 3D convolutional layer, a leaky ReLU activation layer, and a group normalization layer. In parallel, the decoder network consists of 5 blocks with similar layer configurations, incorporating a nearest-neighbor upsampling layer at each level. To create the training dataset, we utilized the high acquisition rate of OCM imaging. This allowed us to obtain temporally consecutive data containing the same structural signal, but with intrinsically different noise patterns, making these noisy data pairs fulfill the main requirement of the Noise2Noise approach. Pairs of recordings with significant structural differences were removed from the training dataset to ensure the integrity of the dataset. To train the network effectively, we further augmented the dataset by randomly cropping the volume pairs into 16x64x64 patches, and applying random rotations and flips. A combination of L1 and L2 loss metrics was used to measure the difference between the prediction and the noisy target during the training process. We utilized the ADAM optimizer with hyperparameters 1=0.5, 2=0.99, and a learning rate of 1e-3, and trained the network for 300 epochs (178).

#### Segmentaion

A visual overview of the segmentation pipeline can be found in **Figure S1C**. Denoised, rotated, and scaled OCM sponge images were segmented into foreground (tissue) and background (water) using a model trained in Ilastik (v1.3.3post3) (168). Subsequently, the water pixels were divided into water “inside” and water “outside” the sponge, using a script in FIJI (v1.53t) (169). The local thickness of the inside water pixels was calculated using the LocalThickness function from the BoneJ FIJI plugin (v7.0.15) (179). Next, the “inside” water pixels were further subdivided into incurrent canal, excurrent canal, and gemmule through iterative cycles of morphological reconstructions using MorpholibJ (v1.6.0) (180) at decreasing local thickness, until no unassigned pixels in the connected regions remained. The process was initiated with manually annotated seeds, serving as starting points for each label’s reconstruction. Small isolated regions were assigned to the choanocyte chamber category. Finally, the resulting label images were manually corrected, if necessary. In the case of endogenous deflation, choanocyte chambers were separated from the excurrent canal system for the quantification of geometrical features (canal volumes and surface areas) by implementing a cut-off (7) in the local thickness map. Geometrical features were calculated using Morpholibj in FIJI. Plots were generated using python (181). Relative surface and volume changes were calculated by normalizing the measurements against one representative frame (per time-lapse) with the sponge in inflated state. Plot movies were produced from plots generated in RStudio with ggplot, and concatenated in FIJI. 3D renderings were produced using napari (v0.4.17) (170). Histogram calculations of excurrent canal diameters were performed using python (numpy v1.24.3).

### 2D Time-lase imaging, pharmacological testing and data processing

#### 2D imaging and pharmacological testing

Time-lapse imaging during the testing of pharmacological compounds was performed in a modified version from Musser et al. (13) using a consumer camera Panasonic DMC-G2 camera attached to a dissecting microscope (Zeiss AXIO) through a custom-made ocular adapter (MFT-T2, including an 8x Olympus projective). The camera was connected to a flash unit (Nikon SB24) using a connector unit (Nikon SC-17). The flash was covered by a red diffuser foil and set up next to the dishes containing the sponges. Both, camera and flash, were connected to a permanent power supply (Panasonic DMW-BLB13 and custom made). Time-lapse imaging was triggered by an intervalometer (Hongdak MC-36B) set to 30 sec intervals. Images were stored on the camera using a wifi-SDcard (ez Share), which was connected to a PC running ez Share Windows Client V1.1.0 under Microsoft Windows, transferring Images directly to a network drive after capture.

For each pharmacological compound tested, triplicate experiments of 2-4 gemmules each were placed in glass-bottom culture dishes (Greiner Bio-One cat# 627860) containing 5 mL of filtered lake water. Sponges were grown for 8-9 days before usage with a complete water exchange one day before. Stock solutions of the compounds were prepared in either ddH_2_O or DMSO, depending on the solubility. Before the start of the time-lapse, 1 mL filtered lake water was carefully removed from the plates. Before the addition of the compounds, sponges were imaged at least for 10 min to ensure that no endogenous deflation was captured. The compounds were diluted in 1 mL filtered lake water and carefully added to the plate to achieve the desired final concentration. We chose dose concentrations following empirical IC_50_ values measured in model systems (NOC-12: 45.5 μM, SC79: 15 μM, Sildenafil: 30 μM, IBMX: 100 μM, 8-Br-cGMP: 30 μM, Rhosin: 3 μM, ML-7: 1 μM, Cytochalasin D: 5 μM, N-ethylmaleimide: 1 mM, 40 μM SMIFH2), and also assessed each chemical’s toxicity in *Spongilla* through washout experiments (**Figure S2D-M**). While adding the compound, we were careful to not directly point towards the sponge to prevent a deflation induced solely due to mechanical perturbation. To further exclude the possibility that reactions evoked by compounds are due to nonspecific chemical stress, we applied toxic concentrations of hydrogen peroxide (10 μM), which did not induce the typical deflation (**Figure S2N**), as well as DMSO (0.3% (v/v)) as a control (**Figure 2C**). Exemplary time-lapse videos of each treatment are deposited in a Zenodo repository (10.5281/zenodo.8116913).

#### Data processing

The time-lapse image stacks were processed in FIJI (169) to adjust brightness, contrast, and crop the images around the sponges. Image sequences were then transferred to Ilastik (v 1.3.3) (168). Three labels were assigned to distinguish the sponge’s canal system from the surrounding tissue and background (1) incurrent canals (yellow), (2) excurrent canals (blue), and (3) remaining tissue and background (red) (**Figure S2C**). 3 - 5 images from each sequence were manually selected for training until accurate segmentations were achieved for the entire sequence. Subsequently, the simple segmentation mask was imported into FIJI and the area of the incurrent and excurrent canal system segmentations was measured and exported for plotting. For the plots, the time of compound addition was set to 0 min. The data was normalized against the first frame in each sequence (sponges in inflated state) to plot relative changes.

### Immunostaining, confocal imaging and Simple Western western blot analysis

Immunostaining was performed as described previously (12, 13, 182). For primary antibody incubation, a commercial pMYL antibody was used (1:50 dilution; Cell Signaling Technology #3671). Secondary antibody incubation was with Alexa Fluor 488 Goat, Anti-Rabbit IgG (H+L; 1:500 dilution; Cat #111-545-144, Jackson ImmunoResearch), Phalloidin-Atto 647N (1:40 dilution of 10 nM stock; Cat #65906, Sigma Aldrich), and DAPI (1 μg/ml). Experiments exclusively using Phalloidin and DAPI as stains only required fixation of the sponges and treatment with PBST. Images were taken with a 63x oil immersion or 40x water immersion objective on a Leica SP8 inverted confocal microscope. Images were processed using ImageJ (169).

To validate the specificity of the pMYL antibody to the *Spongilla* ortholog, we performed a western blot analysis using the Simple Western platform (ProteinSimple, Jess). This was performed as described previously (13). In short, clarified whole cell lysates (lysis buffer: D-PBS, 0.5% Triton X-100, 0.1% SDS, 0.4% Sodium deoxycholate, 1 mM EGTA, 1.5 mM MgCl_2_, 1 x proteinase inhibitor cocktail (cOmpleteTM, Roche), 0.26 U/μL benzonase (Millipore)) of 8-day old *Spongilla lacustris* juveniles were used. pMYL antibody validation was performed using the automated capillary-based immunoassay platform Jess (ProteinSimple). A 12-230 kDa separation module (SM-W004) for anti-rabbit chemiluminescence detection (DM-001) was prepared according to the manufacturer’s instructions. The p-MYL antibody was used at a 1:10 dilution. The run was performed using default settings and analyzed using the Compass Software for Simple Western (version 5.0.1, Protein Simple).

### Laser microdissection

Laser microdissection of peripheral tissue of inflated sponges, including the sponge epithelial tent was performed and recorded on a Leica LMD7 (349 nm laser) equipped with a 10x objective (settings: laser power: 44, aperture: 10, speed: 5, head current: 100, pulse frequency: 120 Hz).

### Thermal Proteome Profiling and sample preparation for MS

Thermal protein profiling was done similarly as previously described (183, 184). TPP for NOC-12 and agitated sponges were done in slightly different ways.

#### NOC-12 (45.5 uM) treatment

4 x 45 *Spongilla lacustris* gemmules were plated in 20 mL M-Medium (in 55 mm culture dishes) and kept at 18°C in the dark. Untreated sponges (2 replicates) were harvested and dissociated in 1.5 mL M-medium and 10 equal aliquots of 150 uL were distributed in PCR tubes. Two plates were treated with 1.5 mL M-medium containing 45.5 uM NOC-12. 5 min after treatment (deflation validated via brightfield microscopy), sponges were harvested and distributed in PCR tubes similarly. Each aliquot was heated for three minutes to a different temperature (20.0 - 30.4 - 32.6 - 37.0 - 41.0 - 44.8 - 49.9 - 55.1 - 61.1 - 67.2°C). Lysis buffer (final concentration 0.8% NP-40 (Sigma, #276855), 1.5mM MgCl_2_, protease inhibitor (Roche), phosphatase inhibitor (Roche), 0.4 U/ml benzonase (Merck, #71206-3)) was added and cells were incubated at 4°C for one hour. Protein aggregates were removed by centrifugation (0.45 um filter plate, Merck, #MSHVN4550). From the soluble fraction the protein concentration was determined (BCA assay) and for sample preparation 10 mg protein (based on two lowest temperatures) were taken further. Proteins were reduced, alkylated and digested with trypsin/Lys-C with the SP3 protocol (185). Peptides were labeled with TMT10plex (Thermo, #90111). Pairs of two temperatures for control and treated samples were combined in one TMT experiment. Pooled samples were fractionated on a reversed phase C18 system running under high pH conditions, resulting in twelve fractions (185).

#### Agitation

TPP of agitated sponges was part of an extended TPP experiment including 8 different conditions, six of which are not included and discussed in this manuscript. Only control and agitation (5 min, 500 rpm) are used in the manuscript. In total, 16 x 30 *Spongilla lacustris* gemmules were plated in 20 mL filtered lake water (collection from Lake Constance, Germany) (in 55 mm culture dishes) and kept at 18°C in the dark. Deflation was induced by agitation in two plates and confirmed via brightfield microscopy. Sponges were harvested and distributed in PCR tubes similarly. Each aliquot was heated for three minutes to a different temperature (30.4 - 32.6 - 37.0 - 41.0 - 44.8 - 49.4 - 49.9 - 50.5 - 52.6 - 57.0 - 61.1 - 64.9 - 69.5°C). Lysis buffer (final concentration 0.8% NP-40 (Sigma, #276855), 1.5mM MgCl_2_, protease inhibitor (Roche), phosphatase inhibitor (Roche), 0.4 U/ml benzonase (Merck, #71206-3)) was added and cells were incubated at 4°C for one hour. Protein aggregates were removed by centrifugation (0.45 um filter plate, Merck, #MSHVN4550). From the soluble fraction the protein concentration was determined (BCA assay) and for sample preparation 10 mg protein (based on two lowest temperatures) were taken further. Proteins were reduced, alkylated and digested with trypsin/Lys-C with the SP3 protocol (185). Peptides were labeled with TMT16plex (Thermo, A44522). Pairs of two temperatures for control and treated samples were combined in one TMT experiment. Pooled samples were fractionated on a reversed phase C18 system running under high pH conditions, resulting in twelve fractions (185).

### Quantitative phosphoproteomics and sample preparation for MS

9 x 50 *Spongilla lacustris* gemmules were plated in 20 mL filtered lake water (in 55 mm culture dishes) and kept at 18°C in the dark. Due to the fact that phosphorylation cascades often occur rapidly, treatment times for agitation and NOC-12 were shortened compared to the TPP experiment. Three plates were agitated (2 min, 500 rpm) and three were treated with 45.5 uM NOC-12 for 2 min. The induction of deflation was monitored via brightfield microscopy. For all plates (including 3 plates as control), the medium was discarded and sponges were immediately flash-frozen by dipping the bottom of the culture dishes into liquid nitrogen. This ensured immediate stop of cellular function and ensured that measured phosphorylation events are solely attributed to the treatments. The plates were thawed at 16°C for about 30 seconds before addition of 300 uL of lysis buffer (4 M guanidinium isothiocyanate, 50 mM 2-[4-(2-hydroxyethyl)piperazin-1-yl]ethanesulfonic acid (HEPES), 10 mM magnesium chloride, 10 mM tris(2-carboxyethyl)phosphine (TCEP), 1% N-lauroylsarcosine, 5% isoamyl alcohol and 40% acetonitrile adjusted to pH 8.5 with 10 M sodium hydroxide). Sponges were scraped off the culture dish surface using a cell scraper and transferred to 1.5 mL reaction cups with the lysis buffer. Samples were centrifuged at 16,000 x g for 10 minutes at room temperature to remove cell debris and nucleic acid precipitates. Three volumes of ice-cold isopropanol were then added to one volume of supernatant to induce protein precipitation, and the protein precipitates were washed 4 times with 80% isopropanol. Next, proteins were digested overnight by adding trypsin (TPCK-Trypsin, Pierce) at a ratio of 1:25 w/w trypsin:protein in 100 μL of digestion buffer (30 mM chloroacetamide, 5 mM TCEP, 100 mM HEPES, pH 8.5). After digestion, peptides were acidified with trifluoroacetic acid (TFA, final concentration of 1%) and desalted with solidphase extraction by loading the samples onto a Waters tC18 Sep-Pak 50-mg column. Samples were washed twice with 1 ml 0.1% TFA and eluted with 400 μl 50% acetonitrile acidified with 0.1% TFA before lyophilization.

For phosphopeptide enrichment, lyophilized peptides were resuspended in buffer A (70% ACN, 0.07% TFA), centrifuged at 16,000 x g, and the supernatant was added to Fe-NTA agarose beads (PureCube) on a multiscreenHTS-HV 0.45 μm 96 well filter plate (Merck Millipore). After loading, beads were washed 6 times with 200 μL buffer A, each time followed by centrifugation at 250 x g for 1 minute at room temperature. Phosphopeptides were eluted by addition of 0.2% diethylamine in 80% ACN (2 x 50 μL), before lyophilization. For TMT labeling, dried phosphopeptides were dissolved in 10 μL 100 mM HEPES pH 8.5 before addition of 4 μL of TMT reagent at a concentration of 20 μg/ μL in acetonitrile. After 1h at room temperature, the labeling reaction was quenched for 15 minutes by addition of 5 μL of 5% hydroxylamine. Labeled peptides were then pooled and subsequently lyophilized.

Labeled phosphopeptides were dissolved into 50 μL of buffer A (20 mM ammonium formate at pH 10) and loaded onto a C18 microcolumn made in-house and packed with 1 mg of C18 bulk material (ReproSil-Pur 120 C18-AQ 5 μm, Dr. Maisch) on top of the C18 resin disk (AttractSPE disks bio - C18, Affinisep). The microcolumn was first washed twice with 20 μL buffer A before phosphopeptides fractionation. The different fractions were collected after addition and elution of 10 μL of the following solutions: 3%, 5%, 7%, 9%, 11%, 13%, 15%, 17%, 19%, 21%, 23%, 25%, 27%, 29%, 31%, 33%, 35% and 40% ACN in buffer A. The fractionation was carried out in a benchtop centrifuge at room temperature and with centrifugation speed matching flow rates of approximately 10 μL/min. Each fraction was pooled with the n + 6 and n + 12 fractions, resulting in 6 fractions. The different fractions were finally lyophilized before LC-MS/MS analysis.

### Secretomic experiment and sample preparation for MS

6 x 25 *Spongilla lacustris* gemmules were plated in 20 mL M-Medium (in 55 mm culture dishes) and grown at 18°C in the dark until usage. The medium in all plates was discarded and 500 uL of fresh filtered lake water was added. For the controls (3 replicates), the water was transferred to 1.5 mL reaction cups. Three plates were agitated (5 min, 500 rpm), deflation confirmed on a brightfield microscope and the medium was transferred to reaction cups afterwards. All samples were incubated at 70 °C for 20 min to inhibit protein degradation. Samples were dried in a vacuum concentrator (eppendorf, Concentrator Plus) at 30 °C and kept at - 20 °C for further processing. Proteins were reduced, alkylated and digested with trypsin/Lys-C with the SP3 protocol (185). Peptides were labeled with TMT6plex (Thermo, #90066).

### Proteomics LC-MS/MS measurements

Measurements were performed similarly described as in Becher et al. (184). Details of chromatography and mass spectrometry runs can be found in Table S2.

### Untargeted Metabolomics sample preparation and LC-MS/MS measurement

#### Sample preparation

10 x 40 *Spongilla lacustris* gemmules were plated in 20 mL filtered lake water (in 55 mm culture dishes) and kept at 18°C in the dark until usage. For controls, the medium of 5 plates was discarded and the sponges were immediately flash frozen by dipping the bottom of the culture dishes into liquid nitrogen. To prevent thawing and potential metabolic activity, the dishes were put on metal blocks cooled to about – 80 °C by dry ice. Metal spatulas were used to scrape off the sponges and transfer into pre-cooled 2 mL cryotubes. Agitated sponges (5 replicates) (5 min, 500 rpm) were processed identically after agitation. The samples were stored at - 80 °C until further processing.

Metabolites were extracted via addition of 500 μL acetonitrile:methanol:water (2:2:1, v/v) and homogenized on dry ice with a bead beater (FastPrep-24; MP Biomedicals, CA, USA) at 6.0 m/s (3 x 30 s, 5 min pause time) using 1.0 mm zirconia/glass beads (Biospec Products, OK, USA). After centrifugation for 10 min at 15,000 x g and 4 °C with a 5415R benchtop microcentrifuge (Eppendorf, Hamburg, Germany), the supernatants were collected and residual sample pellets were reextracted with a 200 μL aliquot of the previously used extraction solvent mixture. Corresponding supernatants were combined after another centrifugation step and dried under a stream of nitrogen. Dried samples were reconstituted in 80 μL of 80% methanol, vortexed for 5 min, and transferred to analytical glass vials. The LC-MS/MS analysis was initiated within one hour after the completion of the sample preparation.

#### LC-MS/MS analysis

LC-MS/MS analysis was performed on a Vanquish UHPLC system coupled to an Orbitrap Exploris 240 high-resolution mass spectrometer (Thermo Fisher Scientific, MA, USA) in positive and negative ESI (electrospray ionization) mode. Chromatographic separation was carried out on an Atlantis Premier BEH Z-HILIC column (Waters, MA, USA; 2.1 mm x 100 mm, 1.7 μm) at a flow rate of 0.25 mL/min. The mobile phase consisted of water:acetonitrile (9:1, v/v; mobile phase phase A) and acetonitrile:water (9:1, v/v; mobile phase B), which were modified with a total buffer concentration of 10 mM ammonium acetate (negative mode) and 10 mM ammonium formate (positive mode), respectively. The aqueous portion of each mobile phase was pH-adjusted (negative mode: pH 9.0 via addition of ammonium hydroxide; positive mode: pH 3.0 via addition of formic acid). The following gradient (20 min total run time including re-equilibration) was applied (time [min]/%B): 0/95, 2/95, 14.5/60, 16/60, 16.5/95, 20/95. Column temperature was maintained at 40°C, the autosampler was set to 4°C and sample injection volume was 5 μL. Analytes were recorded via a full scan with a mass resolving power of 120,000 over a mass range from 60 – 900 m/z (scan time: 100 ms, RF lens: 70%). To obtain MS/MS fragment spectra, data-dependant acquisition was carried out (resolving power: 15,000; scan time: 22 ms; stepped collision energies [%]: 30/50/70; cycle time: 900 ms). Ion source parameters were set to the following values:

spray voltage: 4100 V (positive mode) / -3500 V (negative mode), sheath gas: 30 psi, auxiliary gas: 5 psi, sweep gas: 0 psi, ion transfer tube temperature: 350°C, vaporizer temperature: 300°C. All experimental samples were measured in a randomized manner. Pooled quality control (QC) samples were prepared by mixing equal aliquots from each processed sample. Multiple QCs were injected at the beginning of the analysis in order to equilibrate the analytical system. A QC sample was analyzed after every 5th experimental sample to monitor instrument performance throughout the sequence. For determination of background signals and subsequent background subtraction, an additional processed blank sample was recorded.

## Quantification and Statistical analyses

### Proteomics quantification and statistical analysis

#### 2D Thermal proteome profiling (2D TPP) analysis

Raw MS data was processed with IsobarQuant (59) and peptide and protein identification was performed with the Mascot 2.4 (Matrix Science) search engine. Data was searched against a custom *Spongilla lacustris* proteome (proteins with minimal length of 70 aa identified from transcriptome with TransDecoder (version 3.0.1)) including known contaminants and the reversed protein sequences. Search parameters: trypsin, missed cleavages 3, peptide tolerance 10ppm, 0.02 Da for MS/MS tolerance. Fixed modifications were carbamidomethyl on cysteines and TMT10plex (NOC12 treatment) or TMT16plex (agitation) on lysine. Variable modifications included acetylation on protein N terminus, oxidation of methionine, and TMT10plex or TMT16plex on peptide N-termini. The raw output files of IsobarQuant (protein.txt – files) were processed using the R programming language. Only proteins that were quantified with at least two unique peptides were considered for the analysis. Moreover, only proteins that were identified in at least two mass spec runs in one of the lowest temperatures (23 °C or 30.4 °C for NOC12-treatment, 23 °C or 30.9 °C for agitation treatment) and identified in at least 2 mass spec runs (NOC12-treatment) or 5 mass spec runs (agitation; part of a larger experiment comparing 8 different treatments) over all temperatures were kept for the analysis. 6314 proteins passed the quality control filters in the NOC12-treatment experiment. 3363 proteins passed the quality control filters in the agitation experiment. Raw TMT reporter ion intensities (“signal_sum” columns) were first cleaned for batch effects using limma (186) and further normalized using vsn (variance stabilization normalization (187)). Different normalization coefficients were estimated for each temperature. Abundance and stability scores were calculated as indicated in Mateus et al. (23). A treatment / control ratio was calculated for each temperature and replicate separately. The abundance score was estimated by calculating an average ratio of the first two temperatures (23 °C and 30.4 °C (NOC-12) / 30.9 °C (agitation)) for each replicate. The remaining ratios were then divided by the respective abundance average and summed up to calculate the stability score. Abundance and stability scores were transformed into a z-distribution using the “scale” function in R. The R package limma was used to estimate the significance of abundance and stability score differences for each replicate. The number of identifications in the different temperatures for each replicate has been used as a weight. The t-values (output of limma) were analyzed with the “fdrtool” function of the fdrtool package (188) in order to extract p-values and false discovery rates (fdr - q-values). A protein was annotated as a hit with an absolute score above 3 and a fdr below 0.01 and as a candidate with an absolute score above 2 and a below 0.05.

#### Quantitative phosphoproteomics analysis

Mass spectrometry raw files were converted to mzmL format using MSConvert from Proteowizard (156) using peak picking from the vendor algorithm and keeping the 200 most intense peaks. Files were then searched using MSFragger v3.4 in Fragpipe v17 against the *Spongilla lacustris* proteome (proteins with minimal length of 70 aa identified from transcriptome with TransDecoder version 3.0.1) including known contaminants and the reversed protein sequences. The search parameters were as followed: tryptic digestion with a maximum of 2 missed cleavages, peptide length = 7-50, peptide tolerance = 20 ppm; MS/MS tolerance = 10 ppm; topN peaks = 200; fixed modifications = carbamidomethyl on cysteine and TMT10plex on lysine; variable modifications = acetylation on protein N termini, oxidation of methionine and TMT10plex on peptide N termini, as well as variable phosphorylation of serine, threonine and tyrosine residues with a maximum of 4 phosphorylations per peptides. Peptide Spectrum Matches (PSMs) validation was performed by philosopher version 4.1.0 and the false discovery rate was fixed at 1% at the PSMs, peptides and proteins level. TMT quantification was performed by TMT-Integrator using default settings. The raw output files of FragPipe (psm.tsv – files) (157) were processed using the R programming language (158). Only peptide spectral matches (PSMs) with a phosphorylation probability greater than 0.75 were considered for the analysis. Phosphorylated amino acids were marked with a (P) in the amino acid sequences behind the phosphorylated amino acid and concatenated with the protein ID in order to create a unique ID for each phospho peptide. Raw TMT reporter ion intensities were summed for all PSMs with the same phosphopeptide ID. Log2 transformed summed TMT reporter ion intensities were first cleaned for batch effects using the “removeBatchEffects” function of the limma package (186) and further normalized using the vsn package (variance stabilization normalization (187)). Proteins were tested for differential expression using the limma package. The replicate information was added as a factor in the design matrix given as an argument to the “lmFit” function of limma. A phosphopeptide was annotated as a “hit” with a false discovery rate (fdr) smaller than 5% and a fold-change of at least 100%; we annotated phosphopeptides as a “candidate” with a fdr below 20% and a fold-change of at least 50%.

#### Secretomics analysis

Raw MS files were converted to mzmL format using MSConvert from Proteowizard Chambers2012-yy, using peak picking from the vendor algorithm and keeping the 300 most intense peaks. Files were then searched using MSFragger v3.4 and philosopher (4.2.1) in Fragpipe v17.1 against the custom *Spongilla lacustris* proteome (proteins with minimal length of 70 aa identified from transcriptome with TransDecoder version 3.0.1) including known contaminants and the reversed protein sequences. For the semi-tryptic search the number of enzyme termini was set to 1, whereas for tryptic search this was set to 2. Fixed modifications were carbamidomethyl on cysteines and TMT6plex on lysine. Variable modifications included acetylation on protein N terminus, oxidation of methionine and TMT6plex on peptide N-termini. The raw output files of FragPipe (157) (protein.tsv – files) were processed using the R programming language (158). Contaminants were filtered out and only proteins that were quantified with at least two unique peptides were considered for the analysis. For the detection of potential secreted biologically active peptides, we also considered proteins that were quantified with one unique peptide. Granulin and MIF (total length 120 aa) were both quantified by one unique semi-tryptic peptide. Log2 transformed raw TMT reporter ion intensities were first cleaned for batch effects using the “removeBatchEffects” function of the limma package (186) and further normalized using the vsn package (variance stabilization normalization (187)). Proteins were tested for differential detection using the limma package. The replicate information was added as a factor in the design matrix given as an argument to the “lmFit” function of limma. A protein was annotated as a hit with a false discovery rate (fdr) smaller 5% and a fold-change of at least 100% and as a candidate with a fdr below 20% and a fold-change of at least 50%.

### Gene ontology (GO) enrichment analysis

Gene ontology enrichment analysis on proteins significantly affected in abundance or stability (Thermal proteome profiling) due to agitation of the sponges has been performed with GOATOOLS (165), based on GO terms “Biological Process”, “Molecular Function” and “Cellular Compartment”. GOATOOLS allowed the input of custom gene-to-GO-term mappings. For *S. lacustris* proteins, GO terms were mapped to genes by EggNOG-mapper (emapper v2.1.9, MMSeqs2 default search against eggNOG 5.0 database) (166, 189, 190). The basic version of the GO (go-basic.obo) was downloaded from http://geneontology.org/. All quantified proteins in the experiment represented the corresponding background. Due to the low number of hits and statistical power, no p-value correction was conducted. For visualization purposes only GO-hits with a p-value < 0.005 and GO terms containing > 2 hits were taken into account. Results for “Cellular Compartment” are shown in **Figure 3A**. Further GO enrichment analysis for functional proteomic results can be found in the respective python notebook (78).

### Protein structure prediction and visualization

Structures of all *Spongilla lacustris* proteins were predicted in Ruperti et al. (80) using Colabfold (79). In this work, homologs with low sequence identity were detected by searching for structurally similar proteins using Foldseek (161). The predicted structure of *Spongilla* granulin (c100456_g4_i1_m.44114) was retrieved from ModelArchive (https://dx.doi.org/10.5452/ma-coffe-slac). Visualization and superposition of *Spongilla* granulin with its best Foldseek hit (*Danio rerio* granulin, UniprotID: Q7T3M4) was done in Pymol (v2.3.5) using the “super” command.

### Prediction of N-terminal signal sequences

For the prediction of N-terminal signal sequences in the proteome of *S. lacustris*, we used the web server for SignalP 6.0 (77). We submitted 12 jobs with approximately 3.500 sequences each (Organism: Eukarya, Model mode: fast). Prediction probability > 0.9 was chosen as the cut-off to call the presence of a signal peptide, returning 840 unique genes. Gene ontology term enrichment of these genes was conducted in the respective jupyter notebook (78).

### Prediction and enrichment of phosphosites and kinases

For the prediction of phosphosites and kinases, we used the GUI version of Group-based prediction system GPS v5.0 (63). Sequences of all proteins detected in NOC12-treated as well as agitated phosphoproteomic experiments were used as an input. All possible kinases were selected for the prediction, with the threshold set to “High”. Results were imported to python for further enrichment analysis (191). Enrichment of kinase activity due to NOC12-treatment or agitation was calculated by comparing all predicted phosphosites (and responsible kinases) with experimentally detected phosphosites. All phosphopeptides with a logFC > 0.25 were considered. Enrichments were calculated using a hypergeometric test/contingency table with all predicted phosphosites as a background. Adjusted p-values were calculated using the Benjamini/Hochberg correction.

### Calculation of maximum likelihood phylogenetic trees for MYLK kinase domain

For the calculation of the MYLK kinase domain phylogenetic tree, we used a representative selection of 15 animal proteomes, as well as the proteome of the choanoflagellate *Monosiga brevicollis*. To detect MYLK kinase domain homologs in the respective proteomes, *H. sapiens* MYLK1 (Uniprot accession number: Q15746) protein kinase domain (Pfam PF00069) (aa 1464 - 1719) was used as a query (MM-seqs2 (version 14-7e284 (160) default easy-search). Result domains were filtered (bit score > 130) and an MSA was built (domain sequences only) using Mafft (162) in default mode as implemented in Jalview web service (163, 192). The MSA was manually trimmed by removing extended (> 10 aa) gaps created by single or few sequences in the MSA.

The resulting MSA was used to calculate a maximum likelihood tree using IQ-TREE (v2.2.2.5) with the automatic phylogenetic model finder option (159, 193) (selected model: *Q.insect+I+R9* according to the Bayesian Information Criterion). Ultrafast bootstrap approximation (UFBoot) (194) with 1000 bootstrap replicates was performed. Tree visualization was done using the iTOL v6 webserver (164). For visualization purposes, branches with bootstrap values < 70 were deleted.

### Untargeted Metabolomics data analysis

Data was processed using MS-DIAL (167) and FreeStyle 1.8 SP2 (Thermo Fisher Scientific) and raw peak area data was normalized via total ion count (TIC) for metabolite quantification. Feature identification was based on accurate mass, isotope pattern, MS/MS fragment scoring and retention time matching to an in-house library. Quantification of detected metabolites in the sponge body after agitation-induced relaxation was done by manual integration of TIC normalized areas. P-values were calculated after log-transformation using two-tailed Student’s t tests. Precision was calculated as the coefficient of variation in quality control (QC) measurements (n = 5). Power was calculated via standardized effect size (Cohen’s d) at significance level α = 0.05.

### Differential gene expression in Spongilla lacustris single-cell transcriptomics data

Processed single-cell RNA sequencing data was obtained from (13, 80). Expression of genes of interest in terminally differentiated cell types was explored using scanpy (195). Code and detailed explanations for visualizing cell-type specific expression of genes (via bar plots and dotplots) are available in the corresponding jupyter notebooks (196–198).

## Supplementary Figures, Tables and Notes

**Supplement Figure 1.**
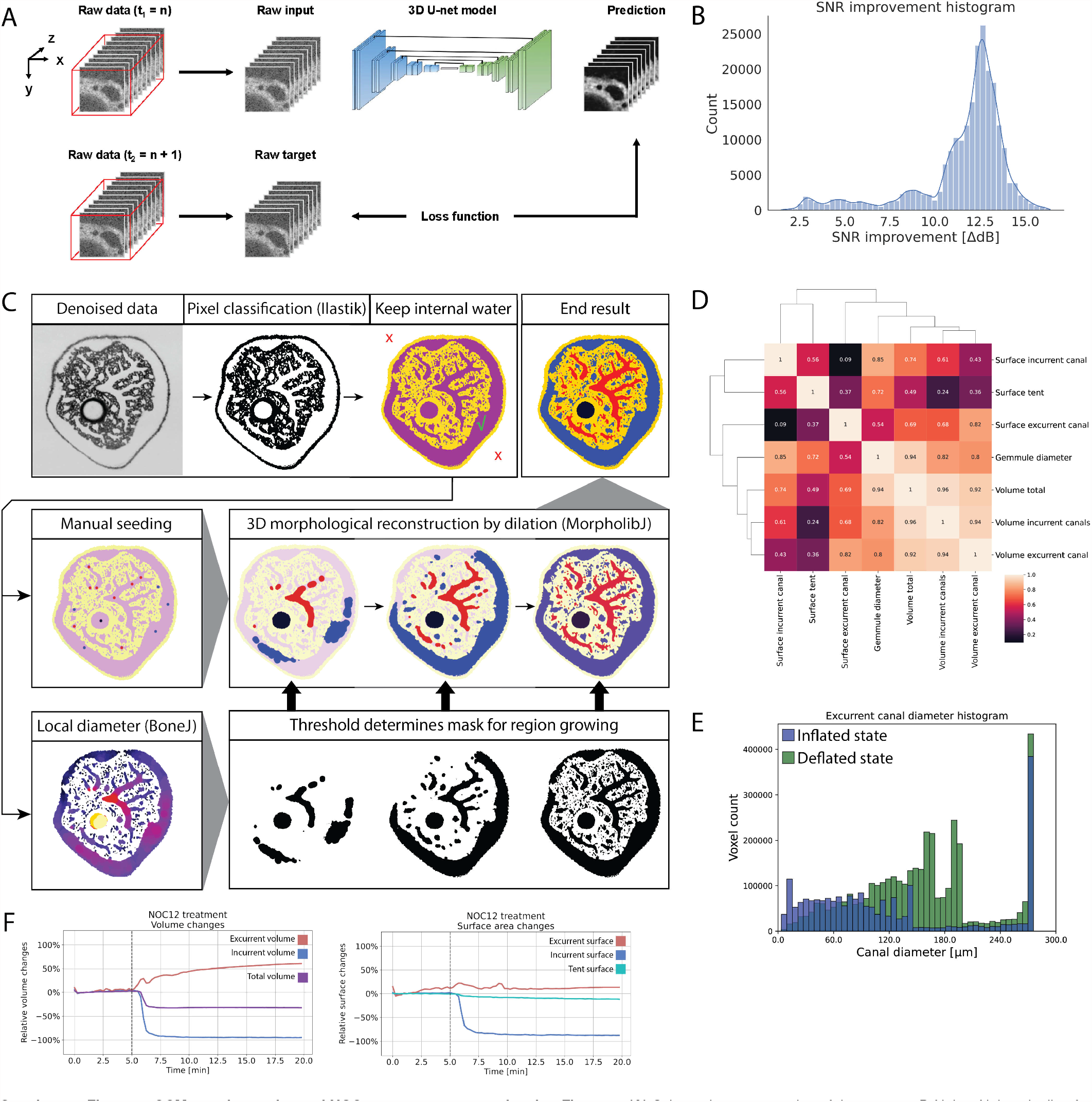
OCM morphometrics and NOC12 measurements, related to Figure 1. **(A)** Schematic representation of the custom 3D-Noise2Noise pipeline for OCM denoising, showing the 3D U-net model that is trained to minimize the discrepancy between the raw and predicted OCM images. **(B)** Histogram of OCM image SNR improvement due to denoising, calculated over 143.000 OCM image pairs (i.e. raw vs. denoised). **(C)** Schematic representation of OCM image segmentation pipeline. **(D)** Pearson correlation of morphometrics of 8-day old *Spongilla* specimens (n=5) in the inflated state. Incurrent, excurrent and total volume as well as gemmule diameter show positive correlation. With growing gemmule size (corresponding to a larger quantity of undifferentiated stem cells), the bigger the sponge and its canal system will grow under equal conditions. **(E)** Histogram of voxel counts per canal diameter in the excurrent canal system of a sponge in inflated state (blue) vs. deflated state (green). In the deflated state, excurrent canals shift to larger diameters, equivalent to an expansion of the excurrent system. **(F)** OCM measurements of NOC-12 (Nitric oxide donor) treated *Spongilla* specimen. Relative volume (incurrent (red), excurrent (blue) system and total sponge (purple)) and surface area (incurrent (red), excurrent (blue) and tent (turquoise)) changes from a single *Spongilla* specimen during NOC-12 (45.5 μm) induced deflation. Dashed line highlights compound addition.

**Supplement Figure 2.**
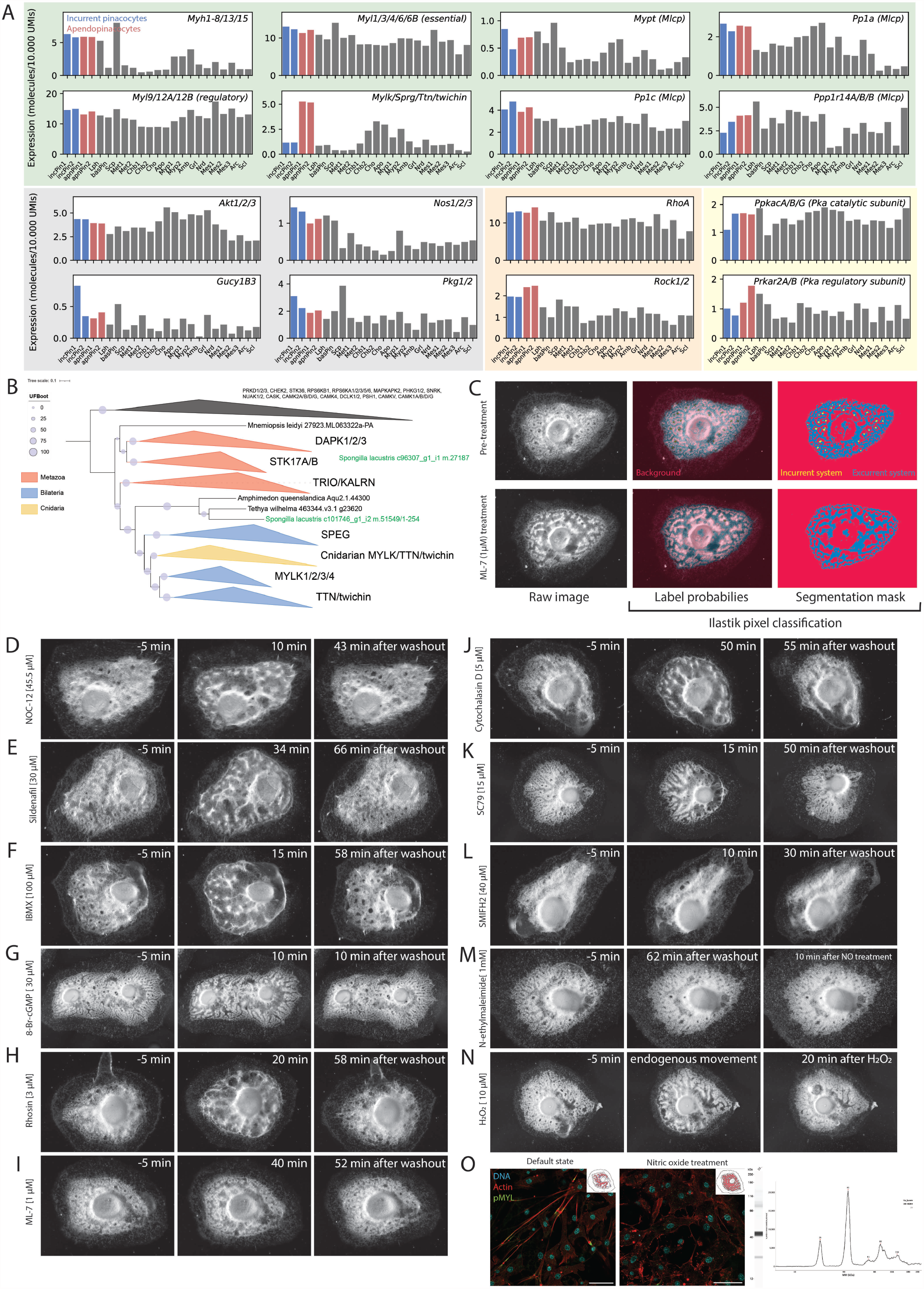
(Single cell expression) survey of contractile genes and quantification and reversibility of pharmacological treatments, related to Figure 2. **(A)** Normalized single-cell RNAseq expression of actomyosin activity and regulatory genes. Background colors refer to pathways depicted in **Figure 2A**. Incurrent pinacocytes 1 (incPin1) and 2 (incPin2) are shown in blue. Excurrent/Apendopinacocytes 1 (apnPin1) and 2 (apnPin2) are shown in red. Lph: Lophocytes, basPin: Basopinacocytes, Scp: Sclerophorocytes, Met1: Metabolocytes 1, Met2: Metabolocytes 2, Chb1: Choanoblasts 1, Chb2: Choanoblasts 2, Cho: Choanocytes, Apo: Apopylar cells, Myp1: Myopeptidocytes 1, Myp2: Myopeptidocytes 2, Amb: Amoebocytes, Grl: Granulocytes, Nrd: Neuroid cells, Mes1: Mesocytes 1, Mes2: Mesocytes 2, Mes3: Mesocytes 3, Arc: Archaeocytes, Scl: Sclerocytes. **(B)** Maximum likelihood phylogeny of MYLK kinase domain. The tree was built using *H. sapiens* MYLK1 (Uniprot accession number: Q15746) protein kinase domain (Pfam PF00069) (aa 1464 - 1719) as a query against representative animal proteomes and the proteome of the choanoflagellate *M. brevicollis*. Branches were collapsed according to gene orthogroups. The color of branches indicate the existence of members from across Bilateria (blue), Metazoa (red) or Cnidaria only (yellow). *Spongilla lacustris* genes are highlighted in green. The tree was rooted using kinase domains outside of the MYLK kinase family as an outgroup. Ultrafast bootstrapping results are depicted as the size of the circles sitting on the branches. The analysis showed that sponges, including *Spongilla*, possess one ortholog of the MYLK/TTN/twichin/SPEG kinase family. Additionally, *Spongilla* has a STK17A/B ortholog. Tree visualization was done using the iTOL v6 webserver (164). **(C)** 2D time-lapse image segmentation and analysis. Exemplary 2D brightfield microscopy images of a juvenile *S. lacustris* specimen before and after treatment with MYLK inhibitor ML-7 (1 uM). Pixels of raw brightfield images were classified using the pixel classification workflow in Ilastik (v1.4.0) (168) Labels used are incurrent canal system (yellow), excurrent canal system (blue) and background (red). Segmentation masks were exported and the areas of incurrent and excurrent canal system were measured in Fiji (169) and analyzed in R. **(D)-(N)** Washout/toxicity experiments of pharmacological compound treatments on *S. lacustris*. Brightfield images of juvenile *S. lacustris* specimens treated with pharmacological compounds before and after treatment as well as after washout. (D: Nitric oxide donor NOC-12 (45.5 μM), E: Sildenafil (30 μM), F: IBMX (100 μM), G: 8-Br-cAMP (30 μM), H: Rhosin (3 μM), I: ML-7 (1 μM), J: Cytochalasin D (5 μM), K: SC79 (15 μM), L: SMIFH2 (40 μM)). After application and induction of the respective behavior, filtered lake water was replaced twice and the health of sponges was monitored by observing their capability to return to their original state without apparent toxic effects. For (D) - (L), images on the left side show the sponge 5 min before application of the compound. Images in the center show the sponge reacting to the compound as described in **Figure 2**. Images on the right side show the sponge after water exchange (after the indicated time). For treatment with N-ethylmaleimide (NEM) (1 mM) (M), the center image shows the sponge 62 min after washout (after 10 min incubation in NEM). The right image shows the sponge 10 min after treatment with 45.5 μM NOC-12. NEM irreversibly forms covalent bonds with cysteine residues and is not expected to lose its capability to block NO-induced movement after washout. Treatment with toxic concentrations of hydrogen peroxide (H_2_ O_2_) (10 μM) (N) leads to clearly observable cell and tissue necrosis, including formation of gas bubbles due to endogenous catalase presence. This contrasts an endogenous deflation (center image) of the same sponge before treatment with H_2_ O_2_. **(O)** Immunostaining of tent pinacocyte actin fibers with p-MYL antibody. Confocal images of tent pinacocytes immunostained for phosphorylated myosin light chain (MYL, p-Ser19) (green, Cell Signaling Technology 3671), phalloidin (red), and Hoechst (cyan), reproducing the results of (12) The left image shows localization of p-MYL signal to the actin fibers of sponges in the inflated (tense) state as highlighted by the illustration in the right top corner (incurrent system: blue, excurrent system: red). p-MYL signal often shows localization close to cell-cell junctions points. The right image shows the same staining of a second specimen after treatment with 45.5 μM NOC-12. Actin fibers and p-MYL signals are concurrently absent. Scale bars, 30 um. Antibody specificity was determined using Simple Western western blot analysis of p-MYL antibody with *S. lacustris* protein lysate as input. Chemiluminescent signal plot (and virtual gel representation) of the Simple Western results show signals at 26 and 43 kDa. The theoretical mass of *S. lacustris* regulatory p-MYL is 19.4 kDa. The western blot signals likely are showing the monomeric regulatory p-MYL and a potential dimeric form or the interaction with the essential MYL (16.7 kDa).

**Supplement Figure 3.**
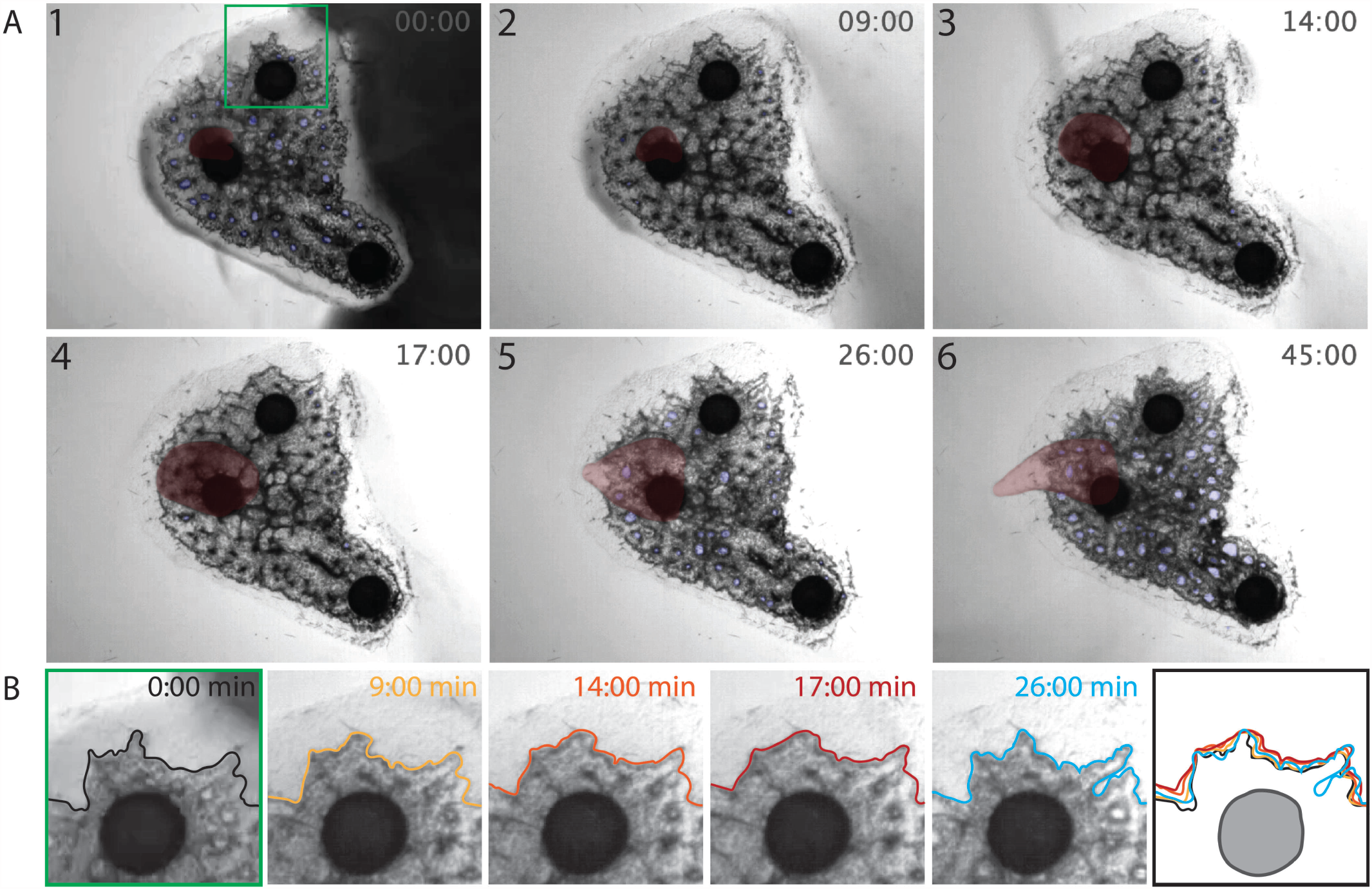
Time-lapse imaging of a juvenile *S. lacustris* after application of inedible ink particles, related to Figure 2. **(A)** Frame 1 shows the application of ink particles on the right side of the sponge. Incurrent canals (blue) are open and the osculum (red) is visible. The green square is magnified in (B). After 9 min (Frame 2), some of the ink has been ingested by the sponge which in turn reacts with the collapse and closure of the incurrent canal system and the simultaneous expansion of the excurrent system. In this state, the sponge choanocyte chambers continue to pump, leading to the growth and extreme dilation of the osculum in Frame 3 and 4 filling with water and ink particles. During this time, ink particles are visibly accumulating in the osculum. Only after about 20 min (Frame 5 and 6), the incurrent canal system opens again, concurrent with a size reduction of the osculum, the expulsion of ink particles and the return of the sponge in the inflated state (Video available in the Zenodo repository). **(B)** Expansion of peripheral excurrent canals of the same sponge. Between 0 and 17 min, excurrent canals are visibly expanding outwards, only possible through passive expansion. Contraction of excurrent canals at 26 min leads to a return in the original state.

**Supplement Figure 4.**
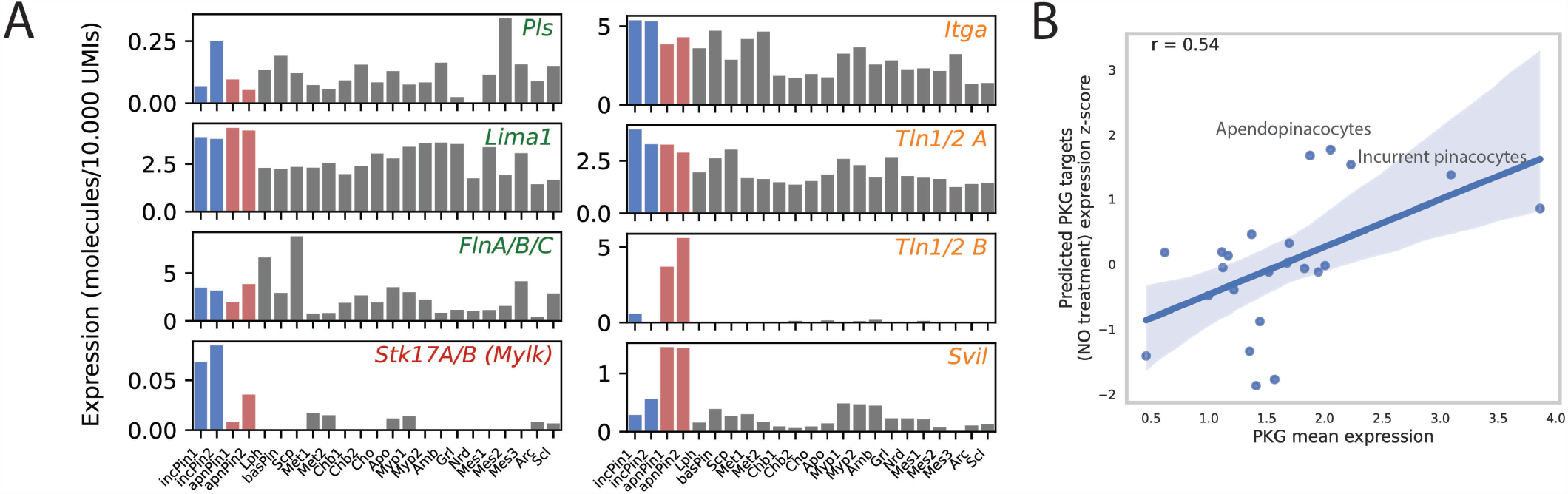
Single cell RNAseq expression of stress fiber genes showing significant differential phosphorylation and/or changes in stability and abundance due to deflation of the sponge, related to Figure 3. **(A)** Scaled single cell RNAseq expression of selected genes from **Figure 3B** and C. Incurrent pinacocytes 1 (incPin1) and 2 (incPin2) are shown in blue. Excurrent/Apendopinacocytes 1 (apnPin1) and 2 (apnPin2) are shown in red. Lph: Lophocytes, basPin: Basopinacocytes, Scp: Sclerophorocytes, Met1: Metabolocytes 1, Met2: Metabolocytes 2, Chb1: Choanoblasts 1, Chb2: Choanoblasts 2, Cho: Choanocytes, Apo: Apopylar cells, Myp1: Myopeptidocytes 1, Myp2: Myopeptidocytes 2, Amb: Amoebocytes, Grl: Granulocytes, Nrd: Neuroid cells, Mes1: Mesocytes 1, Mes2: Mesocytes 2, Mes3: Mesocytes 3, Arc: Archaeocytes, Scl: Sclerocytes. **(B)** Correlation (r=0.54, Pearson) of mean cell-type specific PKG (protein kinase G) expression and cell-type specific z-scores of GPS 5.0 predicted and experimentally confirmed (with log fold change > 0.25) PKG phosphorylation events after NOC-12 treatment. Cell types of the pinacocyte family show both high PKG as well as PKG target gene expression.

**Supplement Figure 5.**
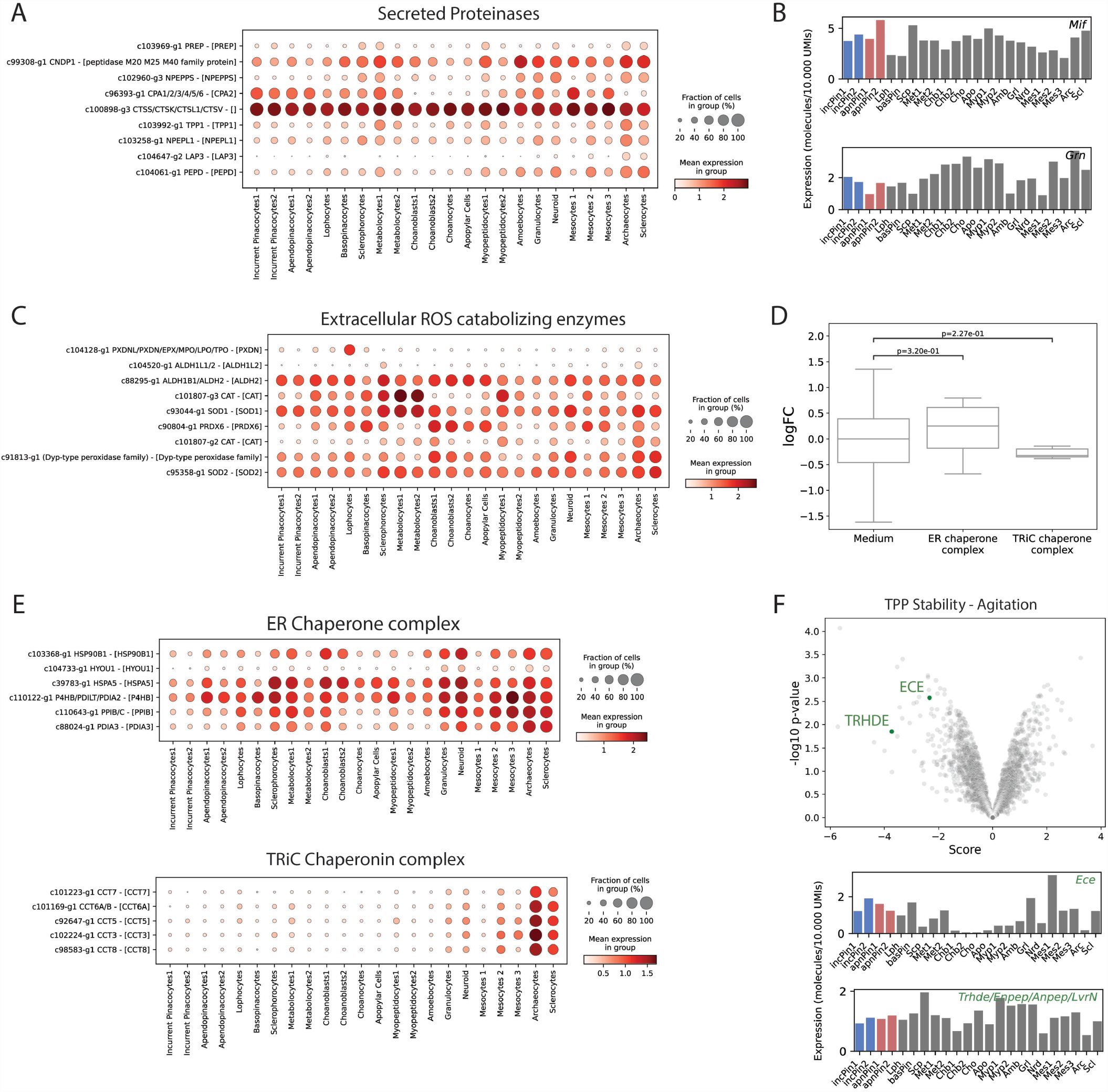
Protein quantification and scRNA-seq expression of secreted proteinases, ROS catabolizing enzymes, MIF, granulin and chaperon complexes, related to Figure 4. **(A, C)** Dotplots showing the expression of (A) secreted proteinases and (C) ROS catabolizing enzymes. Gene names in square brackets are annotated by structural alignment via MorF (80). **(B)** Scaled single cell RNAseq expression values of MIF and granulin. Incurrent pinacocytes 1 (incPin1) and 2 (incPin2) are shown in blue. Excurrent/Apendopinacocytes 1 (apnPin1) and 2 (apnPin2) are shown in red. Lph: Lophocytes, basPin: Basopinacocytes, Scp: Sclerophorocytes, Met1: Metabolocytes 1, Met2: Metabolocytes 2, Chb1: Choanoblasts 1, Chb2: Choanoblasts 2, Cho: Choanocytes, Apo: Apopylar cells, Myp1: Myopeptidocytes 1, Myp2: Myopeptidocytes 2, Amb: Amoebocytes, Grl: Granulocytes, Nrd: Neuroid cells, Mes1: Mesocytes 1, Mes2: Mesocytes 2, Mes3: Mesocytes 3, Arc: Archaeocytes, Scl: Sclerocytes. **(D)** Quantification of secreted chaperon complexes (ER chaperon complex, TRiC chaperon complex) after agitation-induced deflation. P-values were calculated using the Wilcox rank sum test. Chaperon complexes were detected in the medium but did not change concentration significantly due to agitation. **(E)** Dotplots showing the expression of components from the ER chaperon and TRiC chaperon complexes. Gene names in square brackets are annotated by structural alignment via MorF (80). **(F)** Peptide hormone processing enzymes in TPP experiment of agitated sponges. The volcano plot shows the stability changes of Endothelin-converting enzyme (ECE) and thyrotropin releasing hormone degrading enzyme (TRHDE) after agitation. Single cell expression of the corresponding transcripts is shown in the bar plots (cell type description identical to (B)).

**Supplement Table 1.** Morphometrics of juvenile *Spongilla lacustris* specimens (n=5) in inflated state, related to Figure 1. Mean values and respective standard deviation of incurrent and excurrent system volume/surface area, total sponge volume as well as tent surface area were measured using the segmented OCM images (Morpholibj in FIJI). Measurement of excurrent canal system volume/surface area as well as tent surface area excluded the upper osculum portion owed to decreasing resolution along the z-axis. The volume of the incurrent canal system is between 1.6 and 2.2 times larger than that of the excurrent system. Morphometric correlations are shown in Figure S1D.

| Morphometric | Value |
| --- | --- |
| Total volume | $0.47 \pm 0.08 \text{ mm}^3$ |
| Incurrent canal system volume | $0.13 \pm 0.04 \text{ mm}^3$ |
| Excurrent canal system volume | $0.077 \pm 0.029 \text{ mm}^3$ |
| Incurrent canal system surface area | $10.54 \pm 2.20 \text{ mm}^2$ |
| Excurrent canal system surface area | $3.95 \pm 1.13 \text{ mm}^2$ |
| Tent surface area | $5.54 \pm 0.70 \text{ mm}^2$ |

**Supplement Table 2.** LC-MS/MS Parameters used for TPP, Phosphoproteomics and Secretomics experiments, related to STAR Methods.

| Parameter | TPP | Phosphoproteomics | Secretomics |
| --- | --- | --- | --- |
| LC equipment | UltiMate 3000 RSLC nano LC system (Thermo Fisher Scientific) equipped with a trapping cartridge (Precolumn C18 PepMap 100, 5 mm, 300 mm i.d. x 5 mm, 100 Å) and an analytical column (Acclaim PepMap 100, 75 mm x 50 cm C18, 3 mm, 100 Å). Solvent A 99.9% LC-MS grade water (Thermo Scientific) with 0.1% formic acid, solvent B 99.9% acetonitrile (Thermo Scientific) with 0.1% formic acid. Flow rate loading 30 µl/min, separation at 0.3 mL/min. Pico-Tip Emitter 360 mm OD x 20 mm ID; 10 mm tip (New Objective). |  |  |
| LC settings |  |  |  |
| Gradient length [min] | 120 | 120 | 60 |
| Gradient | 2% to 4% B in 4 min, from 4% to 8% in 2 min, 8% to 28% in 96 min, and from 28% to 40% in another 10 min. A column cleaning step using 80% B for 3 min was applied before the system was set again to its initial conditions (2% B) for re-equilibration for 10 minutes | 2% to 6% B in 4 min, from 6% to 8% in 2 min, 8% to 26% in 94 min, and from 28% to 40% in another 12 min. A column cleaning step using 80% B for 3 min was applied before the system was set again to its initial conditions (2% B) for re-equilibration for 10 minutes | 2% to 4% B in 4 min, from 4% to 8% in 2 min, 8% to 28% in 42 min, and from 28% to 40% in another 4 min. A column cleaning step using 80% B for 4 min was applied before the system was set again to its initial conditions (2% B) for re-equilibration for 4 minutes |
| MS equipment and settings |  |  |  |
| MS equipment | Q Exactive Plus (Thermo Fisher Scientific) using a Nanospray-Flex ion source | Orbitrap Fusion Lumos (Thermo Fisher Scientific) using a Nanospray-Flex ion source | Orbitrap Fusion Lumos (Thermo Fisher Scientific) using a Nanospray-Flex ion source |
| Spray voltage [kV] | 2.3 | 2.4 | 2.4 |
| Capillary temperature [C] | 275 | 275 | 275 |
| Full scan mass range [m/z] | 375-1200 | 375-1400 | 375-1500 |
| MS1 Orbitrap resolution | 70000 | 120000 | 60000 |
| MS1 Maximum IT [ms] | 10 | 50 | 50 |
| MS1 AGC target | 3e6 | 4e5 | 4e5 |
| MS2 resolution | 35000 | 30000 | 15000 |
| MS2 AGC target | 2e5 | 1e5 | 1e5 |
| MS2 maximum IT [ms] | 120 | 110 | 54 |
| TopN | 10 | Top 3s | 10 |
| MS2 isolation window | 0.7 | 0.7 | 0.7 |
| MS2 fixed first mass | 100 | 110 | 110 |
| MS2 nCE | 32 | 36 | 36 |
| MS2 dynamic exclusion [s] | 30 | 20 | 60 |

## Supplemental Video Descriptions

**Video S1: Animation of OCM 3D reconstruction of a *Spongilla* specimen in inflated state, related to Figure 1**.

**Video S2: Time-lapse of a segmented sponge specimen (orthogonal view) during endogenous sponge deflation, related to Figure 1**.

**Video S3: 3D time-lapse video of segmented excurrent and incurrent canal systems during endogenous sponge deflation, related to Figure 1**.

**Video S4: Laser microdissection of peripheral tent regions of a sponge in inflated state, related to Figure 2**.

## Supplemental Files Descriptions

**Suppl. File 1: TPP result tables**

**Suppl. File 2: Quantitative phosphorylation result tables**

**Suppl. File 3: Secretomics result tables**

## Supplemental Note: Terminology of the sponge “movement”

The specific “movement” of sponges discussed in this article has fascinated humans throughout millennia, dating back to descriptions of **Aristotle** in his *History of Animals*, written around **350 B.C**. There, Aristotle notes that sponges are known to sense their surroundings and as evidence (“σημειν”) he provides the fact that the sponge “collects himself” (“συναγει εαυτν”) when disturbed, indicating an inward movement (“συναγει”) by the organism itself (“εαυτν”), i.e. as a clear intrinsic behavior, as opposed to something facilitated by external forces. A sponge “*collects himself if it perceives any purpose of tearing it up, and renders the task more difficult. The sponge does the same thing when the winds and waves are violent* […]”. Interestingly, even then, this movement appeared to be debated, as “*there are some persons who dispute this, as the natives of Torona*”. Almost identical to Aristotle, **Plinius the Elder** (*The Natural History*, 77 B.C.) describes how sponges “contract” when strong waves are hitting them (“*Intellectum inesse iis apparet, quia, ubi avulsorem sensere, contractae multo difficilius abstrahuntur; hoc idem fluctu pulsante faciunt*.”). The term “contraction” (from Latin “con-” meaning “together” and “trahere” meaning “to pull”) has survived the centuries and can be found in **Ellis and Knight** (1765) (199), describing the contraction of “fecal orifices” (ostia) on the surface of the sponge. For this, they use the pairs “contraction” and “dilation” as well as “systole” and “diastole”. In the 19th and 20th century many authors described their observations using the antonyms “contraction” and “relaxation” (**Lieberkühn (1859)** (200): “Contractionszustant”, **McNair (1923)** (112): “Contraction/Constriction” vs “Relaxation” in *Ephydatia muelleri*, **Pavans de Ceccatty (1960)** (201): “Contraction” in *Tethya lyncurium*, **Prosser (1962)** (202): “Contraction” vs “Relaxation” in multiple marine and freshwater species, **Emson (1966)** (114): “Contraction” vs “Relaxation in *Cliona celata*, **De Vos, Van de Vyver (1981)** (203): “Contraction spontanée” in *E. fluviatilis*). **Parker (1910)** (204) used slightly different descriptions for the different states such as “shriveled, rugose” vs. “filled out” or “contraction” vs “expansion” when he worked on *Stylotella heliophila*. In the late 20th century, **Weissenfels (1984, 1990)** worked extensively on freshwater sponges such as *E. muelleri* and *S. lacustris*. Whereas in earlier publications, he describes his observations as “endogene Kontraktionsrhythmik” (“endogenous contraction rhythm”) (205), at later stages he uses the term “condensation rhythm” (Latin “con-” meaning “together”, “densus” meaning “dense”) to potentially avoid the implication of active processes (25). He states that the condensation of the mesohyle “*giv*[es] *the impression of a contraction*” and that “*rhythmic changes* […] *at first glance appear to be contractions* […]. [A]*ctually however, the process is a condensation initiated by* […] *a rapid swelling of all mesenchymal cells*” (25). Despite this, **Nickel (2006, 2007, 2011), Leys (2007, 2010)** and **Nichols (2021)** used different descriptors for different phases of the behavior. In general, Nickel used the terms “contraction” vs. “expansion” in *Tethya wilhelma* (10, 11). In the expanded state, all canal systems are “inflated” (17). Leys introduced the term “inflation-contraction” response in *E. muelleri* (8, 9). “Inflation” suddenly described the “contraction” of McNair, De Vos and Weissenfels in which the excurrent canal system of freshwater sponges is expanding and the top view gives the impression of an “inflation”. In contrast, “contraction” describes the return into the default state. At the same time, this return is called “relaxation”. Most recently, Nichols returned to “contraction cycle” describing the whole-body volume reduction in *E. muelleri* (12).

This list gives an account of the complication regarding the consistent and precise terminology of the observation of sponge movement. We devised a straightforward naming convention, taking the history of this field into account and combining it with new insights from this study. For this, we recognized the importance of separating different levels of organization: (1) organismal (whole-body) changes, (2) canal changes and (3) cellular changes (see Table below). On an organismal level (1), we decided for a terminology agnostic to the cellular processes and would therefore abstain from the usage of “contraction” as it suggests an active process, commonly through actomyosin activity. We decided to use the term “deflation” to describe the process in which the sponge is reducing its body volume, irrespective of the individual tissue changes. The colloquial usage of “sneezing” fits well and is warranted. In contrast, we call the expansion of body volume “inflation”. As noted by Weissenfels, Leys, Nickel and Nichols, incurrent canal systems together with the epithelial tent and the excurrent canal system behave antagonistically (2). During deflation, the incurrent canal system and the tent are “collapsing” (also (12)) while the excurrent system is “expanding”. Only on the level of individual cells (3), such as pinacocytes, we decided to utilize “contraction” vs. “relaxation” because it directly relates to the cellular process of actomyosin contraction vs. relaxation. Since contraction equals increased tension in actomyosin stress fibers, we also use the phrase “tension increase” for contraction and opposing “tension decrease” for relaxation. Notably actomyosin contraction can be isometric, i.e. involving tension increase without length changes, which is assumed here for the epithelial tent during the re-establishment of the inflated default state.

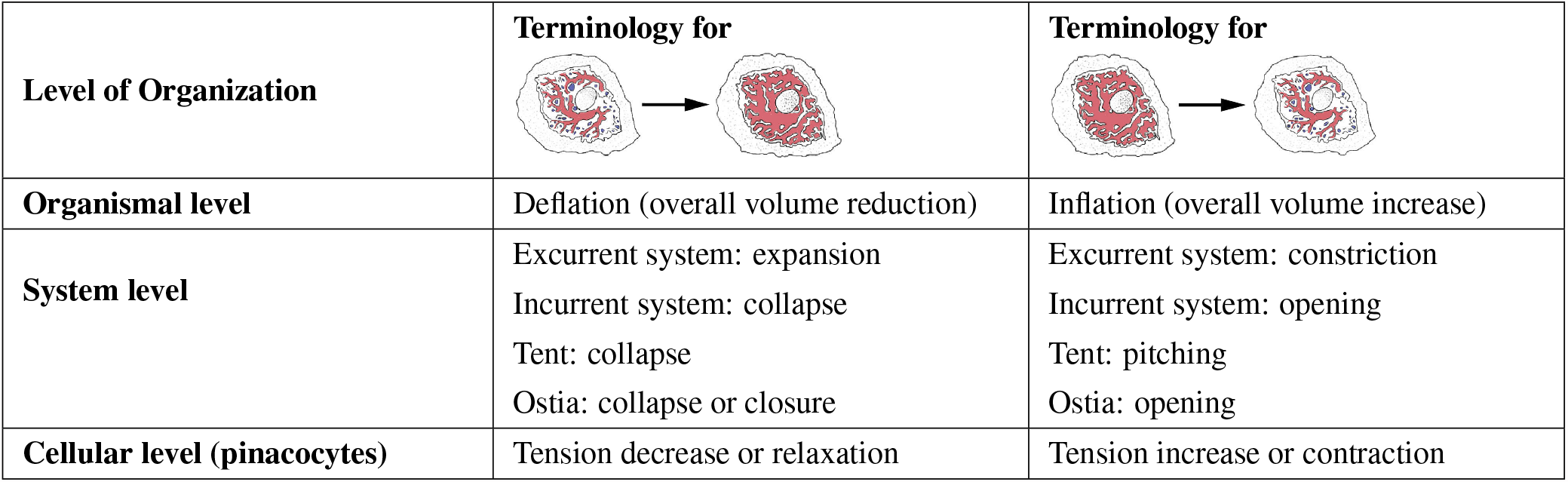

| Level of Organization | Terminology for | Terminology for |
| --- | --- | --- |
| Organismal level | Deflation (overall volume reduction) | Inflation (overall volume increase) |
| System level | Excurrent system: expansion<br>Incurrent system: collapse<br>Tent: collapse<br>Ostia: collapse or closure | Excurrent system: constriction<br>Incurrent system: opening<br>Tent: pitching<br>Ostia: opening |
| Cellular level (pinacocytes) | Tension decrease or relaxation | Tension increase or contraction |

## Notes

https://zenodo.org/record/8116913

https://git.embl.de/grp-arendt/spongeprot

### Competing Interest Statement

The authors have declared no competing interest.

## Bibliography

1. Thibaut Brunet and Nicole King. The origin of animal multicellularity and cell differentiation. Dev. Cell, 43(2):124–140, October 2017.

2. T L Simpson. The Cell Biology of Sponges. Springer Science & Business Media, December 2012.

3. Seyed Saeed Asadzadeh, Poul S Larsen, Hans Ulrik Riisgård, and Jens H Walther. Hydrodynamics of the leucon sponge pump. J. R. Soc. Interface, 16(150):20180630, January 2019.

4. Josephine Goldstein, Nicklas Bisbo, Peter Funch, and Hans Ulrik Riisgård. Contraction-Expansion and the effects on the aquiferous system in the demosponge halichondria panicea. Front. Mar. Sci., 7, February 2020.

5. H M Reiswig. In situ pumping activities of tropical demospongiae. Mar. Biol., 9(1):38–50, April 1971.

6. Niklas A Kornder, Yuki Esser, Daniel Stoupin, Sally P Leys, Benjamin Mueller, Mark J A Vermeij, Jef Huisman, and Jasper M de Goeij. Sponges sneeze mucus to shed particulate waste from their seawater inlet pores. Curr. Biol., 32(17):3855–3861.e3, September 2022.

7. Lars Kumala and Donald Eugene Canfield. Contraction dynamics and respiration of small Single-Osculum explants of the demosponge halichondria panicea. Front. Mar. Sci., 5, November 2018.

8. Glen R D Elliott and Sally P Leys. Coordinated contractions effectively expel water from the aquiferous system of a freshwater sponge. J. Exp. Biol., 210(Pt 21):3736–3748, November 2007.

9. Glen R D Elliott and Sally P Leys. Evidence for glutamate, GABA and NO in coordinating behaviour in the sponge, ephydatia muelleri (demospongiae, spongillidae). J. Exp. Biol., 213(Pt 13):2310–2321, July 2010.

10. Kornelia Ellwanger and Michael Nickel. Neuroactive substances specifically modulate rhythmic body contractions in the nerveless metazoon tethya wilhelma (demospongiae, porifera). Front. Zool., 3:7, April 2006.

11. Kornelia Ellwanger, Andre Eich, and Michael Nickel. GABA and glutamate specifically induce contractions in the sponge tethya wilhelma. J. Comp. Physiol. A Neuroethol. Sens. Neural Behav. Physiol., 193(1):1–11, January 2007.

12. J Colgren and S A Nichols. MRTF specifies a muscle-like contractile module in porifera. Nat. Commun., 13(1):4134, July 2022.

13. Jacob M Musser, Klaske J Schippers, Michael Nickel, Giulia Mizzon, Andrea B Kohn, Constantin Pape, Paolo Ronchi, Nikolaos Papadopoulos, Alexander J Tarashansky, Jörg U Hammel, Florian Wolf, Cong Liang, Ana Hernández-Plaza, Carlos P Cantalapiedra, Kaia Achim, Nicole L Schieber, Leslie Pan, Fabian Ruperti, Warren R Francis, Sergio Vargas, Svenja Kling, Maike Renkert, Maxim Polikarpov, Gleb Bourenkov, Roberto Feuda, Imre Gaspar, Pawel Burkhardt, Bo Wang, Peer Bork, Martin Beck, Thomas R Schneider, Anna Kreshuk, Gert Wörheide, Jaime Huerta-Cepas, Yannick Schwab, Leonid L Moroz, and Detlev Arendt. Profiling cellular diversity in sponges informs animal cell type and nervous system evolution. Science, 374(6568):717–723, November 2021.

14. Danielle A Ludeman, Nathan Farrar, Ana Riesgo, Jordi Paps, and Sally P Leys. Evolutionary origins of sensation in metazoans: functional evidence for a new sensory organ in sponges. BMC Evol. Biol., 14:3, January 2014.

15. Michael Nickel. Kinetics and rhythm of body contractions in the sponge tethya wilhelma (porifera: Demospongiae). J. Exp. Biol., 207(Pt 26):4515–4524, December 2004.

16. Michael Nickel. Like a ‘rolling stone’: quantitative analysis of the body movement and skeletal dynamics of the sponge Tethya wilhelma, 2006.

17. Michael Nickel, Corina Scheer, Jörg U Hammel, Julia Herzen, and Felix Beckmann. The contractile sponge epithelium sensu lato–body contraction of the demosponge tethya wilhelma is mediated by the pinacoderm. J. Exp. Biol., 214(Pt 10):1692–1698, May 2011.

18. Arnau Sebé-Pedrós, Elad Chomsky, Kevin Pang, David Lara-Astiaso, Federico Gaiti, Zohar Mukamel, Ido Amit, Andreas Hejnol, Bernard M Degnan, and Amos Tanay. Early metazoan cell type diversity and the evolution of multicellular gene regulation. Nat Ecol Evol, 2(7): 1176–1188, July 2018.

19. Jennyfer M Mitchell and Scott A Nichols. Diverse cell junctions with unique molecular composition in tissues of a sponge (porifera). Evodevo, 10:26, October 2019.

20. Josephine Goldstein, Hans Ulrik Riisgård, and Poul S Larsen. Exhalant jet speed of single-osculum explants of the demosponge halichondria panicea and basic properties of the sponge-pump. J. Exp. Mar. Bio. Ecol., 511:82–90, February 2019.

21. Sindhuja Sridharan, Alberto Hernandez-Armendariz, Nils Kurzawa, Clement M Potel, Dan-ish Memon, Pedro Beltrao, Marcus Bantscheff, Wolfgang Huber, Sara Cuylen-Haering, and Mikhail M Savitski. Systematic discovery of biomolecular condensate-specific protein phosphorylation. Nat. Chem. Biol., 18(10):1104–1114, October 2022.

22. Mikhail M Savitski, Friedrich B M Reinhard, Holger Franken, Thilo Werner, Maria Fälth Savitski, Dirk Eberhard, Daniel Martinez Molina, Rozbeh Jafari, Rebecca Bakszt Dovega, Susan Klaeger, Bernhard Kuster, Pär Nordlund, Marcus Bantscheff, and Gerard Drewes. Tracking cancer drugs in living cells by thermal profiling of the proteome. Science, 346 (6205):1255784, October 2014.

23. André Mateus, Johannes Hevler, Jacob Bobonis, Nils Kurzawa, Malay Shah, Karin Mitosch, Camille V Goemans, Dominic Helm, Frank Stein, Athanasios Typas, and Mikhail M Savitski. The functional proteome landscape of escherichia coli. Nature, 588(7838):473– 478, December 2020.

24. Anniek Stokkermans, Aditi Chakrabarti, Kaushikaram Subramanian, Ling Wang, Sifan Yin, Prachiti Moghe, Petrus Steenbergen, Gregor Mönke, Takashi Hiiragi, Robert Prevedel, L Mahadevan, and Aissam Ikmi. Muscular hydraulics drive larva-polyp morphogenesis. Curr. Biol., 32(21):4707–4718.e8, November 2022.

25. N Weissenfels. Condensation rhythm of fresh-water sponges (spongillidae, porifera). Eur. J. Cell Biol., 53(2):373–383, December 1990.

26. Thibaut Brunet, Antje Hl Fischer, Patrick Rh Steinmetz, Antonella Lauri, Paola Bertucci, and Detlev Arendt. The evolutionary origin of bilaterian smooth and striated myocytes. Elife, 5, December 2016.

27. Patrick R H Steinmetz, Johanna E M Kraus, Claire Larroux, Jörg U Hammel, Annette Amon-Hassenzahl, Evelyn Houliston, Gert Wörheide, Michael Nickel, Bernard M Degnan, and Ulrich Technau. Independent evolution of striated muscles in cnidarians and bilaterians. Nature, 487(7406):231–234, July 2012.

28. I F Ghalayini. Nitric oxide–cyclic GMP pathway with some emphasis on cavernosal contractility. Int. J. Impot. Res., 16(6):459–469, July 2004.

29. Kazuo Katoh, Yumiko Kano, and Yasuko Noda. Rho-associated kinase-dependent contraction of stress fibres and the organization of focal adhesions. J. R. Soc. Interface, 8(56):305–311, March 2011.

30. Wei Liu, Zhen-Tang Jing, Shu-Xiang Wu, Yun He, Yan-Ting Lin, Wan-Nan Chen, Xin-Jian Lin, and Xu Lin. A novel AKT activator, SC79, prevents acute hepatic failure induced by Fas-Mediated apoptosis of hepatocytes. Am. J. Pathol., 188(5):1171–1182, May 2018.

31. S Dimmeler, I Fleming, B Fisslthaler, C Hermann, R Busse, and A M Zeiher. Activation of nitric oxide synthase in endothelial cells by akt-dependent phosphorylation. Nature, 399 (6736):601–605, June 1999.

32. Anar Dossumbekova, Evgeny V Berdyshev, Irina Gorshkova, Zuohui Shao, Changqing Li, Phillip Long, Atul Joshi, Viswanathan Natarajan, and Terry L Vanden Hoek. Akt activates NOS3 and separately restores barrier integrity in H2O2-stressed human cardiac microvascular endothelium. Am. J. Physiol. Heart Circ. Physiol., 295(6):H2417–26, December 2008.

33. Telmo A Mejía-García, Camila C Portugal, Thaísa G Encarnação, Marco Antônio M Prado, and Roberto Paes-de Carvalho. Nitric oxide regulates AKT phosphorylation and nuclear translocation in cultured retinal cells. Cell. Signal., 25(12):2424–2439, December 2013.

34. N Schweighofer and G Ferriol. Diffusion of nitric oxide can facilitate cerebellar learning: A simulation study. Proc. Natl. Acad. Sci. U. S. A., 97(19):10661–10665, September 2000.

35. I V Turko, S A Ballard, S H Francis, and J D Corbin. Inhibition of cyclic GMP-binding cyclic GMP-specific phosphodiesterase (type 5) by sildenafil and related compounds. Mol. Pharmacol., 56(1):124–130, July 1999.

36. Bin-Nan Wu, Rong-Jyh Lin, Yi-Ching Lo, Kuo-Pyng Shen, Chao-Chuan Wang, Young-Tso Lin, and Ing-Jun Chen. KMUP-1, a xanthine derivative, induces relaxation of guinea-pig isolated trachea: the role of the epithelium, cyclic nucleotides and k+ channels. Br. J. Pharmacol., 142(7):1105–1114, August 2004.

37. Alejandro E Leroux, Jörg O Schulze, and Ricardo M Biondi. AGC kinases, mechanisms of regulation and innovative drug development. Semin. Cancer Biol., 48:1–17, February 2018.

38. Wei-Chiao Chiu, Jiun-Yang Chiang, and Fu-Tien Chiang. Small chemical compounds Y16 and rhosin can inhibit calcium sensitization pathway in vascular smooth muscle cells of spontaneously hypertensive rats. J. Formos. Med. Assoc., 120(10):1863–1868, October 2021.

39. Rita Rosenthal, Lars Choritz, Sebastian Schlott, Nikolaos E Bechrakis, Jan Jaroszewski, Michael Wiederholt, and Hagen Thieme. Effects of ML-7 and Y-27632 on carbachol- and endothelin-1-induced contraction of bovine trabecular meshwork. Exp. Eye Res., 80(6): 837–845, June 2005.

40. M S Goligorsky, H Abedi, E Noiri, A Takhtajan, S Lense, V Romanov, and I Zachary. Nitric oxide modulation of focal adhesions in endothelial cells. Am. J. Physiol., 276(6):C1271–81, June 1999.

41. L Atkinson, M Z Yusuf, A Aburima, Y Ahmed, S G Thomas, K M Naseem, and S D J Calaminus. Reversal of stress fibre formation by nitric oxide mediated RhoA inhibition leads to reduction in the height of preformed thrombi. Sci. Rep., 8(1):3032, February 2018.

42. Robert W Meech. Non-neural reflexes: sponges and the origins of behaviour. Curr. Biol., 18(2):R70–2, January 2008.

43. Elena Kassianidou, Jasmine H Hughes, and Sanjay Kumar. Activation of ROCK and MLCK tunes regional stress fiber formation and mechanics via preferential myosin light chain phosphorylation. Mol. Biol. Cell, 28(26):3832–3843, December 2017.

44. Ariel Livne and Benjamin Geiger. The inner workings of stress fibers-from contractile machinery to focal adhesions and back. J. Cell Sci., 129(7):1293–1304, 2016.

45. S Pemrick and A Weber. Mechanism of inhibition of relaxation by n-ethylmaleimide treatment of myosin. Biochemistry, 15(23):5193–5198, November 1976.

46. Marina Orman, Maya Landis, Aisha Oza, Deepika Nambiar, Joana Gjeci, Kristen Song, Vivian Huang, Amanda Klestzick, Carla Hachicho, Su Qing Liu, Judith M Kamm, Francesca Bartolini, Jean J Vadakkan, Christian M Rojas, and Christina L Vizcarra. Alterations to the broad-spectrum formin inhibitor SMIFH2 modulate potency but not specificity. Sci. Rep., 12(1):13520, August 2022.

47. Naomi Courtemanche. Mechanisms of formin-mediated actin assembly and dynamics. Biophys. Rev., 10(6):1553–1569, December 2018.

48. Fernando R Valencia, Eduardo Sandoval, Joy Du, Ernest Iu, Jian Liu, and Sergey V Plotnikov. Force-dependent activation of actin elongation factor mdia1 protects the cytoskeleton from mechanical damage and promotes stress fiber repair. Dev. Cell, 56(23):3288– 3302.e5, December 2021.

49. S P Leys and R W Meech. Physiology of coordination in sponges. Can. J. Zool., 84(2):288–306, February 2006.

50. Jörg U Hammel and Michael Nickel. A new flow-regulating cell type in the demosponge tethya wilhelma - functional cellular anatomy of a leuconoid canal system. PLoS One, 9 (11):e113153, November 2014.

51. E Lemichez, Y Wu, J P Sanchez, A Mettouchi, J Mathur, and N H Chua. Inactivation of AtRac1 by abscisic acid is essential for stomatal closure. Genes Dev., 15(14):1808–1816, July 2001.

52. Dong Qian, Zhe Zhang, Juanxia He, Pan Zhang, Xiaobin Ou, Tian Li, Lipan Niu, Qiong Nan, Yue Niu, Wenliang He, Lizhe An, Kun Jiang, and Yun Xiang. Arabidopsis ADF5 promotes stomatal closure by regulating actin cytoskeleton remodeling in response to ABA and drought stress. J. Exp. Bot., 70(2):435–446, January 2019.

53. Yihao Li, Xin Zhang, Yi Zhang, and Haiyun Ren. Controlling the gate: The functions of the cytoskeleton in stomatal movement. Front. Plant Sci., 13:849729, February 2022.

54. George R R Bell, Alan M Kuzirian, Stephen L Senft, Lydia M Mäthger, Trevor J Wardill, and Roger T Hanlon. Chromatophore radial muscle fibers anchor in flexible squid skin. Invertebr. Biol., 132(2):120–132, June 2013.

55. Poul Scheel Larsen and Hans Ulrik Riisgåd. The sponge pump. J. Theor. Biol., 168(1): 53–63, May 1994.

56. Clement M Potel, Miao-Hsia Lin, Albert J R Heck, and Simone Lemeer. Defeating major contaminants in fe3+-immobilized metal ion affinity chromatography (IMAC) phosphopeptide enrichment. Mol. Cell. Proteomics, 17(5):1028–1034, May 2018.

57. Clément M Potel, Nils Kurzawa, Isabelle Becher, Athanasios Typas, André Mateus, and Mikhail M Savitski. Impact of phosphorylation on thermal stability of proteins. Nat. Methods, 18(7):757–759, July 2021.

58. Cristina Viéitez, Bede P Busby, David Ochoa, André Mateus, Danish Memon, Marco Galardini, Umut Yildiz, Matteo Trovato, Areeb Jawed, Alexander G Geiger, and Others. High-throughput functional characterization of protein phosphorylation sites in yeast. Nat. Biotechnol., 40(3):382–390, 2022.

59. Holger Franken, Toby Mathieson, Dorothee Childs, Gavain M A Sweetman, Thilo Werner, Ina Tögel, Carola Doce, Stephan Gade, Marcus Bantscheff, Gerard Drewes, Friedrich B M Reinhard, Wolfgang Huber, and Mikhail M Savitski. Thermal proteome profiling for unbiased identification of direct and indirect drug targets using multiplexed quantitative mass spectrometry. Nat. Protoc., 10(10):1567–1593, October 2015.

60. André Mateus, Tomi A Määttä, and Mikhail M Savitski. Thermal proteome profiling: unbiased assessment of protein state through heat-induced stability changes. Proteome Sci., 15(1):13, 2016.

61. André Mateus, Nils Kurzawa, Isabelle Becher, Sindhuja Sridharan, Dominic Helm, Frank Stein, Athanasios Typas, and Mikhail M Savitski. Thermal proteome profiling for interrogating protein interactions. Mol. Syst. Biol., 16(3):e9232, March 2020.

62. Isabelle Becher, Amparo Andrés-Pons, Natalie Romanov, Frank Stein, Maike Schramm, Florence Baudin, Dominic Helm, Nils Kurzawa, André Mateus, Marie-Therese Mackmull, Athanasios Typas, Christoph W Müller, Peer Bork, Martin Beck, and Mikhail M Savitski. Pervasive protein thermal stability variation during the cell cycle. Cell, 173(6):1495–1507.e18, May 2018.

63. Chenwei Wang, Haodong Xu, Shaofeng Lin, Wankun Deng, Jiaqi Zhou, Ying Zhang, Ying Shi, D. Peng, and Yu Xue. GPS 5.0: An update on the prediction of kinase-specific phosphorylation sites in proteins. Genomics Proteomics Bioinformatics, 18(1):72–80, February 2020.

64. Poul M Bendix, Gijsje H Koenderink, Damien Cuvelier, Zvonimir Dogic, Bernard N Koele-man, William M Brieher, Christine M Field, L Mahadevan, and David A Weitz. A quantitative analysis of contractility in active cytoskeletal protein networks. Biophys. J., 94(8): 3126–3136, April 2008.

65. A E Carlsson. Contractile stress generation by actomyosin gels. Phys. Rev. E Stat. Nonlin. Soft Matter Phys., 74(5 Pt 1):051912, November 2006.

66. Muna Taha, Mohammed Aldirawi, Sigrid März, Jochen Seebach, Maria Odenthal-Schnittler, Olga Bondareva, Vesna Bojovic, Thomas Schmandra, Benedikt Wirth, Mag-dalena Mietkowska, Klemens Rottner, and Hans Schnittler. EPLIN-α and -β isoforms modulate endothelial cell dynamics through a spatiotemporally differentiated interaction with actin. Cell Rep., 29(4):1010–1026.e6, October 2019.

67. Bassam Janji, Adeline Giganti, Veerle De Corte, Marie Catillon, Erik Bruyneel, Delphine Lentz, Julie Plastino, Jan Gettemans, and Evelyne Friederich. Phosphorylation on ser5 increases the f-actin-binding activity of l-plastin and promotes its targeting to sites of actin assembly in cells. J. Cell Sci., 119(Pt 9):1947–1960, May 2006.

68. Natalya G Dulyaninova, Vladimir N Malashkevich, Steven C Almo, and Anne R Bresnick. Regulation of myosin-IIA assembly and mts1 binding by heavy chain phosphorylation. Biochemistry, 44(18):6867–6876, May 2005.

69. Mei-Ying Han, Hidetaka Kosako, Toshiki Watanabe, and Seisuke Hattori. Extracellular signal-regulated kinase/mitogen-activated protein kinase regulates actin organization and cell motility by phosphorylating the actin cross-linking protein EPLIN. Mol. Cell. Biol., 27 (23):8190–8204, December 2007.

70. Timothy R Baffi, Gema Lordén, Jacob M Wozniak, Andreas Feichtner, Wayland Yeung, Alexandr P Kornev, Charles C King, Jason C Del Rio, Ameya J Limaye, Julius Bogomolovas, Christine M Gould, Ju Chen, Eileen J Kennedy, Natarajan Kannan, David J Gonzalez, Eduard Stefan, Susan S Taylor, and Alexandra C Newton. mTORC2 controls the activity of PKC and akt by phosphorylating a conserved TOR interaction motif. Sci. Signal., 14(678), April 2021.

71. Richard Seonghun Nho and Polla Hergert. FoxO3a and disease progression. World J. Biol. Chem., 5(3):346–354, August 2014.

72. Kebin Zhang, Xiaoqin Guo, Hui Yan, Yuxin Wu, Quan Pan, James Zheng Shen, Xiaopeng Li, Yunmei Chen, Ling Li, Yajuan Qi, Zihui Xu, Wei Xie, Weiping Zhang, David Threadgill, Ling He, Daniel Villarreal, Yuxiang Sun, Morris F White, Hongting Zheng, and Shaodong Guo. Phosphorylation of forkhead protein FoxO1 at S253 regulates glucose homeostasis in mice. Endocrinology, 160(5):1333–1347, May 2019.

73. A Brunet, A Bonni, M J Zigmond, M Z Lin, P Juo, L S Hu, M J Anderson, K C Arden, J Blenis, and M E Greenberg. Akt promotes cell survival by phosphorylating and inhibiting a forkhead transcription factor. Cell, 96(6):857–868, March 1999.

74. W Wimmer, S Perovic, M Kruse, H C Schröder, A Krasko, R Batel, and W E Müller. Origin of the integrin-mediated signal transduction. functional studies with cell cultures from the sponge suberites domuncula. Eur. J. Biochem., 260(1):156–165, February 1999.

75. John C Salerno, Dipak K Ghosh, Raj Razdan, Katy A Helms, Christopher C Brown, Jonathan L McMurry, Emily A Rye, and Carol A Chrestensen. Endothelial nitric oxide synthase is regulated by ERK phosphorylation at ser602. Biosci. Rep., 34(5), September 2014.

76. Chaohong Li, Yanhua Hu, Manuel Mayr, and Qingbo Xu. Cyclic strain stress-induced mitogen-activated protein kinase (MAPK) phosphatase 1 expression in vascular smooth muscle cells is regulated by Ras/Rac-MAPK pathways*. J. Biol. Chem., 274(36):25273–25280, September 1999.

77. Felix Teufel, José Juan Almagro Armenteros, Alexander Rosenberg Johansen, Magnús Halldór Gíslason, Silas Irby Pihl, Konstantinos D Tsirigos, Ole Winther, Søren Brunak, Gunnar von Heijne, and Henrik Nielsen. SignalP 6.0 predicts all five types of signal peptides using protein language models. Nat. Biotechnol., 40(7):1023–1025, July 2022.

78. Fabian Ruperti. Go enrichment and signal sequence analysis. https://git.embl.de/grp-arendt/spongeprot/-/blob/main/Data%20analysis/Proteomics/proteomics_GO_enrichment.ipynb, 2023. Accessed: 2023-07.

79. Milot Mirdita, Konstantin Schütze, Yoshitaka Moriwaki, Lim Heo, Sergey Ovchinnikov, and Martin Steinegger. ColabFold: making protein folding accessible to all. Nat. Methods, 19 (6):679–682, June 2022.

80. Fabian Ruperti, Nikolaos Papadopoulos, Jacob M Musser, Milot Mirdita, Martin Steinegger, and Detlev Arendt. Cross-phyla protein annotation by structural prediction and alignment. Genome Biol., 24(1):113, May 2023.

81. Jia-Shin Lin, Shuo-Kang Lee, Yeh Chen, Wei-De Lin, and Chao-Hung Kao. Purification and characterization of a novel extracellular tripeptidyl peptidase from rhizopus oligosporus. J. Agric. Food Chem., 59(20):11330–11337, October 2011.

82. Eva Vidak, Urban Javoršek, Matej Vizovišek, and Boris Turk. Cysteine cathepsins and their extracellular roles: Shaping the microenvironment. Cells, 8(3), March 2019.

83. Peter J Lyons, Myrasol B Callaway, and Lloyd D Fricker. Characterization of carboxypeptidase a6, an extracellular matrix peptidase. J. Biol. Chem., 283(11):7054–7063, March 2008.

84. Urska Repnik, Amanda E Starr, Christopher M Overall, and Boris Turk. Cysteine cathepsins activate ELR chemokines and inactivate Non-ELR chemokines. J. Biol. Chem., 290 (22):13800–13811, May 2015.

85. Mare Cudic and Gregg B Fields. Extracellular proteases as targets for drug development. Curr. Protein Pept. Sci., 10(4):297–307, August 2009.

86. Ivica Letunic, Tobias Doerks, and Peer Bork. SMART 7: recent updates to the protein domain annotation resource. Nucleic Acids Res., 40(Database issue):D302–5, January 2012.

87. Luis Alfonso Yañez-Guerra, Daniel Thiel, and Gáspár Jékely. Premetazoan origin of neuropeptide signaling. Mol. Biol. Evol., 39(4), April 2022.

88. Swetha Mohan, Paul J Sampognaro, Andrea R Argouarch, Jason C Maynard, Mackenzie Welch, Anand Patwardhan, Emma C Courtney, Jiasheng Zhang, Amanda Mason, Kathy H Li, Eric J Huang, William W Seeley, Bruce L Miller, Alma Burlingame, Mathew P Jacobson, and Aimee W Kao. Processing of progranulin into granulins involves multiple lysosomal proteases and is affected in frontotemporal lobar degeneration. Mol. Neurodegener., 16 (1):51, August 2021.

89. Sandra Almeida, Lijuan Zhou, and Fen-Biao Gao. Progranulin, a glycoprotein deficient in frontotemporal dementia, is a novel substrate of several protein disulfide isomerase family proteins. PLoS One, 6(10):e26454, October 2011.

90. Laurent Meunier, Young-Kwang Usherwood, Kyung Tae Chung, and Linda M Hendershot. A subset of chaperones and folding enzymes form multiprotein complexes in endoplasmic reticulum to bind nascent proteins. Mol. Biol. Cell, 13(12):4456–4469, December 2002.

91. Hwan-Jin Hwang, Tae Woo Jung, Ho Cheol Hong, Hae Yoon Choi, Ji-A Seo, Sin Gon Kim, Nan Hee Kim, Kyung Mook Choi, Dong Seop Choi, Sei Hyun Baik, and Hye Jin Yoo. Progranulin protects vascular endothelium against atherosclerotic inflammatory reaction via Akt/eNOS and nuclear factor-κB pathways. PLoS One, 8(9):e76679, September 2013.

92. Shi-Qiong Xu, Simone Buraschi, Ryuta Tanimoto, Manuela Stefanello, Antonino Belfiore, Renato V Iozzo, and Andrea Morrione. Analysis of Progranulin-Mediated akt and MAPK activation. Methods Mol. Biol., 1806:121–130, 2018.

93. H Lue, M Thiele, J Franz, E Dahl, S Speckgens, L Leng, G Fingerle-Rowson, R Bucala, B Lüscher, and J Bernhagen. Macrophage migration inhibitory factor (MIF) promotes cell survival by activation of the akt pathway and role for CSN5/JAB1 in the control of autocrine MIF activity. Oncogene, 26(35):5046–5059, August 2007.

94. Hongqi Lue, Aphrodite Kapurniotu, Günter Fingerle-Rowson, Thierry Roger, Lin Leng, Michael Thiele, Thierry Calandra, Richard Bucala, and Jürgen Bernhagen. Rapid and transient activation of the ERK MAPK signalling pathway by macrophage migration inhibitory factor (MIF) and dependence on JAB1/CSN5 and src kinase activity. Cell. Signal., 18(5):688–703, May 2006.

95. Rachael A Harrison and Colin Sumners. Redox regulation of macrophage migration inhibitory factor expression in rat neurons. Biochem. Biophys. Res. Commun., 390(1):171– 175, December 2009.

96. Manish Mittal, Mohammad Rizwan Siddiqui, Khiem Tran, Sekhar P Reddy, and Asrar B Malik. Reactive oxygen species in inflammation and tissue injury. Antioxid. Redox Signal., 20(7):1126–1167, March 2014.

97. Adelheid Weidinger and Andrey V Kozlov. Biological activities of reactive oxygen and nitrogen species: Oxidative stress versus signal transduction. Biomolecules, 5(2):472– 484, April 2015.

98. Anna Podgórska, Maria Burian, and Bozena Szal. Extra-Cellular but Extra-Ordinarily im-portant for cells: Apoplastic reactive oxygen species metabolism. Front. Plant Sci., 8:1353, August 2017.

99. Guijie Ren, Zhongcai Ma, Maria Hui, Lili C Kudo, Koon-Sea Hui, and Stanislav L Karsten. Cu, zn-superoxide dismutase 1 (SOD1) is a novel target of puromycin-sensitive aminopeptidase (PSA/NPEPPS): PSA/NPEPPS is a possible modifier of amyotrophic lateral sclerosis. Mol. Neurodegener., 6:29, May 2011.

100. O I Aruoma, B Halliwell, B M Hoey, and J Butler. The antioxidant action of taurine, hypotaurine and their metabolic precursors. Biochem. J, 256(1):251–255, November 1988.

101. Italo Rodrigo Calori, Luiza Araújo Gusmão, and Antonio Claudio Tedesco. B6 vitamers as generators and scavengers of reactive oxygen species. Journal of Photochemistry and Photobiology, 7:100041, September 2021.

102. Shoshana M Bartell, Ha-Neui Kim, Elena Ambrogini, Li Han, Srividhya Iyer, S Serra Ucer, Peter Rabinovitch, Robert L Jilka, Robert S Weinstein, Haibo Zhao, Charles A O’Brien, Stavros C Manolagas, and Maria Almeida. FoxO proteins restrain osteoclastogenesis and bone resorption by attenuating H2O2 accumulation. Nat. Commun., 5:3773, April 2014.

103. Yan Tao, Shanhui Liu, Jianzhong Lu, Shengjun Fu, Lanlan Li, Jing Zhang, Zhiping Wang, and Mei Hong. FOXO3a-ROS pathway is involved in androgen-induced proliferation of prostate cancer cell. BMC Urol., 22(1):1–10, April 2022.

104. Safak Yalcin, Dragan Marinkovic, Sathish Kumar Mungamuri, Xin Zhang, Wei Tong, Rani Sellers, and Saghi Ghaffari. ROS-mediated amplification of AKT/mTOR signalling pathway leads to myeloproliferative syndrome in foxo3(-/-) mice. EMBO J., 29(24):4118–4131, December 2010.

105. Emile A Kraus, Lauren E Mellenthin, Sara A Siwiecki, Dawei Song, Jing Yan, Paul A Janmey, and Alison M Sweeney. Rheology of marine sponges reveals anisotropic mechanics and tuned dynamics. J. R. Soc. Interface, 19(195):20220476, October 2022.

106. K C Holmes, R R Schröder, H L Sweeney, and Anne Houdusse. The structure of the rigor complex and its implications for the power stroke. Philos. Trans. R. Soc. Lond. B Biol. Sci., 359(1452):1819–1828, December 2004.

107. Takashi Fujii and Keiichi Namba. Structure of actomyosin rigour complex at 5.2 å resolution and insights into the ATPase cycle mechanism. Nat. Commun., 8:13969, January 2017.

108. Oleg Andruchov, Olena Andruchova, and Stefan Galler. The catch state of mollusc catch muscle is established during activation: experiments on skinned fibre preparations of the anterior byssus retractor muscle of mytilus edulis l. using the myosin inhibitors orthovanadate and blebbistatin. J. Exp. Biol., 209(Pt 21):4319–4328, November 2006.

109. Sabrena Noria, Feng Xu, Shannon McCue, Mara Jones, Avrum I Gotlieb, and B Lowell Langille. Assembly and reorientation of stress fibers drives morphological changes to endothelial cells exposed to shear stress. Am. J. Pathol., 164(4):1211–1223, April 2004.

110. Jaime Millán, Robert J Cain, Natalia Reglero-Real, Carolina Bigarella, Beatriz Marcos-Ramiro, Laura Fernández-Martín, Isabel Correas, and Anne J Ridley. Adherens junctions connect stress fibres between adjacent endothelial cells. BMC Biol., 8:11, February 2010.

111. Mack H Wu. Endothelial focal adhesions and barrier function. J. Physiol., 569(Pt 2):359– 366, December 2005.

112. George T McNAIR. MOTOR REACTIONS OF THE FRESH-WATER SPONGE, EPHYDA-TIA FLUVIATILIS. Biol. Bull., 44(4):153–166, April 1923.

113. Sally P Leys. Elements of a ‘nervous system’in sponges. J. Exp. Biol., 218(4):581–591, 2015.

114. R H Emson. The reactions of the sponge cliona celata to applied stimuli. Comp. Biochem. Physiol., 18(4):805–827, August 1966.

115. C C Wu and D F Bohr. Mechanisms of calcium relaxation of vascular smooth muscle. Am. J. Physiol., 261(5 Pt 2):H1411–6, November 1991.

116. Denis Martinvalet and Michael Walch. Editorial: The role of reactive oxygen species in protective immunity. Front. Immunol., 12:832946, 2021.

117. Thierry Calandra and Thierry Roger. Macrophage migration inhibitory factor: a regulator of innate immunity. Nat. Rev. Immunol., 3(10):791–800, October 2003.

118. C Bogdan, M Röllinghoff, and A Diefenbach. The role of nitric oxide in innate immunity. Immunol. Rev., 173:17–26, February 2000.

119. Ana Riesgo, Nadia Santodomingo, Vasiliki Koutsouveli, Lars Kumala, Michelle M Leger, Sally P Leys, and Peter Funch. Molecular machineries of ciliogenesis, cell survival, and vasculogenesis are differentially expressed during regeneration in explants of the demo-sponge halichondria panicea. BMC Genomics, 23(1):858, December 2022.

120. Lucía Pita, Marc P Hoeppner, Marta Ribes, and Ute Hentschel. Differential expression of immune receptors in two marine sponges upon exposure to microbial-associated molecular patterns. Sci. Rep., 8(1):16081, October 2018.

121. Yu-Chen Wu, Soeren Franzenburg, Marta Ribes, and Lucía Pita. Wounding response in porifera (sponges) activates ancestral signaling cascades involved in animal healing, regeneration, and cancer. Sci. Rep., 12(1):1307, January 2022.

122. Alpha S Yap, Kinga Duszyc, and Virgile Viasnoff. Mechanosensing and mechanotransduction at Cell-Cell junctions. Cold Spring Harb. Perspect. Biol., 10(8), August 2018.

123. Etienne Roux, Pauline Bougaran, Pascale Dufourcq, and Thierry Couffinhal. Fluid shear stress sensing by the endothelial layer. Front. Physiol., 11:861, July 2020.

124. Georgina G J Hazell, Alasdair M G Peachey, Jack E Teasdale, Graciela B Sala-Newby, Gianni D Angelini, Andrew C Newby, and Stephen J White. PI16 is a shear stress and inflammation-regulated inhibitor of MMP2. Sci. Rep., 6:39553, December 2016.

125. Kristopher S Cunningham and Avrum I Gotlieb. The role of shear stress in the pathogenesis of atherosclerosis. Lab. Invest., 85(1):9–23, January 2005.

126. Kazuo Katoh, Yumiko Kano, and Shigeo Ookawara. Role of stress fibers and focal adhesions as a mediator for mechano-signal transduction in endothelial cells in situ. Vasc. Health Risk Manag., 4(6):1273–1282, 2008.

127. Tung-Lin Yang, Pei-Ling Lee, Ding-Yu Lee, Wei-Li Wang, Shu-Yi Wei, Chih-I Lee, and Jeng-Jiann Chiu. Differential regulations of fibronectin and laminin in smad2 activation in vascular endothelial cells in response to disturbed flow. J. Biomed. Sci., 25(1):1, January 2018.

128. Narayan Dahal, Sabita Sharma, Binh Phan, Annie Eis, and Ionel Popa. Mechanical regulation of talin through binding and history-dependent unfolding. Sci Adv, 8(28):eabl7719, July 2022.

129. Congzhen Qiao, Shengdi Li, Haocheng Lu, Fan Meng, Yanbo Fan, Yanhong Guo, Y Eugene Chen, and Jifeng Zhang. Laminar flow attenuates macrophage migration inhibitory factor expression in endothelial cells. Sci. Rep., 8(1):2360, February 2018.

130. H Matsushita, K H Lee, and P S Tsao. Cyclic strain induces reactive oxygen species production via an endothelial NAD(P)H oxidase. J. Cell. Biochem. Suppl., Suppl 36:99–106, 2001.

131. Hua Cai, Joseph S McNally, Martina Weber, and David G Harrison. Oscillatory shear stress upregulation of endothelial nitric oxide synthase requires intracellular hydrogen peroxide and CaMKII. J. Mol. Cell. Cardiol., 37(1):121–125, July 2004.

132. Danielle A Ludeman, Matthew A Reidenbach, and Sally P Leys. The energetic cost of filtration by demosponges and their behavioural response to ambient currents. J. Exp. Biol., 220(Pt 6):995–1007, March 2017.

133. María López-Acosta, Clémence Potel, Morgane Gallinari, Fiz F Pérez, and Aude Leynaert. Nudibranch predation boosts sponge silicon cycling. Sci. Rep., 13(1):1178, January 2023.

134. Jochen Gugel. Life cycles and ecological interactions of freshwater sponges (porifera, spongillidae) in the river rhine in germany. Limnologica, 31(3):185–198, September 2001.

135. I Tjensvoll, T Kutti, J H Fosså, and R J Bannister. Rapid respiratory responses of the deep-water sponge geodia barretti exposed to suspended sediments. Aquat. Biol., 19(1): 65–73, September 2013.

136. M A Koehl. Ecological biomechanics of benthic organisms: life history, mechanical design and temporal patterns of mechanical stress. J. Exp. Biol., 202(Pt 23):3469–3476, December 1999.

137. Lara Schmittmann, Sören Franzenburg, and Lucía Pita. Individuality in the immune repertoire and induced response of the sponge halichondria panicea. Front. Immunol., 12: 689051, June 2021.

138. Olga Yu Gasheva, David C Zawieja, and Anatoliy A Gashev. Contraction-initiated NO-dependent lymphatic relaxation: a self-regulatory mechanism in rat thoracic duct. J. Physiol., 575(Pt 3):821–832, September 2006.

139. Amélie Sabine, Esther Bovay, Cansaran Saygili Demir, Wataru Kimura, Muriel Jaquet, Yan Agalarov, Nadine Zangger, Joshua P Scallan, Werner Graber, Elgin Gulpinar, Brenda R Kwak, Taija Mäkinen, Inés Martinez-Corral, Sagrario Ortega, Mauro Delorenzi, Friedemann Kiefer, Michael J Davis, Valentin Djonov, Naoyuki Miura, and Tatiana V Petrova. FOXC2 and fluid shear stress stabilize postnatal lymphatic vasculature. J. Clin. Invest., 125(10):3861–3877, October 2015.

140. Joshua P Scallan, Luz A Knauer, Huayan Hou, Jorge A Castorena-Gonzalez, Michael J Davis, and Ying Yang. Foxo1 deletion promotes the growth of new lymphatic valves. J. Clin. Invest., 131(14), July 2021.

141. J I Gillespie, M Markerink-van Ittersum, and J de Vente. Expression of neuronal nitric oxide synthase (nNOS) and nitric-oxide-induced changes in cGMP in the urothelial layer of the guinea pig bladder. Cell Tissue Res., 321(3):341–351, September 2005.

142. Lynn Stothers, Ismail Laher, and George T Christ. A review of the l-arginine - nitric oxide - guanylate cyclase pathway as a mediator of lower urinary tract physiology and symptoms. Can. J. Urol., 10(5):1971–1980, October 2003.

143. C Pinna, I Eberini, L Puglisi, and G Burnstock. Presence of constitutive endothelial nitric oxide synthase immunoreactivity in urothelial cells of hamster proximal urethra. Eur. J. Pharmacol., 367(1):85–89, February 1999.

144. Carsten Skurk, Yasuhiro Izumiya, Henrike Maatz, Peter Razeghi, Ichiro Shiojima, Marco Sandri, Kaori Sato, Ling Zeng, Stephan Schiekofer, David Pimentel, Stewart Lecker, Heinrich Taegtmeyer, Alfred L Goldberg, and Kenneth Walsh. The FOXO3a transcription factor regulates cardiac myocyte size downstream of AKT signaling*. J. Biol. Chem., 280(21): 20814–20823, May 2005.

145. Geng Bin, Zhang Bo, Wang Jing, Jiang Jin, Tan Xiaoyi, Chen Cong, An Liping, Ma Jinglin, Wang Cuifang, Chen Yonggang, and Xia Yayi. Fluid shear stress suppresses TNF-α-induced apoptosis in MC3T3-E1 cells: Involvement of ERK5-AKT-FoxO3a-Bim/FasL signaling pathways. Exp. Cell Res., 343(2):208–217, May 2016.

146. Frederik Seiler, Jan Hellberg, Philipp M Lepper, Andreas Kamyschnikow, Christian Herr, Markus Bischoff, Frank Langer, Hans-Joachim Schäfers, Frank Lammert, Michael D Menger, Robert Bals, and Christoph Beisswenger. FOXO transcription factors regulate innate immune mechanisms in respiratory epithelial cells. J. Immunol., 190(4):1603–1613, February 2013.

147. Qian Zhong, Yixin Liu, Michele Ramos Correa, Crystal Nicole Marconett, Parviz Minoo, Changgong Li, David K Ann, Beiyun Zhou, and Zea Borok. FOXO1 couples KGF and PI-3K/AKT signaling to NKX2.1-Regulated differentiation of alveolar epithelial cells. Cells, 11(7), March 2022.

148. Y Lu, L Parkyn, L E Otterbein, Y Kureishi, K Walsh, A Ray, and P Ray. Activated akt protects the lung from oxidant-induced injury and delays death of mice. J. Exp. Med., 193 (4):545–549, February 2001.

149. Indiwari Gopallawa, Li Eon Kuek, Nithin D Adappa, James N Palmer, and Robert J Lee. Small-molecule akt-activation in airway cells induces NO production and reduces IL-8 transcription through nrf-2. Respir. Res., 22(1):267, October 2021.

150. Diane Bridge, Alexander G Theofiles, Rebecca L Holler, Emily Marcinkevicius, Robert E Steele, and Daniel E Martínez. FoxO and stress responses in the cnidarian hydra vulgaris. PLoS One, 5(7):e11686, July 2010.

151. Casey W Dunn, Sally P Leys, and Steven H D Haddock. The hidden biology of sponges and ctenophores. Trends Ecol. Evol., 30(5):282–291, May 2015.

152. Emrah Eroglu, Benjamin Gottschalk, Suphachai Charoensin, Sandra Blass, Helmut Bischof, Rene Rost, Corina T Madreiter-Sokolowski, Brigitte Pelzmann, Eva Bernhart, Wolfgang Sattler, Seth Hallström, Tadeusz Malinski, Markus Waldeck-Weiermair, Wolfgang F Graier, and Roland Malli. Development of novel FP-based probes for live-cell imaging of nitric oxide dynamics. Nat. Commun., 7:10623, February 2016.

153. Yan Zhang, Márton Rózsa, Yajie Liang, Daniel Bushey, Ziqiang Wei, Jihong Zheng, Daniel Reep, Gerard Joey Broussard, Arthur Tsang, Getahun Tsegaye, Sujatha Narayan, Christopher J Obara, Jing-Xuan Lim, Ronak Patel, Rongwei Zhang, Misha B Ahrens, Glenn C Turner, Samuel S-H Wang, Wyatt L Korff, Eric R Schreiter, Karel Svoboda, Jeremy P Hasseman, Ilya Kolb, and Loren L Looger. Fast and sensitive GCaMP calcium indicators for imaging neural populations. Nature, 615(7954):884–891, March 2023.

154. Yasset Perez-Riverol, Jingwen Bai, Chakradhar Bandla, David García-Seisdedos, Suresh Hewapathirana, Selvakumar Kamatchinathan, Deepti J Kundu, Ananth Prakash, Anika Frericks-Zipper, Martin Eisenacher, Mathias Walzer, Shengbo Wang, Alvis Brazma, and Juan Antonio Vizcaíno. The PRIDE database resources in 2022: a hub for mass spectrometry-based proteomics evidences. Nucleic Acids Res., 50(D1):D543–D552, January 2022.

155. Kenneth Haug, Keeva Cochrane, Venkata Chandrasekhar Nainala, Mark Williams, Jiakang Chang, Kalai Vanii Jayaseelan, and Claire O’Donovan. MetaboLights: a resource evolving in response to the needs of its scientific community. Nucleic Acids Res., 48(D1):D440– D444, January 2020.

156. Matthew C Chambers, Brendan Maclean, Robert Burke, Dario Amodei, Daniel L Ruderman, Steffen Neumann, Laurent Gatto, Bernd Fischer, Brian Pratt, Jarrett Egertson, Katherine Hoff, Darren Kessner, Natalie Tasman, Nicholas Shulman, Barbara Frewen, Tahmina A Baker, Mi-Youn Brusniak, Christopher Paulse, David Creasy, Lisa Flashner, Kian Kani, Chris Moulding, Sean L Seymour, Lydia M Nuwaysir, Brent Lefebvre, Frank Kuhlmann, Joe Roark, Paape Rainer, Suckau Detlev, Tina Hemenway, Andreas Huhmer, James Langridge, Brian Connolly, Trey Chadick, Krisztina Holly, Josh Eckels, Eric W Deutsch, Robert L Moritz, Jonathan E Katz, David B Agus, Michael MacCoss, David L Tabb, and Parag Mallick. A cross-platform toolkit for mass spectrometry and proteomics. Nat. Biotechnol., 30(10):918–920, October 2012.

157. Andy T Kong, Felipe V Leprevost, Dmitry M Avtonomov, Dattatreya Mellacheruvu, and Alexey I Nesvizhskii. MSFragger: ultrafast and comprehensive peptide identification in mass spectrometry-based proteomics. Nat. Methods, 14(5):513–520, May 2017.

158. R Development Core Team and Others. R: A language and environment for statistical computing. R foundation for statistical computing, vienna, austria. ISBN 3-900051-07-0, 2008.

159. Lam-Tung Nguyen, Heiko A Schmidt, Arndt von Haeseler, and Bui Quang Minh. IQ-TREE: a fast and effective stochastic algorithm for estimating maximum-likelihood phylogenies. Mol. Biol. Evol., 32(1):268–274, January 2015.

160. Martin Steinegger and Johannes Söding. MMseqs2 enables sensitive protein sequence searching for the analysis of massive data sets. Nat. Biotechnol., 35(11):1026–1028, November 2017.

161. Michel van Kempen, Stephanie S Kim, Charlotte Tumescheit, Milot Mirdita, Jeongjae Lee, Cameron L M Gilchrist, Johannes Söding, and Martin Steinegger. Fast and accurate protein structure search with foldseek. Nat. Biotechnol., May 2023.

162. Kazutaka Katoh and Daron M Standley. MAFFT multiple sequence alignment software version 7: improvements in performance and usability. Mol. Biol. Evol., 30(4):772–780, April 2013.

163. Andrew M Waterhouse, James B Procter, David M A Martin, Michèle Clamp, and Geoffrey J Barton. Jalview version 2—a multiple sequence alignment editor and analysis workbench. Bioinformatics, 25(9):1189–1191, January 2009.

164. Ivica Letunic and Peer Bork. Interactive tree of life (iTOL) v5: an online tool for phylogenetic tree display and annotation. Nucleic Acids Res., 49(W1):W293–W296, July 2021.

165. D V Klopfenstein, Liangsheng Zhang, Brent S Pedersen, Fidel Ramírez, Alex Warwick Vesztrocy, Aurélien Naldi, Christopher J Mungall, Jeffrey M Yunes, Olga Botvinnik, Mark Weigel, Will Dampier, Christophe Dessimoz, Patrick Flick, and Haibao Tang. GOA-TOOLS: A python library for gene ontology analyses. Sci. Rep., 8(1):10872, July 2018.

166. Carlos P Cantalapiedra, Ana Hernández-Plaza, Ivica Letunic, Peer Bork, and Jaime Huerta-Cepas. eggNOG-mapper v2: Functional annotation, orthology assignments, and domain prediction at the metagenomic scale. Mol. Biol. Evol., 38(12):5825–5829, December 2021.

167. Hiroshi Tsugawa, Tomas Cajka, Tobias Kind, Yan Ma, Brendan Higgins, Kazutaka Ikeda, Mitsuhiro Kanazawa, Jean VanderGheynst, Oliver Fiehn, and Masanori Arita. MS-DIAL: data-independent MS/MS deconvolution for comprehensive metabolome analysis. Nat. Methods, 12(6):523–526, June 2015.

168. Stuart Berg, Dominik Kutra, Thorben Kroeger, Christoph N Straehle, Bernhard X Kausler, Carsten Haubold, Martin Schiegg, Janez Ales, Thorsten Beier, Markus Rudy, and Others. Ilastik: interactive machine learning for (bio) image analysis. Nat. Methods, 16(12):1226– 1232, 2019.

169. Johannes Schindelin, Ignacio Arganda-Carreras, Erwin Frise, Verena Kaynig, Mark Longair, Tobias Pietzsch, Stephan Preibisch, Curtis Rueden, Stephan Saalfeld, Benjamin Schmid, Jean-Yves Tinevez, Daniel James White, Volker Hartenstein, Kevin Eliceiri, Pavel Tomancak, and Albert Cardona. Fiji: an open-source platform for biological-image analysis. Nat. Methods, 9(7):676–682, June 2012.

170. Nicholas Sofroniew, Talley Lambert, Kira Evans, Juan Nunez-Iglesias, Grzegorz Bokota, Philip Winston, Gonzalo Peña-Castellanos, Kevin Yamauchi, Matthias Bussonnier, Draga Doncila Pop, Ahmet Can Solak, Ziyang Liu, Pam Wadhwa, Alister Burt, Genevieve Buckley, Andrew Sweet, Lukasz Migas, Volker Hilsenstein, Lorenzo Gaifas, Jordão Bragantini, Jaime Rodríguez-Guerra, Hector Muñoz, Jeremy Freeman, Peter Boone, Alan Lowe, Christoph Gohlke, Loic Royer, Andrea Pierré, Hagai Har-Gil, and Abigail McGovern. napari: a multi-dimensional image viewer for python, 2022.

171. M Ashburner, C A Ball, J A Blake, D Botstein, H Butler, J M Cherry, A P Davis, K Dolinski, S S Dwight, J T Eppig, M A Harris, D P Hill, L Issel-Tarver, A Kasarskis, S Lewis, J C Matese, J E Richardson, M Ringwald, G M Rubin, and G Sherlock. Gene ontology: tool for the unification of biology. the gene ontology consortium. Nat. Genet., 25(1):25–29, May 2000.

172. Gene Ontology Consortium, Suzi A Aleksander, James Balhoff, Seth Carbon, J Michael Cherry, Harold J Drabkin, Dustin Ebert, Marc Feuermann, Pascale Gaudet, Nomi L Harris, David P Hill, Raymond Lee, Huaiyu Mi, Sierra Moxon, Christopher J Mungall, Anushya Muruganugan, Tremayne Mushayahama, Paul W Sternberg, Paul D Thomas, Kimberly Van Auken, Jolene Ramsey, Deborah A Siegele, Rex L Chisholm, Petra Fey, Maria Cristina Aspromonte, Maria Victoria Nugnes, Federica Quaglia, Silvio Tosatto, Michelle Giglio, Suvarna Nadendla, Giulia Antonazzo, Helen Attrill, Gil Dos Santos, Steven Marygold, Victor Strelets, Christopher J Tabone, Jim Thurmond, Pinglei Zhou, Saadullah H Ahmed, Praoparn Asanitthong, Diana Luna Buitrago, Meltem N Erdol, Matthew C Gage, Mohamed Ali Kadhum, Kan Yan Chloe Li, Miao Long, Aleksandra Michalak, Angeline Pesala, Armalya Pritazahra, Shirin C C Saverimuttu, Renzhi Su, Kate E Thurlow, Ruth C Lovering, Colin Logie, Snezhana Oliferenko, Judith Blake, Karen Christie, Lori Corbani, Mary E Dolan, Harold J Drabkin, David P Hill, Li Ni, Dmitry Sitnikov, Cynthia Smith, Alayne Cuzick, James Seager, Laurel Cooper, Justin Elser, Pankaj Jaiswal, Parul Gupta, Pankaj Jaiswal, Sushma Naithani, Manuel Lera-Ramirez, Kim Rutherford, Valerie Wood, Jeffrey L De Pons, Melinda R Dwinell, G Thomas Hayman, Mary L Kaldunski, Anne E Kwitek, Stanley J F Laulederkind, Marek A Tutaj, Mahima Vedi, Shur-Jen Wang, Peter D’Eustachio, Lucila Aimo, Kristian Axelsen, Alan Bridge, Nevila Hyka-Nouspikel, Anne Morgat, Suzi A Aleksander, J Michael Cherry, Stacia R Engel, Kalpana Karra, Stuart R Miyasato, Robert S Nash, Marek S Skrzypek, Shuai Weng, Edith D Wong, Erika Bakker, Tanya Z Berardini, Leonore Reiser, Andrea Auchincloss, Kristian Axelsen, Ghislaine Argoud-Puy, Marie-Claude Blatter, Emmanuel Boutet, Lionel Breuza, Alan Bridge, Cristina Casals-Casas, Elisabeth Coudert, Anne Estreicher, Maria Livia Famiglietti, Marc Feuermann, Arnaud Gos, Nadine Gruaz-Gumowski, Chantal Hulo, Nevila Hyka-Nouspikel, Florence Jungo, Philippe Le Mercier, Damien Lieberherr, Patrick Masson, Anne Morgat, Ivo Pedruzzi, Lucille Pourcel, Sylvain Poux, Catherine Rivoire, Shyamala Sundaram, Alex Bateman, Emily Bowler-Barnett, Hema Bye-A-Jee, Paul Denny, Alexandr Ignatchenko, Rizwan Ishtiaq, Antonia Lock, Yvonne Lussi, Michele Magrane, Maria J Martin, Sandra Orchard, Pedro Raposo, Elena Speretta, Nidhi Tyagi, Kate Warner, Rossana Zaru, Alexander D Diehl, Raymond Lee, Juancarlos Chan, Stavros Diamantakis, Daniela Raciti, Magdalena Zarowiecki, Malcolm Fisher, Christina James-Zorn, Virgilio Ponferrada, Aaron Zorn, Sridhar Ramachandran, Leyla Ruzicka, and Monte Westerfield. The gene ontology knowledgebase in 2023. Genetics, 224(1), May 2023.

173. John Jumper, Richard Evans, Alexander Pritzel, Tim Green, Michael Figurnov, Olaf Ronneberger, Kathryn Tunyasuvunakool, Russ Bates, Augustin Žídek, Anna Potapenko, Alex Bridgland, Clemens Meyer, Simon A A Kohl, Andrew J Ballard, Andrew Cowie, Bernardino Romera-Paredes, Stanislav Nikolov, Rishub Jain, Jonas Adler, Trevor Back, Stig Petersen, David Reiman, Ellen Clancy, Michal Zielinski, Martin Steinegger, Michalina Pacholska, Tamas Berghammer, Sebastian Bodenstein, David Silver, Oriol Vinyals, Andrew W Senior, Koray Kavukcuoglu, Pushmeet Kohli, and Demis Hassabis. Highly accurate protein structure prediction with AlphaFold. Nature, 596(7873):583–589, August 2021.

174. Optical coherence tomography, 2015.

175. Jaakko Lehtinen, Jacob Munkberg, Jon Hasselgren, Samuli Laine, Tero Karras, Miika Aittala, and Timo Aila. Noise2Noise: Learning image restoration without clean data. March 2018.

176. Olaf Ronneberger, Philipp Fischer, and Thomas Brox. U-Net: Convolutional networks for biomedical image segmentation. In Medical Image Computing and Computer-Assisted Intervention – MICCAI 2015, pages 234–241. Springer International Publishing, 2015.

177. Xinyang Li, Guoxun Zhang, Jiamin Wu, Yuanlong Zhang, Zhifeng Zhao, Xing Lin, Hui Qiao, Hao Xie, Haoqian Wang, Lu Fang, and Qionghai Dai. Reinforcing neuron extraction and spike inference in calcium imaging using deep self-supervised denoising. Nat. Methods, 18(11):1395–1400, November 2021.

178. Diederik P Kingma and Jimmy Ba. Adam: A method for stochastic optimization. December 2014.

179. Michael Doube, Michał M Kłosowski, Ignacio Arganda-Carreras, Fabrice P Cordelières, Robert P Dougherty, Jonathan S Jackson, Benjamin Schmid, John R Hutchinson, and Sandra J Shefelbine. BoneJ: Free and extensible bone image analysis in ImageJ. Bone, 47(6):1076–1079, December 2010.

180. David Legland, Ignacio Arganda-Carreras, and Philippe Andrey. MorphoLibJ: integrated library and plugins for mathematical morphology with ImageJ. Bioinformatics, 32(22):3532–3534, November 2016.

181. Fabian Ruperti. Ocm segmentation analysis. https://git.embl.de/grp-arendt/spongeprot/-/blob/main/Image%20analysis/OCM/OCM_measurement_correlation.ipynb, 2023. Accessed: 2023-07.

182. Klaske J Schippers and Scott A Nichols. Evidence of signaling and adhesion roles for β-Catenin in the sponge ephydatia muelleri. Mol. Biol. Evol., 35(6):1407–1421, June 2018.

183. Friedrich B M Reinhard, Dirk Eberhard, Thilo Werner, Holger Franken, Dorothee Childs, Carola Doce, Maria Fälth Savitski, Wolfgang Huber, Marcus Bantscheff, Mikhail M Savitski, and Gerard Drewes. Thermal proteome profiling monitors ligand interactions with cellular membrane proteins. Nat. Methods, 12(12):1129–1131, December 2015.

184. Isabelle Becher, Thilo Werner, Carola Doce, Esther A Zaal, Ina Tögel, Crystal A Khan, Anne Rueger, Marcel Muelbaier, Elsa Salzer, Celia R Berkers, Paul F Fitzpatrick, Marcus Bantscheff, and Mikhail M Savitski. Thermal profiling reveals phenylalanine hydroxylase as an off-target of panobinostat. Nat. Chem. Biol., 12(11):908–910, November 2016.

185. Christopher S Hughes, Sophia Foehr, David A Garfield, Eileen E Furlong, Lars M Stein-metz, and Jeroen Krijgsveld. Ultrasensitive proteome analysis using paramagnetic bead technology. Mol. Syst. Biol., 10(10):757, October 2014.

186. G K Smyth. limma: Linear models for microarray data. In Robert Gentleman, Vincent J Carey, Wolfgang Huber, Rafael A Irizarry, and Sandrine Dudoit, editors, Bioinformatics and Computational Biology Solutions Using R and Bioconductor, pages 397–420. Springer New York, New York, NY, 2005.

187. Wolfgang Huber, Anja von Heydebreck, Holger Sültmann, Annemarie Poustka, and Martin Vingron. Variance stabilization applied to microarray data calibration and to the quantification of differential expression. Bioinformatics, 18 Suppl 1:S96–104, 2002.

188. Korbinian Strimmer. fdrtool: a versatile R package for estimating local and tail area-based false discovery rates. Bioinformatics, 24(12):1461–1462, June 2008.

189. Jaime Huerta-Cepas, Damian Szklarczyk, Davide Heller, Ana Hernández-Plaza, Sofia K Forslund, Helen Cook, Daniel R Mende, Ivica Letunic, Thomas Rattei, Lars J Jensen, Christian von Mering, and Peer Bork. eggNOG 5.0: a hierarchical, functionally and phylogenetically annotated orthology resource based on 5090 organisms and 2502 viruses. Nucleic Acids Res., 47(D1):D309–D314, January 2019.

190. Fabian Ruperti. Import proteomics results. https://git.embl.de/grp-arendt/spongeprot/-/blob/main/Data%20analysis/Proteomics/import_proteomic_results.ipynb, 2023. Accessed: 2023-07.

191. Fabian Ruperti. Phosphoproteomics analysis. https://git.embl.de/grp-arendt/spongeprot/-/blob/main/Data%20analysis/Proteomics/phosphorylation_analysis.ipynb, 2023. Accessed: 2023-07.

192. Kazutaka Katoh and Hiroyuki Toh. Parallelization of the MAFFT multiple sequence alignment program. Bioinformatics, 26(15):1899–1900, August 2010.

193. Subha Kalyaanamoorthy, Bui Quang Minh, Thomas K F Wong, Arndt von Haeseler, and Lars S Jermiin. ModelFinder: fast model selection for accurate phylogenetic estimates. Nat. Methods, 14(6):587–589, June 2017.

194. Diep Thi Hoang, Olga Chernomor, Arndt von Haeseler, Bui Quang Minh, and Le Sy Vinh. UFBoot2: Improving the ultrafast bootstrap approximation. Mol. Biol. Evol., 35(2):518–522, February 2018.

195. F Alexander Wolf, Philipp Angerer, and Fabian J Theis. SCANPY: large-scale single-cell gene expression data analysis. Genome Biol., 19(1):15, February 2018.

196. Fabian Ruperti. single-cell rnaseq barplots. https://git.embl.de/grp-arendt/spongeprot/-/blob/main/Data%20analysis/scRNAseq/scRNA_barplots.ipynb, 2023. Accessed: 2023-07.

197. Fabian Ruperti. scrna expression of proteomics results. https://git.embl.de/grp-arendt/spongeprot/-/blob/main/Data%20analysis/scRNAseq/scRNA_proteomic_results.ipynb, 2023. Accessed: 2023-07.

198. Fabian Ruperti. Secretomics dotplots. https://git.embl.de/grp-arendt/spongeprot/-/blob/main/Data%20analysis/Proteomics/suppl-secret-dotplots.ipynb, 2023. Accessed: 2023-07.

199. J Ellis. XXXI. on the nature and formation of sponges: In a letter from john ellis, esquire, f. r. s. to dr. solander, f. r. S. Philos. Trans. R. Soc. Lond., 55(0):280–289, December 1765.

200. N Lieberkühn. Neue beitrage zur anatomie der spongien. Arch. Anat. Physiol.

201. M Pavans De Ceccatty, M Cargouïl, and E Corabœuf. LES RÉACTIONS MOTRICES DE LEPONGE TETHYA LYNCURIUM (lmk.) a QUELQUES STIMULATIONS EXPERIMENTALES. https://hal.sorbonne-universite.fr/hal-02890256/document. Accessed: 2023-7-2

202. C L Prosser, T Nagai, and R A Nystrom. Oscular contractions in sponges. Comp. Biochem. Physiol., 6:69–74, June 1962.

203. L de Vos and G Van de Vyver. Etude de la contraction spontanée chez l’éponge d’eau douce ephydatia fluviatilis cultivée in vitro. Ann. Soc. R. Zool. Belg./Ann. K. Belg. Ver. Dierkd., 111(1-4):21–31, 1981.

204. George Howard Parker. The Reactions of Sponges: With a Consideration of the Origin of the Nervous System. Museum of Comparative Zoölogy at Harvard College, 1910.

205. Norbert Weissenfels. Bau und funktion des süßwasserschwamms ephydatia fluviatilis (porifera). Zoomorphology, 104(5):292–297, October 1984.

